# Multi-axial DNA origami force spectroscopy unlocks conformational dynamics hidden under single-axial tension

**DOI:** 10.1101/2025.06.10.658941

**Authors:** Gde Bimananda Mahardika Wisna, Ayush Saurabh, Deepak Karna, Ranjan Sasmal, Prathamesh Chopade, Steve Pressé, Rizal F. Hariadi

## Abstract

Biomolecules in living cells experience complex multi-directional mechanical forces that regulate their structure, dynamics, and function. However, most single-molecule techniques primarily exert force along a single axis, thereby failing to emulate the mechanical environment of cells. Here, we demonstrate that single-axis force application fundamentally restricts conformational dynamics by kinetically trapping molecules within distinct dynamic classes, preventing interconversion and exploration of the full conformational landscape. We developed MAESTRO (Multi-Axial Entropic Spring Tweezer along a Rigid Origami), a molecular platform that applies up to 9 pN forces from up to 4 directions simultaneously using programmable ssDNA entropic springs anchored to a DNA origami scaffold. We applied MAESTRO to Holliday junctions (HJs), 4-way DNA intermediates that experience multi-directional tension during homologous recombination. Counterintuitively, we discovered >5× slower kinetics of the HJ conformations under multi-axial tension than under tension-free conditions, enabling direct observations of previously hidden HJ conformational intermediates. Most remarkably, we discovered that multi-axial forces unlock conformational dynamics, enabling interconversion between 5 distinct kinetic classes that remain kinetically inaccessible under zero force or single-axial tension. Furthermore, we demonstrated that tension regulates T7 endonuclease I cleavage site selection, directly linking mechanical environments and molecular mechanics to enzymatic function. By overcoming single-axis limitations, MAESTRO opens new frontiers in molecular mechanobiology, revealing how multi-directional cellular force environments are essential for unlocking the full conformational landscape of biomolecules, and that these complex force patterns serve as master regulators of biological function through mechanisms hidden from conventional approaches.

## Introduction

Mechanical forces on the order of pN acting at the single-molecule level have a profound effect on biomolecular function by inducing nanometer-scale structural changes. Mechanical forces arise through either active mechanisms, where biomolecules convert chemical energy into mechanical work, or passive mechanisms, where biomolecules experience pulling and stretching from their surroundings. The strengths of this mechanical work are comparable to the thermal energy scale of k*_B_*T ∼ 4.2 pN.nm, suggesting that multi-axial forces broadly influence biomolecular processes^1^ and yet, their effects remain poorly understood.

To investigate the mechanics at the single-molecule level, a variety of revolutionary force spectroscopy tools have been utilized, including optical ^2^ and magnetic tweezers^3^, atomic force microscopy ^4^, centrifugal force spectroscopy ^5^, DNA-based tension probes ^6–9^, and a DNA nanoscopic force clamp^10,11^, often in tandem with single-molecule fluorescence microscopy.^12^ These approaches have successfully quantified the mechanical energy generated by the rotary motor F1-ATPase under specified loads ^13^, identified the stall force of motor proteins such as myosins and kinesins^14,15^, measured the stalling tension in T7-DNA polymerase^16^, investigated stacking forces among DNA bases ^17^, observed conformational changes of DNA Holliday junctions under tension^18^, analyzed the activation of different mechanosensitive membrane proteins^6–9^, and explored the energetics involved in protein folding and unfolding.^19,20^ These techniques have provided transformative insights into molecular mechanics of single-molecule biophysics, achieving high precision over a diverse range of tensions, as seen with atomic force microscopy and magnetic tweezers, and providing high-throughput capabilities, as demonstrated by magnetic tweezers, centrifugal force spectroscopy, DNA origami-based spectroscopy, and DNA-based tension probes. However, they are limited to applying forces along a single axis due to challenges related to alignment and stability, the complexity of force calibration and measurement, and the difficulties in incorporating multiple tether or anchor points without inducing twisting or buckling. ^21–23^

Multi-directional forces are ubiquitous in cellular mechanobiology, where biomolecules experience simultaneous tension from multiple sources. Focal adhesion proteins like integrin experience forces from both extracellular matrix attachment and intracellular cytoskeletal networks, creating multi-directional stress patterns essential for cellular mechanosensing. ^24,25^ Similarly, chromatin-associated proteins experience simultaneous forces from nucleosome interactions, transcriptional machinery, and DNA repair complexes during processes like homologous recombination. ^26^ Despite the biological relevance, methods to exert multi-axial tension and simultaneously monitor biomolecular functions have yet to be developed, limiting our understanding of how mechanical forces regulate biomolecular function in living cells.

To address the single-axis limitation, we developed the Multi-Axial Entropic Spring Tweezer along a Rigid Origami (MAESTRO), a tension-inducing molecular platform that leverages DNA origami ^27,28^ and entropic springs^11,29^ to apply multi-axial forces to biomolecules. With DNA origami, any rigid 3D shape can be realized through the folding of a long single-stranded DNA (ssDNA) scaffold with hundreds of short ssDNA staple strands via Watson-Crick base-pairing. We choose a 3D DNA origami ring as a rigid platform with circular symmetry ^30^ to pattern several ssDNAs with specific lengths spanning the diameter of the origami structure. These ssDNA strands act as entropic springs, each generates a defined tension in the range of 0 − 9 pN along the spring due to the thermal fluctuations.^11,29^

To investigate the effects of physiological multi-axial forces, we chose the Holliday junction (HJ) as our model system. HJs form as central intermediates during homologous recombination, the primary mechanism for repairing DNA double-strand breaks caused by ionizing radiation or replication errors ^31–36^ (Fig.1(a)). During this repair process, DNA strands undergo chromatin remodeling while remaining bound to histones, creating multi-directional tension environments where each of the junction’s 4 arms experiences independent forces from chromatin compaction, protein binding, and branch migration processes. We hypothesize that these multi-axial forces influence how resolvases bind to HJs and bias cleavage patterns toward crossover or non-crossover repair products, potentially affecting genetic outcomes during DNA repair. Furthermore, HJs conformational flexibility and force sensitivity ^18^ make them ideal targets for investigating how physiologically relevant multi-directional forces regulate biomolecular function. However, whether the conformational dynamics observed under single-axis tension represent the full range of behaviors accessible to these HJs, or whether multi-axial force patterns unlock qualitatively different dynamics, has not been tested.

**Fig. 1.**
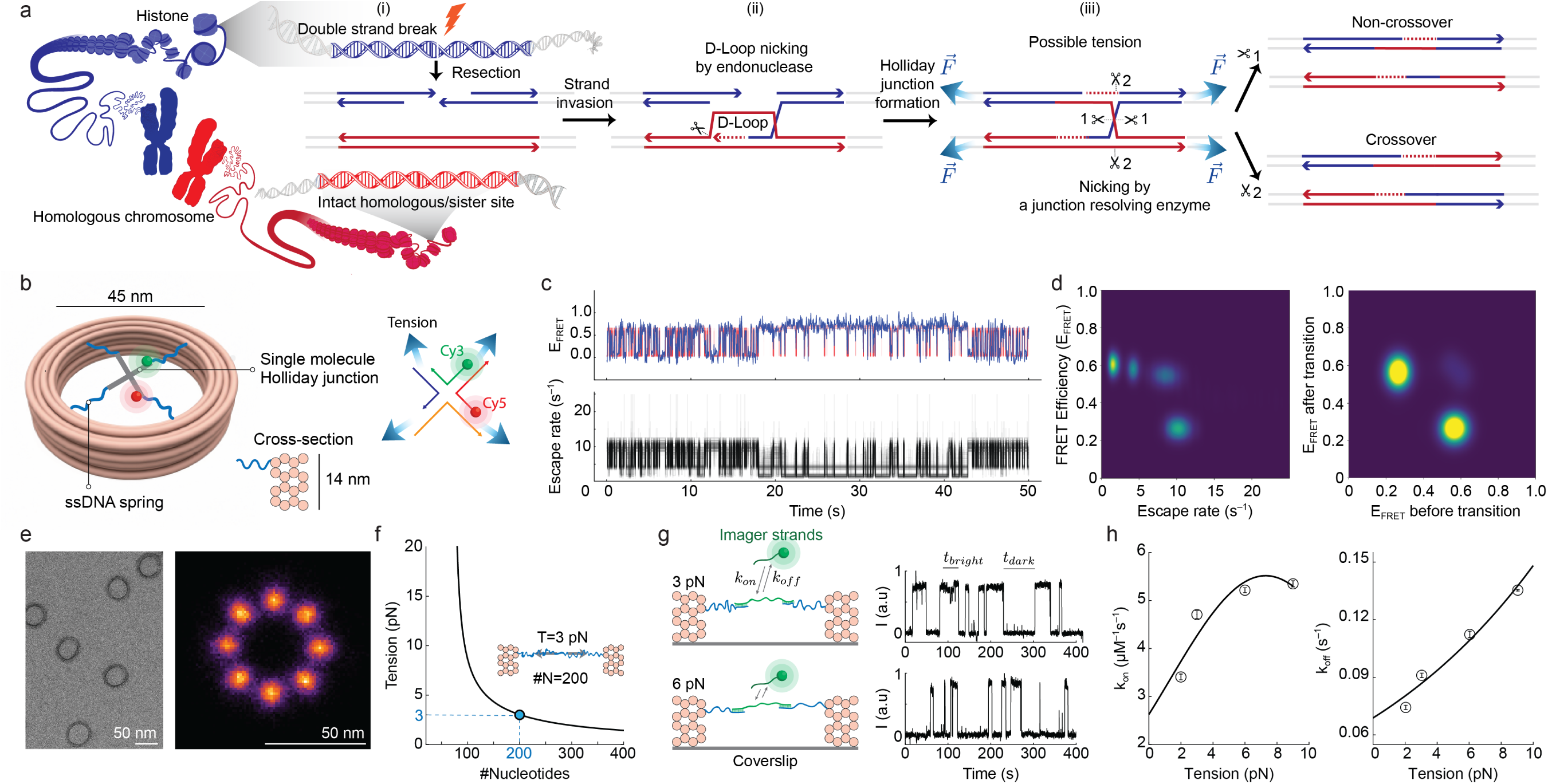
MAESTRO platform enables multi-axial force application, unlocking conformational dynamics. (**a**) Chromatin remodeling enabled formation of Holliday junction during homologous recombination showing double-strand break, D-loop invasion under multi-axial tension. (**b**) MAESTRO schematic showing HJ under controlled multi-axial tension via entropic ssDNA springs. (**c**) Representative smFRET trajectory with Bayesian non-parametric FRET (BNP-FRET) analysis showing HJ dynamics and escape rates (sum of transition rates out of each state). (**d**) Escape rate and transition density heatmaps (N=1000 BNP-FRET samples). (**e**) Structural validation by negative stain-TEM (left) and DNA-PAINT super-resolution imaging (right, N=412). PAINT shows the possible anchoring points for entropic springs. (**f**) Force-extension relationship based on the modified FJC model. (**g**) DNA hybridization kinetics under tension showing bright/dark time analysis. (**h**) Tension-dependent association and dissociation rates with fitted curves. Error bars: SD from bootstrapping.

Our experiments using MAESTRO reveal, HJ dynamics under multi-axial tension. We find these dynamics to be kinetically heterogeneous, with various combinations of fast and slow transition rates for both high and low FRET conformations. Contrary to the common expectation that forces increase reaction rates, we discovered that multi-axial tension dramatically slows the junction’s isomerization kinetics by 5×. This kinetic trapping allowed us to directly observe previously unseen, short-lived intermediate conformations. Furthermore, we discover that multi-axial forces unlock interconversion between five distinct kinetic classes that remain kinetically trapped under simpler force conditions, removing the kinetic constraints that confine molecules under single-axis tension. These findings fundamentally challenge the conventional understanding of how mechanical forces regulate molecular dynamics, revealing a hidden landscape that only emerges under multi-axial tension.

## Results

### MAESTRO recreates a physiological tension environment where biomolecules experience tension from multiple sources, overcoming the single-axis limitation of existing force spectroscopy

We designed MAESTRO to enable systematic investigation of how force directionality affects molecular conformational dynamics. The platform consists of a rigid 16-helix bundle DNA origami scaffold (Fig. 1(b))^30^ with 8 attachment points for single-stranded DNA (ssDNA) entropic springs positioned at 45*^◦^* intervals around the inner sidewall. Target molecules are positioned at the center of the scaffold, and ssDNA springs (40-80 nucleotides) can be attached to different arms of the target to apply forces from multiple directions simultaneously, while maintaining structural rigidity for precise control (Figs. 1(b), S1 and Table S1). HJs exist in dynamic equilibrium between 2 stacked isomers, namely Iso-I and Iso-II, that differ in which pairs of DNA arms stack together, with transitions mediated by transient open conformations in which all 4 arms are unstacked. We investigated the dynamics of HJ (Table S2 for the complete DNA sequences) under various tension configurations using single-molecule Förster Resonance Energy Transfer (smFRET), where the junctions transition between 2 stacked isomers (Iso-I and Iso-II) with distinct FRET efficiencies (E_FRET_) of ∼0.6 and ∼0.25, respectively (Fig. 1(c and d)).

Our smFRET experiments revealed a wide range of kinetic behaviors among the HJs under tension, implying that each smFRET trajectory can have a distinct set of transition rates between the HJ conformations. Consequently, to observe previously unseen dynamics, we avoid fixing the number of states in advance for the entire population when analyzing our smFRET datasets, as is commonly performed when using traditional hidden Markov model (HMM)-based techniques. ^37^ Therefore, we chose to use Bayesian non-parametric FRET (BNP-FRET)^37,38^ to analyze the smFRET trajectories and extract key kinetic parameters. This method implements an infinite-dimensional HMM and uses Markov chain Monte Carlo (MCMC) techniques to generate samples for the number of states, state trajectories, transition rates, and FRET efficiencies. Furthermore, BNP-FRET uses an accurate noise model for our sCMOS camera and incorporates pixel-by-pixel camera calibration maps to optimally recover the signal from the noisy smFRET data. Finally, the BNP-FRET samples can then be collected together to generate heatmaps for escape rates λ (sum of all transition rates out of a given state) as well as transition densities (Fig. 1(c and d)), and estimate the uncertainties.

### The rigidity and addressability of MAESTRO enable precise multi-directional force application

We employed negative-stain transmission electron microscopy (ns-TEM) to validate that MAESTRO retained its programmed rigid and circular structure (Fig. 1(e)). This rigidity is critical for reliably anchoring entropic ssDNA springs and ensuring accurate tension force estimates based on the modified freely jointed chain (FJC) model. ^11,29^ Additionally, we performed DNA-PAINT super-resolution imaging ^39^ to demonstrate the addressability of 8 sites on the inner sidewall of MAESTRO (Tables S1 and S3). Fig. 1(e) shows eight specific attachment sites where ssDNA springs can be precisely anchored to MAESTRO and meet at a target molecule at the center of MAESTRO (Fig. S2). The tension along the ssDNA spring can be tuned by varying the contour length using the modified FJC model (Fig. 1(f); Methods). For example, a 200 nt ssDNA spring that spans the inner diameter of MAESTRO (Fig. 1(f) inset) produces 3 pN. This exquisite positional control provides the necessary foundation for applying well-defined magnitudes and net directions to pN-scale forces.

### Force-dependent DNA hybridization kinetics validates the precision of MAESTRO across the physiologically relevant 0–9 pN range

To quantitatively validate the ability of MAESTRO to exert precise tension on target molecules, we developed an assay grounded in the tension-dependent hybridization kinetics of complementary short single-stranded DNAs. One strand (the docking strand) is held under a defined tension on the MAESTRO, whereas the complementary strand freely diffuses until hybridization under tension occurs (Fig. 1(g), Tables S1 and S4). Kinetic measurements were performed using DNA-PAINT super-resolution imaging. By varying the lengths of the ssDNA springs, we generated predicted tension forces of 2, 3, 6, and 9 pN along the docking strands. The DNA-PAINT under tension assay confirmed that MAESTRO applies precise and predictable pN-scale forces. We observed that both the association (k_on_) and dissociation (k_off_) rates of a short DNA duplex were systematically increased with the applied tension (Figs. 1(h) and S3). These trends, including the asymptotic plateau of k_on_ and the exponential growth of k_off_, are in agreement with previous studies on force-dependent hybridization kinetics (Methods). ^40^ Furthermore, our extrapolated rate at 0 pN force (k_on_(0) = 2.63 µ/M/sec) closely matches the literature value for this specific DNA sequence,^41^ providing independent confirmation of our force calibration.

### HJ dynamics reveal ergodicity breaking through heterogeneity across 5 non-interconverting kinetic classes

To establish baseline conformational dynamics, we first examined Holliday junction behavior under zero force and single-axis tension–the conditions accessible to existing single-molecule force spectroscopy approaches. To achieve a tension-free condition, each HJ was connected to a MAESTRO by a single ssDNA tether (Fig. 2(a)), allowing the observation of intrinsic conformational dynamics under zero force. HJs transition between 2 stacked isomers (Iso-I and Iso-II) through transient open states (Figs. 2(b)), displaying kinetic heterogeneity within the population. We classified individual molecular dynamics into 5 distinct kinetic classes based on the ratio of escape rates (Figs. 2(c–g), S4). These classes ranged from kinetically trapped (Classes I and V, with escape rates <1 s*^−^*^1^) to dynamically balanced (Class III, with comparable escape rates ∼5 s*^−^*^1^), reflecting the complex energy landscape governing the HJ conformational dynamics. Distinct kinetic signatures are evident in both the escape rate distributions and transition density heatmaps, where trapped classes show narrow, single-peaked histograms, whereas dynamic classes exhibit broader, multi-peaked distributions, reflecting more frequent state sampling. Our classification reproduces the kinetic heterogeneity and non-interconverting behavior of surface-immobilized HJs without MAESTRO, validating that MAESTRO preserves the intrinsic HJ dynamics.^42^ Critically, within our 50-second observation window, individual molecules remain kinetically trapped within their initial kinetic class with no observed interconversions between classes. The 50-sec exposure time is >100× longer than the inverse of individual conformational escape rates in the escape rate heatmaps in Fig. 2. The absence of kinetic class interconversion underscores the non-ergodic nature of HJs under 0 pN, where kinetic class interconversion timescales far exceed the observation time, a kinetic constraint that would prove critical for detecting the profound effects of multi-axial forces in Fig. 3.

**Fig. 2.**
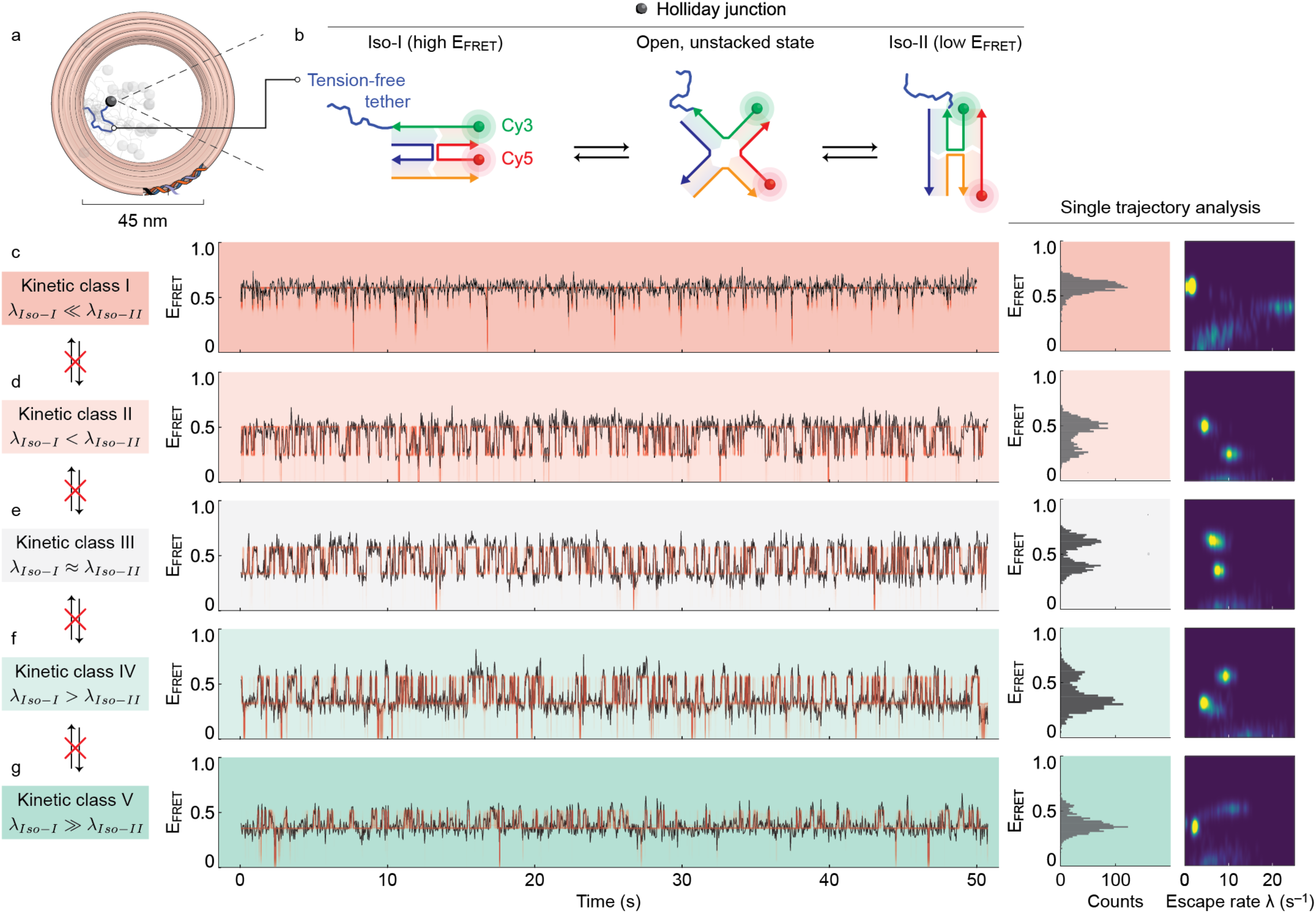
Five kinetic classes of Holliday junction dynamics characterized by FRET efficiency with no interconversion under 0 force. (**a**) MAESTRO platform with HJ tethered (black sphere) to a 0 pN entropic spring (blue). Transparent spheres illustrate HJ spatial dynamics. (**b**) HJ conformational switching between 2 states (Iso-I: E_FRET_∼ 0.6; Iso-II: E_FRET_∼ 0.25) monitored through a FRET pair. (**c-g**) Five kinetic classes with distinct HJ dynamics were observed in different HJ molecules of the same sequence within a single field of view (left panels). (Left) Each row shows a representative 50-sec smFRET trajectory (black) and its corresponding BNP-FRET trajectory (red). (Middle) Histograms of E_FRET_ for different kinetic classes. (Right) Escape rate heatmaps define the characteristic features of each kinetic class. The color scheme for kinetic classes in (**c**–**g**) is consistent with Fig. 3.

**Fig. 3.**
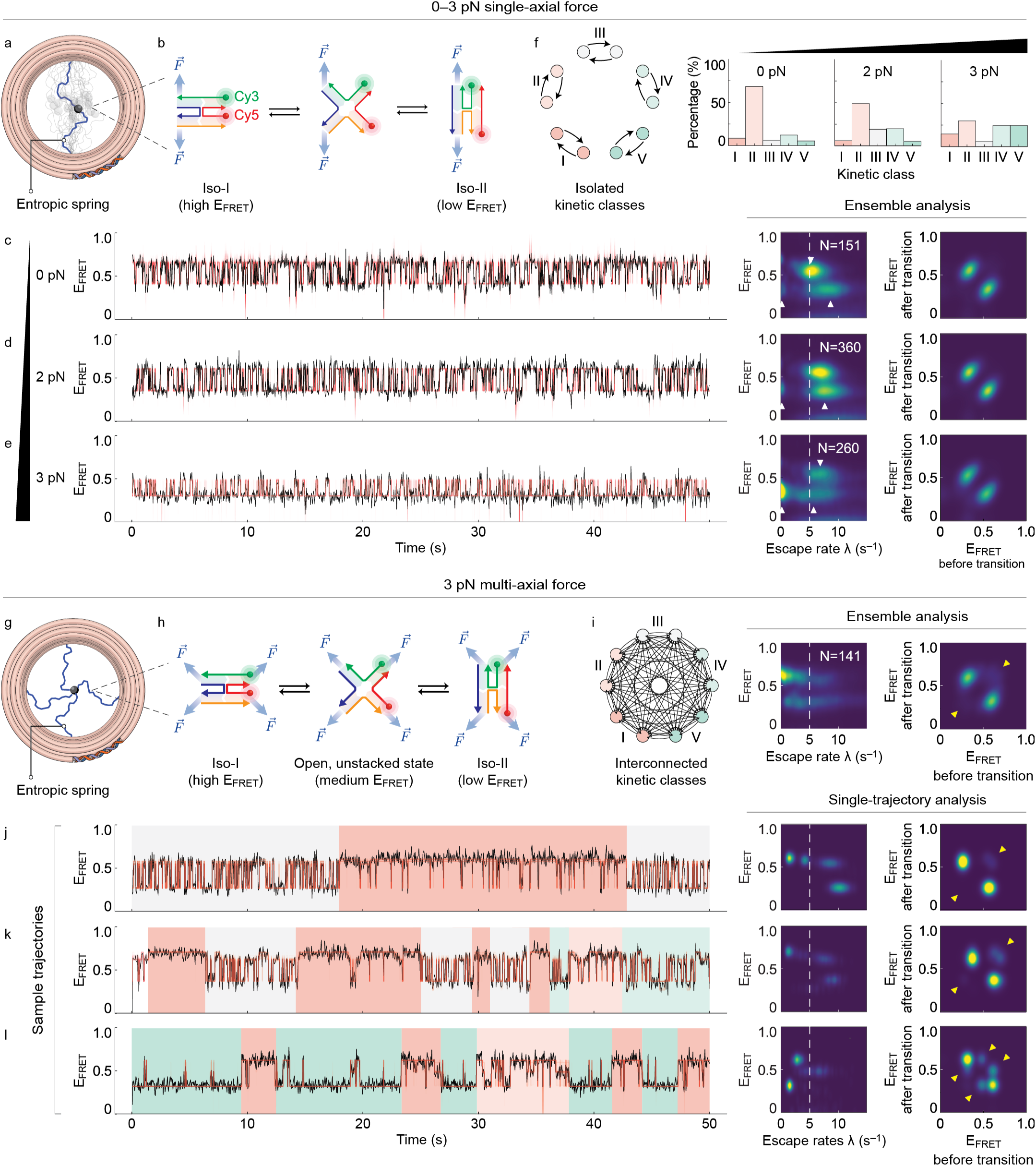
(Previous page) Multi-axial forces unlock kinetic class interconversion. (**a,b**) Single-axial tension setup: Two HJ arms are attached to a pair of entropic springs (blue) spanning the MAESTRO diameter and anchored at diametrically opposite attachment sites. The force direction is indicated by blue arrows. (**c**–**e**) smFRET trajectories (black) and their corresponding BNP-FRET trajectories (red) demonstrate progressive shifts in class populations and dynamics as the tension increases from 0 to 3 pN (N = 151–360 for each condition), with corresponding escape rate heatmaps. (**f**) Ensemble analysis showing the kinetic class population distributions at the denoted applied tensions. The simplified schematic (left) illustrates the 5 distinct kinetic classes, which remain isolated under single-axial tension, representing an isolated conformational landscape. (**g,h**) Multi-axial configuration: Each of the four HJ arms is coupled to its own entropic spring. A four-way tension of 3 pN was applied along the orthogonal axes as the HJ was pulled outward from the center. (**i**) Ensemble heatmaps of escape rates and transition E_FRET_. The simplified transition network (left) illustrates the extensive interconversion between all 5 kinetic classes, representing an unlocked conformational landscape. (**j**–**l**) FRET trajectories (black) and their corresponding BNP-FRET trajectories (red) show dramatic class switching within 50-sec observation windows (N = 141). Colored background regions highlight the periods of different kinetic classes. White dashed line indicates the escape rate of Iso-I under 0 pN tension. White arrows indicate the escape rates of Iso-I or Iso-II which shows the effect of single-axial tension on the escape rates, and yellow arrows highlight previously undetected intermediate states. The color scheme for the kinetic classes in (**f**) and (**i**–ℓ) is consistent with Fig. 2.

### Single-axial forces create a static kinetic landscape that traps HJ conformations along the direction of the applied tension

Single-axial tension acted as a predictable kinetic clamp, stabilizing the HJ conformation aligned with the force vector in agreement with established mechanical models. Tension along the y-axis (Figs. 3(a and b)) favored the Iso-II conformation (Figs. 3(c–e)) and S5). The observed trends are consistent with earlier findings from optical tweezers experiments on HJs under tension ^18^ and DNA origami-based force using entropic springs. ^11^ Together, these findings confirm that MAESTRO effectively applies mechanical tension to HJ molecules, reshaping their dynamical behavior. This stabilization arises from a force-induced tilt in the free-energy landscape. While still obeying the principle of detailed balance, this tilt in the energy landscape increased the escape rate for the conformation orthogonal to the imposed tilt while significantly slowing the escape rate from the conformation aligned to the applied force. Consequently, the force-aligned conformation is effectively trapped. This kinetic trapping was confirmed by a progressive redistribution of kinetic classes under increasing tension, which favored the kinetically trapped Class V (Fig. 3(f)) while depleting more dynamic populations (Class II; Fig. 3(f)). Notably, HJs exist in kinetically isolated subpopulations that do not interconvert on experimentally accessible timescales (Fig. 3(f) left panel). Single-axial force conditions restrict molecular dynamics, confining molecules to narrow kinetic regimes and preventing exploration of the full conformational landscape.

### Multi-axial tension slows HJ kinetics

We further explored the dynamics of HJs under multi-axial tension using 4-way tension (Figs. 3(g and h), S6, and S7). Despite the mechanical complexity of applying tension in four directions simultaneously, the ensemble transition heatmaps still show two major hotspots (Fig. 3(i)), while revealing previously hidden states as dimmer hotspots (yellow arrows; Fig. 3(i–ℓ)). In stark contrast to conventional models predicting that the applied force increases the rate of conformational transitions, multi-axial tension drove a dramatic, >5× slowing of HJ kinetics (Fig. 3(i)). Multi-axial forces reshape the energy landscape in two ways: they increase barriers within conformational states (causing the >5-fold kinetic slowing) while simultaneously lowering barriers between kinetic classes. The latter effect (Fig. 3(j–ℓ)) enables individual molecules to explore a vastly broader range of kinetic behaviors over time (Fig. 3(i) left panel). Analysis of escape rates revealed a high density of values that were >5× slower (Fig. 3(i), middle and right panels) than those observed under the 0 pN condition (white vertical dashed lines; Fig. 3(c) compared with Fig. 3(i–ℓ)). This kinetic slowdown of both λ_Iso-I_ and λ_Iso-II_ are prominently visible in the escape rate heatmaps (Fig. 3(j–ℓ)), leading us to hypothesize the stabilization of intermediate open, unstacked conformations. These conformational states are normally too transient to be observed in the absence of multi-axial tension with our 10–25 ms exposure time. This hypothesized stabilization enables previously inaccessible levels of conformational connectivity.

### Multi-axial forces unlock kinetic class interconversion, thereby restoring quasi-ergodicity by transforming the rugged kinetic landscape into an interconnected network

Remarkably, 4-way tension induced quasi-ergodic sampling by collapsing the static kinetic classes observed under either 0 pN (Figs. 2 and 3(c)) or simpler single-axial tension (Fig. 3(a–e)), enabling many individual molecules to interconvert between the kinetic classes multiple times over a 50-sec observation window (Fig. 3(j–ℓ)). Additional examples of the four-way tension dynamics which show kinetic class interconversion are shown in Fig. S7. While the transition density heatmap confirms that most transitions occur between the two main states, the corresponding escape rate trajectory and heatmap reveal multiple degenerate states with distinct escape rates. The quasi-ergodicity emerged because multi-axial tension transformed the kinetic landscape from a set of discrete, isolated pathways into a dynamic, interconnected network, enabling individual molecules to explore previously inaccessible regions of the conformational space. This is a significant difference over the previous work^42^, which could only induce a single interconversion event by chemically stripping Mg^2+^ from the junction region to reset the HJs and then reintroducing Mg^2+^ ions. In contrast, our platform achieved multiple spontaneous stochastic interconversions in equilibrium buffer conditions containing 12.5 mM Mg^2+^, revealing that multi-axial forces expand access to previously hidden kinetic pathways. This suggests that the underlying conformational landscape is far more complex than that revealed by single-directional perturbations. Three-way tension also induced kinetic class interconversion, albeit less common than under 4-way tension (Fig. S5 and S8), confirming that multi-axial forces generally enable dynamic exploration of the energy landscape.

### Multi-axial tension stabilizes transition states, enabling direct observation of open conformations

The dramatic kinetic slowing observed under multi-axial tension suggests the stabilization of previously undetectable transition states. Using the BNP-FRET framework to analyze the regions surrounding interconversion events, we directly observed intermediate FRET states between the canonical Iso-I and Iso-II conformations (green arrows in Fig. 4). These states represent the open, unstacked conformation that mediates isomer transition, a state that typically persists for less than 80 ms under high-divalent-ion conditions but is extended to lifetimes of up to ∼1 s under multi-axial tension. ^43^ This represents a greater than 12-fold increase in transition state lifetime, enabling the first direct single-molecule characterization of this elusive intermediate under physiologically relevant buffer conditions containing 12.5 mM Mg^2+^. Our confidence in identifying these transition states stems from the robustness of the BNP-FRET approach, which explicitly accounts for camera noise and background fluctuations, allowing us to distinguish genuine intermediate states from the experimental artifacts. ^38,44,45^ Unlike random noise, which persists for only 1–2 time bins (40 ms each), the observed transition states persist significantly longer and occur repeatedly across multiple trajectories, confirming their physical and biological relevance, especially in facilitating the kinetic class interconversion.

**Fig. 4.**
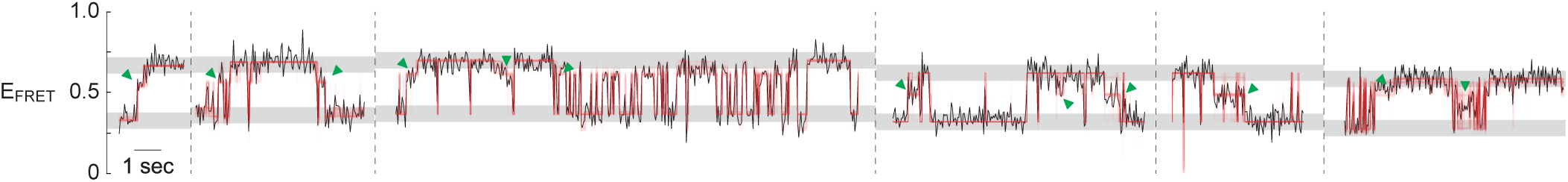
Direct observation of stabilized transition states under 4-way tension. Selected regions of multiple smFRET (black) and their corresponding BNP-FRET trajectories (red) around interconversion events. Green arrows indicate observed transition states stabilized under multi-axial tension, with lifetimes extended from <80 ms to 1 s. Grey-shaded bands with height 0.1 E_FRET_ highlight the ranges corresponding to canonical Iso-I and Iso-II conformations; transition states fall outside these ranges, representing the open, unstacked conformation that mediates kinetic class interconversion.

### Pico Newton forces regulate enzymatic reaction outcomes through control of enzyme-substrate conformations

We tested this enzymatic mechanoregulation hypothesis using T7 endonuclease I, a resolvase that cleaves HJs at specific sites to generate crossover or non-crossover products during DNA repair. T7 endonuclease I is known to bind Holliday junctions in 2 distinct conformations that presumably lead to different cleavage site preferences. Tension at 6 pN induced shifts in enzyme binding conformations (B1 and B2 states) in Mg^2+^ free buffer containing Ca^2+^ (Fig. 5(a and b)). This buffer inhibits cleavage while preserving binding interactions. The observed shift in the distributions of B1 and B2 states demonstrates force-dependent enzyme-substrate interactions. Separately, tension at 6 pN only reduced the enzyme’s cleavage activity by ∼5% compared to tension-free conditions, as quantified by denaturing PAGE analysis of Cy3/Cy5-labeled DNA strands (Fig. 5(c)). Despite the modest reduction in the overall cleavage rate, tension selectively altered cleavage patterns, increasing Cy5-strand cleavage while decreasing Cy3-strand cleavage, demonstrating that forces bias resolution toward specific crossover outcomes rather than simply reducing overall activity. This substantial modulation of enzymatic function directly links the conformational stabilization effects observed in our single-molecule studies to biologically relevant outcomes, demonstrating that single-axial forces can bias DNA repair pathways by mechanically modulating enzyme-substrate interactions. The magnitude of this effect suggests that cellular force environments may play a previously unrecognized role in regulating the balance between crossover and non-crossover repair outcomes during homologous recombination.

**Fig. 5.**
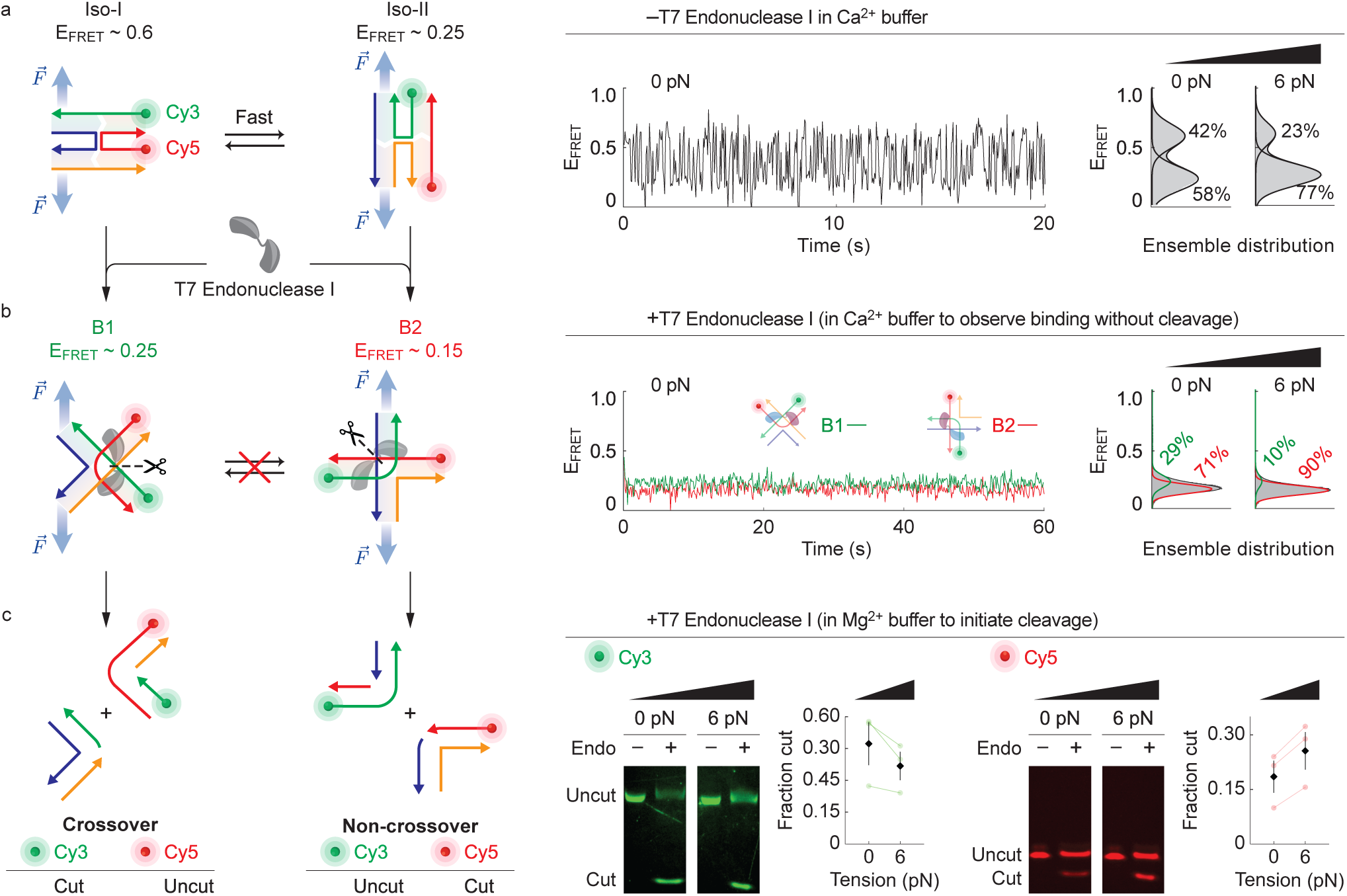
Mechanical tension tunes Holliday junction resolution into crossover and non-crossover products under T7 Endonuclease I. (**a**) Baseline HJ dynamics under single-axial tension in Ca^2+^ buffer. Schematics show HJ isomers (Iso-I and Iso-II) under an applied force (F⃗) with FRET pair labeling (Cy3/Cy5, left). Representative smFRET trace indicates conformational dynamics in Ca^2+^ buffer without enzyme (middle). Ensemble FRET distributions at 0 pN and 6 pN tension show population shifts (N >140, right). (**b**) Enzyme-binding conformations under tension in Ca^2+^ buffer (binding without cleavage). Schematics of T7 endonuclease I binding to HJ under force, showing two distinct bound states (B1: E_FRET_ ∼ 0.25; B2: E_FRET_ ∼ 0.15, left). Representative smFRET trace in Ca^2+^ buffer with the enzyme, revealing stable binding without cleavage (middle). Ensemble distributions demonstrate tension-dependent binding preferences (right). (**c**) Tension-directed resolution outcomes in Mg^2+^ buffer (enabling cleavage). Schematics show two possible cleavage pathways leading to crossover (Cy3 cut, Cy5 intact) or non-crossover (Cy3 intact, Cy5 cut) products (left). Denaturing PAGE analysis and quantification shows that 6 pN tension significantly altered the crossover/non-crossover ratio compared to 0 pN, demonstrating mechanical control of genetic recombination outcomes (N=3 independent experiments; error bars: SD, right).

## Discussion

MAESTRO represents a significant advancement in single-molecule force spectroscopy by enabling programmable multi-axial tension application through a rigid DNA origami scaffold. In contrast to conventional techniques, which are limited to single-axis forces, MAESTRO uses ssDNA entropic springs anchored to a symmetric ring structure to generate physiologically relevant multi-directional force environments spanning 0–9 pN from up to four directions. This capability revealed previously undetectable HJ dynamics, including 5-fold kinetic slowing, stabilization of intermediate states with lifetimes up to ∼1 s, and kinetic class interconversion within individual molecules, which remained inaccessible to single-axis methods despite their proven value in mechanobiology research. These discoveries challenge the conventional understanding that increased mechanical force universally accelerates molecular dynamics. Furthermore, our work elucidates that the complexity of force environments is essential for unlocking molecular conformation dynamics.

Our results point to a detailed mechanism where multi-axial forces reshape the energy landscape topology. We discovered that 4-way tension slows HJ kinetics by >5-fold while simultaneously enabling quasi-ergodicity via kinetic class interconversion. This counterintuitive finding reveals the mechanism of unlocking: multi-axial forces lower the energy barriers *between* kinetic classes while increasing the barriers *within* each state. The net effect is slower local dynamics but enhanced global exploration. This landscape reshaping is mediated by the stabilization of a key intermediate. Under 4-way tension, the open conformation (E_FRET_∼0.45) that mediates isomer transitions persists for up to 1 second, a dramatic extension from its <80 ms lifetime under tension-free conditions. This demonstrates how multi-directional forces can transform transient intermediates into stable, observable states. We propose this stabilization arises because isotropic stress from multi-axial forces favors the symmetric, open state over the asymmetric stacked conformations preferred under unidirectional tension. This is analogous to how divalent cations modulate junction dynamics^46,47^; multi-directional tension may facilitate the release of trapped Mg^2+^ ions that otherwise lock the junction in specific states, creating a more complex energy landscape with new transition pathways.

Our findings have important implications for interpreting kinetic heterogeneity observed in previous single-molecule studies. Our results suggest that the kinetic heterogeneity observed under single-axis tension may reflect the kinetic isolation of subpopulations that would interconvert under more complex force conditions. This does not invalidate previous findings; the kinetic classes identified under single-axis tension represent true features of the energy landscape. However, it suggests that single-axis studies may be observing molecules in mechanically constrained states that do not reflect their behavior in vivo. Under single-axis tension, molecular populations appear non-ergodic, but this may be a consequence of the simplified force geometry. Under multi-axial forces that better approximate cellular conditions, the same system exhibits enhanced ergodic sampling. We emphasize that single-axis force spectroscopy remains an essential tool; our work establishes multi-axial approaches as a critical complement for investigating how force patterns regulate function in complex cellular environments.

This conformational control mechanism extends beyond HJs to establish a direct link between the cellular force environment and enzymatic regulation. The force-dependent nicking of T7 endonuclease I demonstrates how multi-axial forces can bias DNA repair pathways through substrate conformational selection. During homologous recombination, proteins such as RecA/Rad51, and RuvAB exert simultaneous forces on different arms of the HJ. The coordinated action of these proteins likely generates multi-axial force patterns that *liberate* conformational dynamics essential for proper junction processing. This principle of force-dependent conformational control has broad implications for mechanobiology, including tension-dependent integrin activation ^24^ and force-regulated polymerase activity ^16^. The quasi-ergodicity *liberated* by multi-axial forces provides a fundamental mechanism for regulating cellular functions, where complex force networks could serve as molecular switches that control whether biomolecules remain trapped in specific functional states or can dynamically sample their full conformational space.

Building on these mechanistic insights, MAESTRO enables the investigation of biomolecular systems that experience complex force environments in their native cellular contexts while acknowledging current technical limitations. The platform is immediately applicable to mechanosensitive proteins, including ion channels that experience membrane tension from multiple directions, integrins that simultaneously experience forces from the extracellular matrix and cytoskeletal networks, and intrinsically disordered proteins that may adopt different conformations under multi-directional stress. Current constraints include the 0–9 pN force range, which may miss higher-force processes, and ms temporal resolution, which limits the detection of faster dynamics. Near-term technical developments should focus on extending the force range through shorter entropic springs (targeting 10–50 pN), improving temporal resolution through single-photon detection systems (targeting µs timescales), and developing asymmetric force patterns to recreate specific cellular stress environments. These enhancements will establish MAESTRO as an essential tool for understanding how physiological multi-axial forces regulate cellular function, while the current platform already provides unprecedented access to multi-directional force effects that have remained hidden from conventional single-molecule approaches.

## Acknowledgements

The authors gratefully acknowledge Chenxiang Lin for providing the Cadnano file of the NuPOD ring, for valuable scientific discussions, and for suggesting the T7 endonuclease I enzymatic assays. We acknowledge the use of facilities within the Eyring Materials Center and computational resources from the Phoenix and Sol supercomputers at Arizona State University.

## Author Contributions

G.B.M.W. and R.F.H. conceived the idea for the study. G.B.M.W., R.S., D.K., and P.C. assisted with the experiments. G.B.M.W., A.S., and R.F.H. performed the analyses. G.B.M.W., A.S., D.K., P.C., S.P., and R.F.H. assisted with manuscript editing. All authors discussed the results and commented on the manuscript.

## Correspondence

All correspondence and requests for materials should be addressed to G.B.M. Wisna and R.F. Hariadi.

## Materials and methods

### Materials

Unmodified and biotinylated DNA staple strands for DNA origami MAESTRO were purchased from Integrated DNA Technologies (IDT Coralville, IA, USA). Scaffold strands (p8064) were sourced from Bayou Biolabs (Metairie, LA, USA). Amine-modified DNA strands used as imager strands were also obtained from IDT. Cy3B-NHS ester fluorophores (PA63101) were purchased from GE Healthcare (Chicago IL, USA). T7 endonuclease I (M0302L) was obtained from New England Biolabs (Ipswich, MA). BSA-biotin (A8549) and streptavidin (S4762) were purchased from Sigma-Aldrich (St. Louis, MO). All general chemicals were supplied by Sigma-Aldrich, unless otherwise noted. Glass coverslips (48466-205, 24×60 mm, #1.5) and microscope slides (16004-430) were obtained from VWR (Radnor, PA). Kapton tape (PPTDE-2, 0.5 mm thickness) was acquired from Bertech (Skokie, IL, USA). Amicon Ultra-0.5 centrifugal filter units (100 kDa MWCO, UFC510024) were purchased from Millipore Sigma. Carbon-coated TEM grids (01814-F) were purchased from Ted Pella (Redding, CA, USA). Uranyl acetate and all buffer components were obtained from Sigma-Aldrich.

### DNA-PAINT Imager Strands

Imager strands (TATGTAGATC/3’ Cy3B/) were prepared by conjugating amine-modified DNA oligonucleotides with Cy3B fluorophores via NHS ester coupling. The conjugated products were subsequently purified using high-performance liquid chromatography (HPLC).

### Tension Force Estimation Using the Modified Freely-Jointed Chain (FJC) Model

We estimated the tension force generated along the ssDNA springs using a modified freely-jointed chain (FJC) polymer model (Eq. 1).^29^

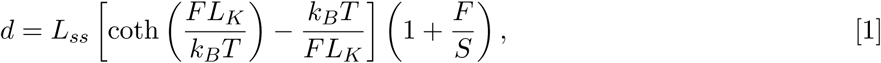

where *d* is the end-to-end distance of the ssDNA spring (45 nm for MAESTRO), L*_ss_* = N · ℓ*_nt_* is the total contour length with N nucleotides and ℓ*_nt_* = 0.6–0.7 nm/nucleotide, F is the applied tension, L*_K_* = 1.5 nm is the Kuhn length, and S is the elastic modulus. ^29^

To achieve the desired tension values, the selected staple strands were extended with uniquely designed ssDNA spring sequences at defined positions (Table S1). Spring sequences were designed using NUPACK^48^ to minimize the unintended intra- and inter-molecular interactions between springs, ensuring consistency with the FJC model’s assumptions of non-interacting polymer chains (Fig. S9).

### MAESTRO Folding and Purification

DNA origami MAESTRO structures were assembled by mixing p8064 scaffold strands (20 nM final concentration) with staple strands, biotinylated strands, and entropic spring strands (each at 200 nM, 10-fold molar excess) in folding buffer (1× TAE, 12.5 mM MgCl_2_, pH 8.0). Total reaction volume was 50 µL (Tables S1 and S4). The annealing protocol consisted of: (1) initial denaturation at 80 *^◦^*C for 5 min, (2) gradual cooling to 4 *^◦^*C at 0.31 *^◦^*C/min (3 min 12 s per degree), and (3) storage at 4 *^◦^*C until use. Folded structures were purified by 5 rounds of centrifugal filtration (Amicon Ultra-0.5, 100 kDa MWCO) at 14,000×g for 10 min each, with buffer exchange to storage buffer (1× TAE, 12.5 mM MgCl_2_).

### Negative-Stain Transmission Electron Microscopy of MAESTRO

Amicon-purified MAESTRO samples were diluted to a final concentration of 2 nM in 1× TAE buffer supplemented with 12.5 mM MgCl_2_. Carbon-coated TEM grids (Ted Pella 01814-F) were glow-discharged for 30–60 s prior to sample application using a PELCO easiGlow glow discharge cleaning system. A 10 µL aliquot of the diluted sample was applied to the grid and incubated for 1–2 min at room temperature. Excess liquid was gently wicked off using a Whatman filter paper without allowing the grid to dry completely. For negative staining, 10 µL of 2% (w/v) aqueous uranyl acetate solution containing 25 mM NaOH was applied and immediately wicked away. A second 10 µL aliquot of the same staining solution was added and incubated on the grid for 30–60 s before removal. The grids were then completely air-dried before imaging. Negatively stained samples were imaged using a Talos L120C G2 transmission electron microscope (Thermo Fisher Scientific) operated at 120 kV with a magnification of 57,000×. Images were acquired using a Ceta 16M camera, and pixel size calibration was performed using a standard calibration grid.

### Holliday Junction (HJ) Formation

HJ samples were prepared by mixing four single-stranded DNA (ssDNA) oligonucleotides at equimolar concentrations. Specific strand extensions were included in the selected strands depending on the desired tension configuration (two-way, three-way, or four-way; see Table S2). The strands were diluted in buffer to a final concentration of 1× TAE supplemented with 12.5 mM MgCl_2_. The mixture was annealed by heating to 80 *^◦^*C for 5 mins, followed by gradual cooling to 4 *^◦^*C at a rate of 3 mins and 12 secs per degree Celsius.

### DNA-PAINT Super-Resolution Imaging for Short Oligo Tension Studies and Data Processing

MAESTRO structures bearing different ssDNA entropic springs corresponding to defined tension values were folded and purified as described above. These were then mixed with short docking oligos at equimolar concentrations and incubated at room temperature for ≥3 h to allow hybridization of the docking strands (Table S1) to the ssDNA springs (spring-1 and spring-2).

DNA-PAINT super-resolution imaging was performed following established protocols. ^39^ MAESTRO structures bearing 8 docking sites on their inner sidewalls (Fig. 1(e), right panel and Fig. S2) or incorporating short oligos under defined mechanical tension (Fig. 1(g), left panel; Tables S3 and S4) were immobilized onto coverslips coated with BSA-biotin and streptavidin. Coverslips were assembled into flow chambers using standard glass microscope slides and double-sided Kapton tape (0.5 mm thickness). For surface preparation, BSA-biotin and streptavidin were diluted to final concentrations of 1 mg/mL and 0.5 mg/mL, respectively, in Buffer A+ (10 mM Tris-HCl, 100 mM NaCl, 0.05% (v/v) Tween 20, pH 8.0). MAESTRO samples were diluted to 2 nM in Buffer B+ (5 mM Tris-HCl, 12.5 mM MgCl_2_, 1 mM EDTA, 0.05% (v/v) Tween 20, pH 8.0). The final imaging buffer contained an oxygen scavenging system comprising 1.25× PCA, 1× PCD, and 1× Trolox, along with 5 nM imager strands (TATGTAGATC/3’ Cy3B/). Flow chambers were sealed with epoxy before imaging.

Imaging was conducted using an Oxford Nanoimager (ONI) Benchtop Nanoimager S Mark II equipped with total internal reflection fluorescence (TIRF) microscopy. Cy3B fluorophores were excited with a 532 nm laser at power densities between 800 and 1250 W/cm^2^, and emission was collected using a 549–623 nm bandpass filter. A 100× oil-immersion Olympus objective (NA 1.49) was used, and images were captured with a Hamamatsu ORCA-Flash4.0 V3 sCMOS camera at exposure times of 200 ms per frame for MAESTRO constructs with eight docking sites and 100 ms for short oligos under tension studies. The system features z-lock autofocus and a piezo-controlled stage for precise positioning. All data were collected at room temperature (∼23 *^◦^*C).

DNA-PAINT movies of MAESTRO constructs with eight docking sites were analyzed using Picasso software ^39^ for localization, rendering, and generation of averaged images from 412 MAESTRO molecules. Binding and unbinding durations were extracted from image stacks using a custom Python script. To estimate λ_on_ and λ_off_, the distributions of the bright and dark times were fitted using the cumulative distribution function (CDF) of an exponential distribution (Fig. S3):

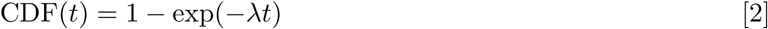

To analyze the force dependence of the oligo binding kinetics, we fitted the experimental data using the models from Hart et al. ^40^, given by:

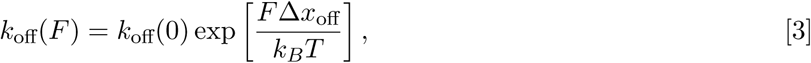

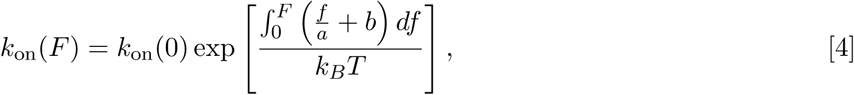

where k*_B_* is the Boltzmann constant, and T is the temperature. Parameters k_off_(0) and k_on_(0) represent the dissociation and association rates at F = 0 pN, respectively. Δx_off_ denotes the extension difference between the transition and bound states. Parameters a and b represent the differences in the stiffness and relaxed extension between the unbound and transition states. Fitted parameter values are summarized in Table S5.

### smFRET of HJs Under Tension

MAESTRO constructs incorporating various single-stranded DNA (ssDNA) entropic springs, which were designed to generate specific tension magnitudes and configurations, were folded and purified. Corresponding HJ samples for each tension configuration were prepared separately. MAESTRO and HJ samples were then mixed in equal volumes to achieve a final concentration of 5 nM each and incubated at room temperature on a shaker (400 rpm) for at least 4 hours to allow hybridization between the HJ extensions and the ssDNA springs.

smFRET samples were prepared on a coverslip assembled into a flow chamber using a glass microscope slide and double-sided Kapton tape. Surface functionalization was performed by sequentially introducing BSA-biotin and streptavidin, diluted to final concentrations of 1 mg/mL and 0.5 mg/mL, respectively, in Buffer A+ (10 mM Tris-HCl, 100 mM NaCl, 0.05% (v/v) Tween 20, pH 8.0). The MAESTRO–HJ complex was diluted to 2 nM in Buffer B+ (5 mM Tris-HCl, 12.5 mM MgCl_2_, 1 mM EDTA, 0.05% (v/v) Tween 20, pH 8.0) and immobilized onto the functionalized coverslip.

Sample introduction into the flow chamber followed this sequence: 20 µL of BSA-biotin, a Buffer A+ wash, streptavidin, a second Buffer A+ wash followed by Buffer B+, the MAESTRO–HJ complex, a final Buffer B+ wash, and then the imaging buffer. Each step was incubated for 2 mins. The imaging buffer consisted of Buffer B+ supplemented with an oxygen-scavenging system containing 1.25× PCA, 1× PCD, and 1× Trolox. The flow chamber was sealed with epoxy prior to imaging.

smFRET imaging was performed using an Oxford Nanoimager (ONI) Benchtop Nanoimager S Mark II equipped with total internal reflection fluorescence (TIRF) optics. Alternating laser excitation (ALEX) was used to sequentially excite green (532 nm) and red (638 nm) lasers, which were synchronized with camera acquisition. Each excitation had a 20 ms exposure time, resulting in an effective binning time of 40 ms. A total of 5,000 frames were acquired for each field of view. Imaging was conducted using a 100× oil-immersion Olympus objective (NA 1.49), and fluorescence was captured with a Hamamatsu ORCA-Flash4.0 V3 sCMOS camera. Green laser power densities ranged from 115–340 W/cm^2^, while red laser power ranged from 75–240 W/cm^2^. The microscope was equipped with a z-lock autofocus system and piezo-controlled stage for high-precision imaging. All data were collected at room temperature (∼23 *^◦^*C).

### smFRET of HJs Under Tension with T7 endonuclease I

The preparation of MAESTRO–HJ samples followed the same protocol described in the smFRET of HJs Under Tension section, with one key modification: all buffers were prepared using 1× TAE containing 50 mM NaCl and 12.5 mM CaCl_2_ instead of MgCl_2_ to prevent cleavage of HJs by T7 endonuclease I.^33^ A final concentration of 2 nM MAESTRO–HJ complex was mixed with T7 endonuclease I (New England Biolabs, catalog no. M0302L) at a final amount of 50 units (22.5 ng) in a 20 µL total reaction volume, yielding an enzyme concentration of approximately 18.7 nM. The mixture was incubated at room temperature for at least 10 mins to allow the enzyme to bind to the HJs prior to sample introduction into the flow chamber.

Flow chamber preparation and flow steps followed the same procedure as described in the smFRET of HJs Under Tension section, with buffer B+ replaced with C+ containing 10 mM Tris-HCl, 50 mM NaCl, 12.5 mM CaCl_2_, 0.05% (v/v) Tween 20, and containing buffers to prevent the cleavage.

The smFRET imaging protocol was identical to the standard method, with a slight modification of the camera exposure time. For control experiments without T7 endonuclease I (Fig. 5b, top panel), a 20 ms exposure per frame was used. For samples containing T7 endonuclease I, the exposure time was increased to 60 ms (Fig. 5b, bottom panel) to improve the signal-to-noise ratio and enhance the resolution between the B1 and B2 smFRET states. All data were collected at room temperature (∼23 *^◦^*C).

### HJ Resolution by T7 endonuclease I Assessed via Denaturing PAGE Gel

To investigate the cleavage activity of T7 endonuclease I on HJs under defined tension conditions, we performed an electrophoretic mobility shift assay using denaturing PAGE. A 12% polyacrylamide gel was prepared in the presence of 8 M urea. MAESTRO DNA origami constructs were prepared to apply two different levels of tension to the HJs: 0 pN (tension-free control) and 6 pN. HJs without MAESTRO were included as an additional control. All samples were tested in both the presence and absence of T7 endonuclease I.

For the digestion assay, approximately 100 nM of each DNA sample was pre-incubated with 50 Units of T7 endonuclease I in reaction buffer (10 mM Tris-HCl, 50 mM NaCl, 12.5 mM CaCl_2_, pH 7.9) at room temperature for 15 min to allow efficient enzyme binding. The cleavage reaction was initiated by incubation at 30 *^◦^*C for 40 min. The reaction was quenched by the addition of 0.5 M EDTA (final concentration 50 mM).

To denature the cleaved DNA fragments, equal volumes of formamide loading buffer were added to each sample, followed by heating at 90*^◦^*C for 10 min. The denatured samples were loaded onto the gel and electrophoresed at 150 V for 1 h in 1× TBE buffer. After electrophoresis, the gel was stained with SYBR Gold nucleic acid gel stain and imaged using an Azure 300 gel documentation system (Azure Biosystems). Band intensities were analyzed using a custom Mathematica script to quantify the ratio of cleaved DNA fragments to total DNA fragments. Cleavage efficiency was calculated as the percentage of substrate converted to product bands.

### BNP-FRET Analysis

We employed Bayesian non-parametric FRET (BNP-FRET) analysis^37,38^ to extract kinetic parameters from smFRET trajectories with an unprecedented accuracy and statistical rigor. This advanced method implements an infinite-dimensional hidden Markov model (HMM) that automatically determines the optimal number of conformational states without prior assumptions, while using Markov chain Monte Carlo (MCMC) techniques to generate comprehensive statistical samples for state trajectories, transition rates, and E_FRET_s. The approach incorporates a sophisticated noise model specifically calibrated for our sCMOS camera system, including pixel-by-pixel calibration maps that maximize the signal recovery from noisy single-molecule data. The resulting MCMC samples enable the generation of detailed heatmaps for escape rates λ (representing the sum of all transition rates departing from a given state) and transition density plots (Fig. 1(c and d)), while providing robust uncertainty estimates for all the extracted parameters.

For each experimental dataset, we analyzed a minimum of three fields of view, with each field containing at least 2,500 frames to ensure the statistical robustness of our findings. Single-molecule FRET traces were extracted from more than 100 individual molecules per condition, providing sufficient data for reliable statistical inferences. To maintain data quality, we applied stringent filtering criteria, including ALEX stoichiometry filtering^49^ to confirm single-molecule behavior, assessment of total donor and acceptor fluorescence stability during donor excitation to identify photobleaching or blinking artifacts, and a minimum requirement of at least one observable FRET state transition per trace to ensure dynamic behavior. Camera calibration involved acquiring 2,500 dark noise frames under identical experimental conditions but without laser excitation or sample presence. The local background fluorescence was systematically subtracted during trace extraction. The filtered and calibrated single-molecule traces were subsequently processed through the BNP-FRET computational pipeline on Arizona State University’s high-performance computing clusters Sol and Phoenix, with detailed methodology described in Saurabh et al. ^38^

### Code Availability

The BNP-FRET analysis pipeline is available at: https://github.com/LabPresse/ BNP-FRET-Binned.

## Supporting Information

**Table S1.**
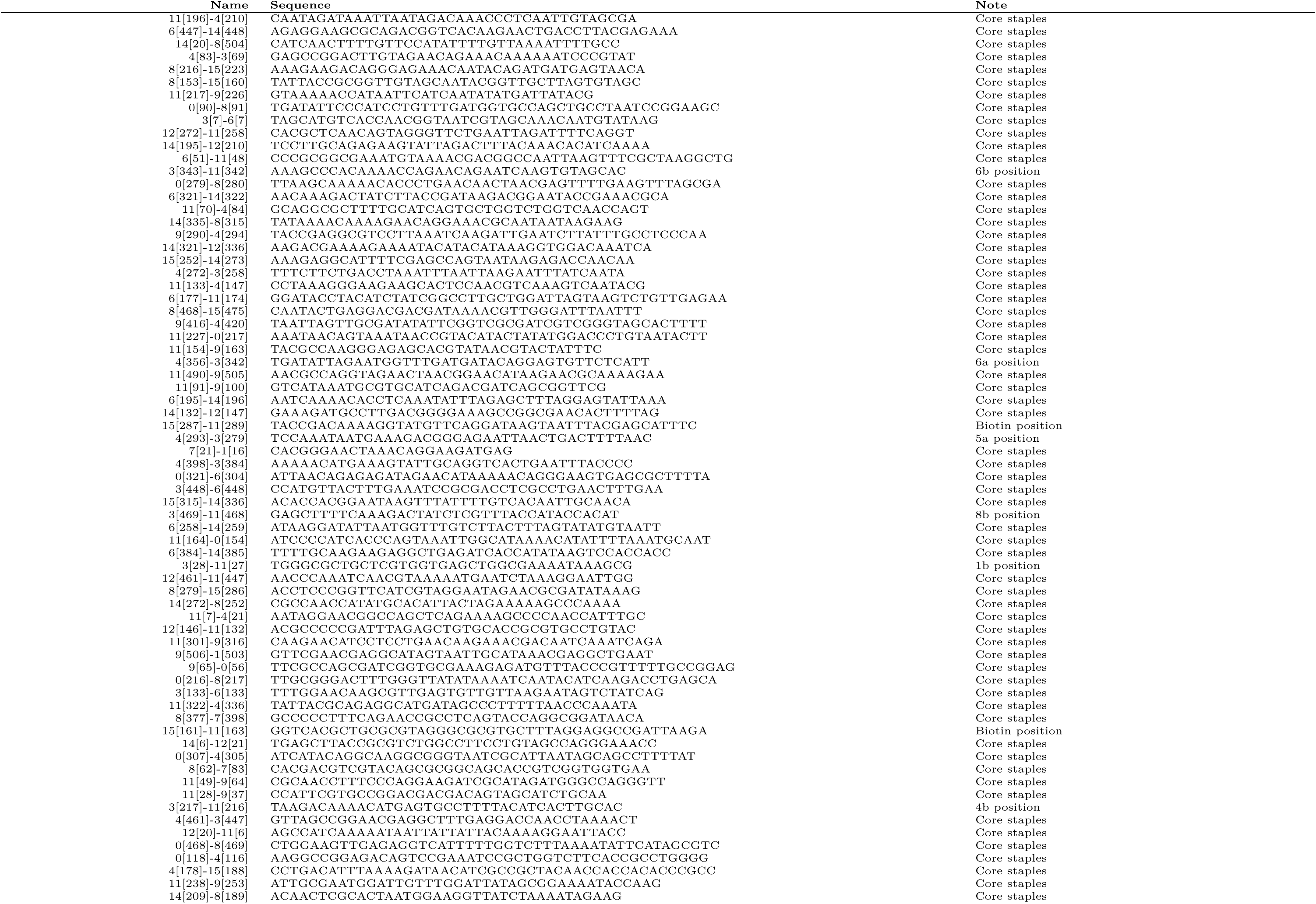
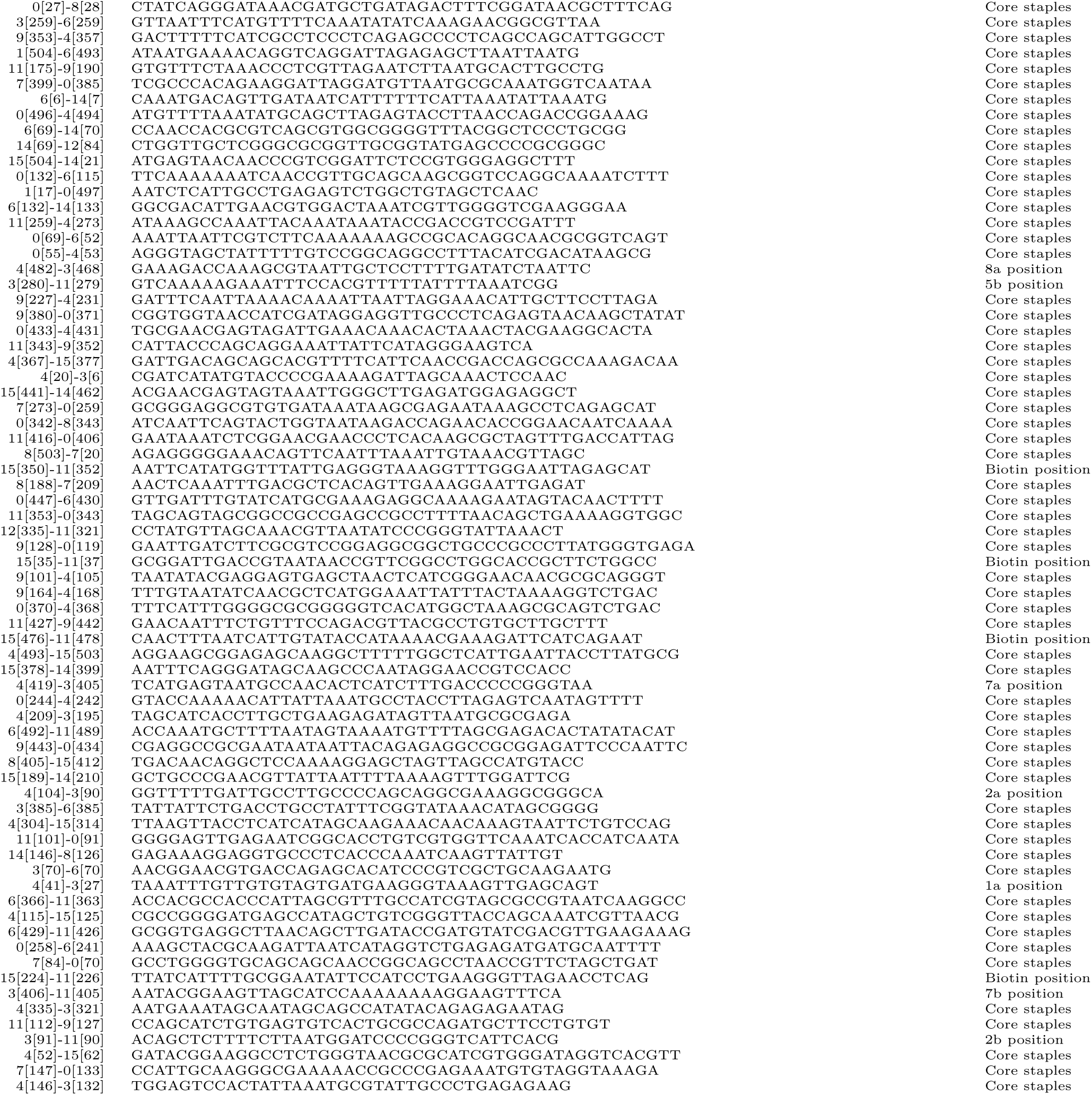
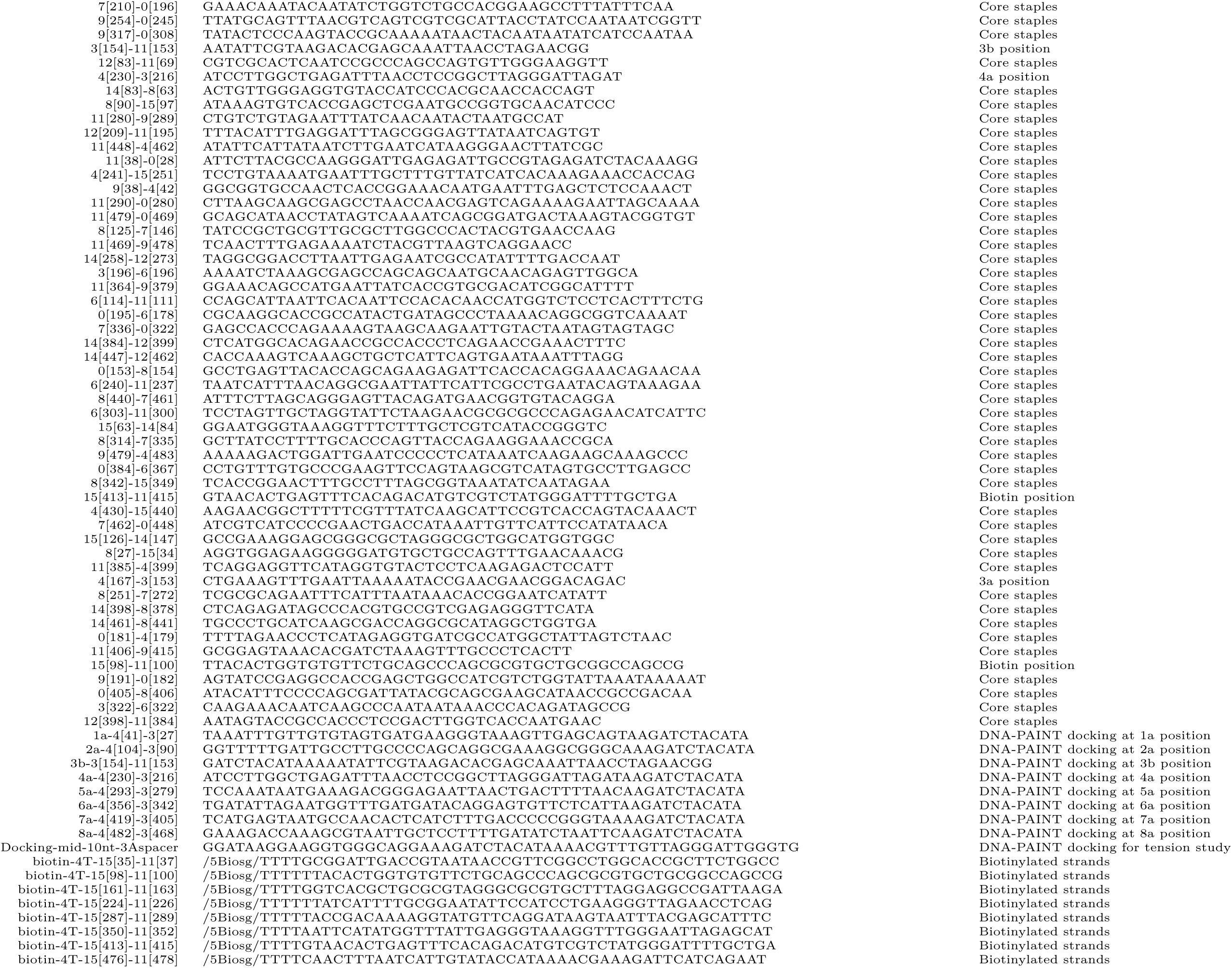
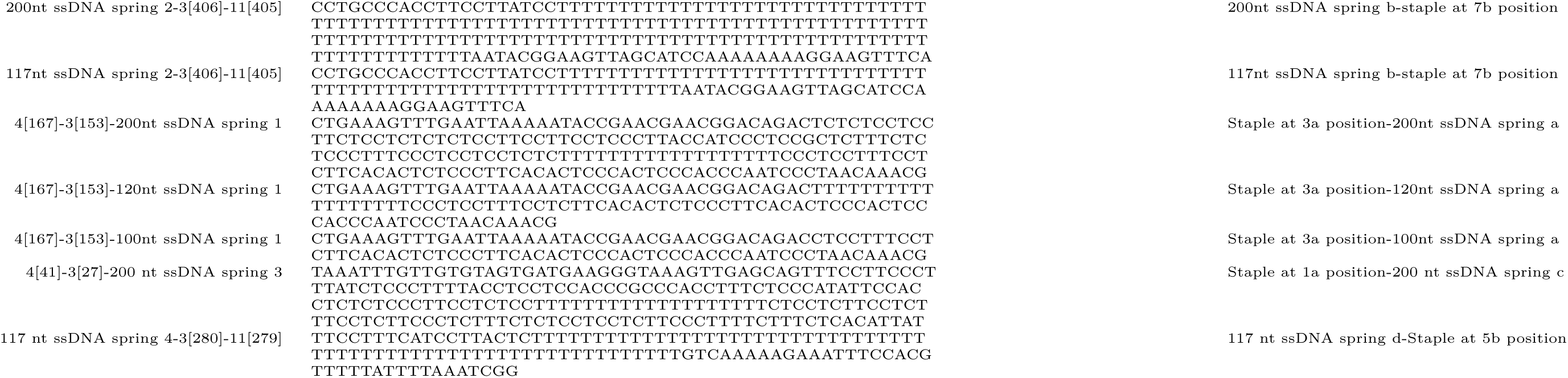
MAESTRO structure, DNA-PAINT, and ssDNA entropic spring strands. The scaffold is p8064. Core staples are all the strands that form the structure. Biotinylated staples were modified with biotin to immobilize MAESTRO on coverslip surfaces through a BSA-biotin-streptavidin-biotin-DNA origami arrangement. DNA-PAINT docking strands were used to acquire super-resolution DNA-PAINT images of MAESTRO (Fig. 1e, right panel). ssDNA entropic spring strands are selected to form MAESTRO with specific tension force configurations. Positions 1 (a,b) to 8 (a,b) correspond to specific sites on the inner sidewall of MAESTRO based on (Fig. S1).

**Table S2.**
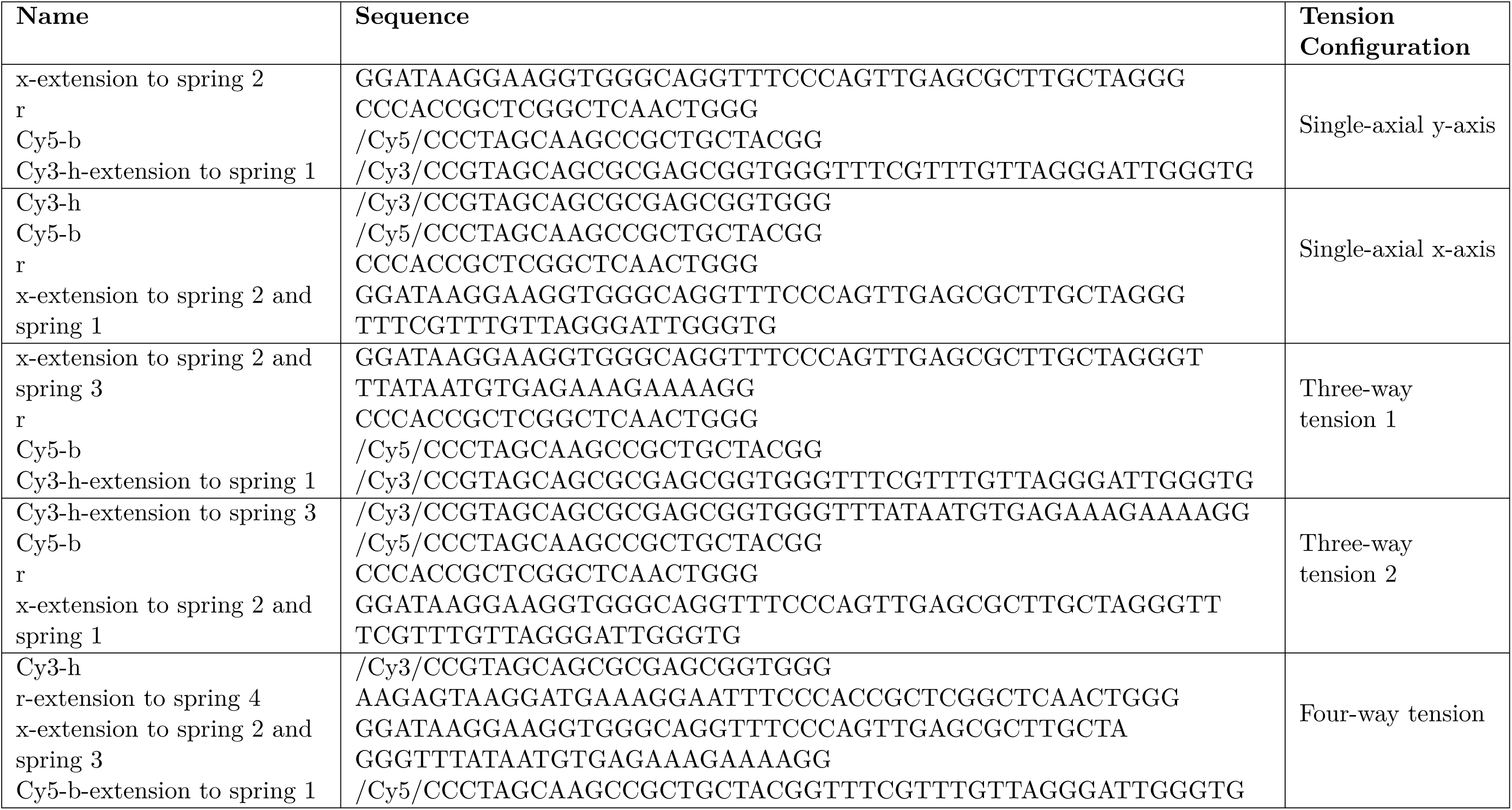
HJ strands for various tension forces configuration. Set of DNA strands to form HJ for specific tension force configuration. Each set consists of 4 strands in equimolar final concentration of 200 nM in a final 1× TAE 12.5 mM Mg^2+^.

**Table S3.**
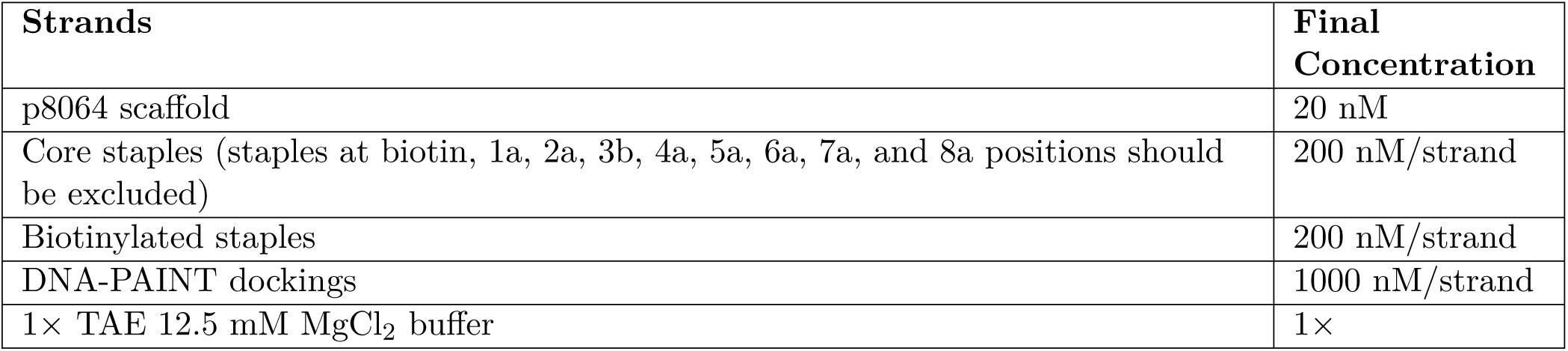
MAESTRO mixing concentrations for DNA-PAINT assay.

**Table S4.**
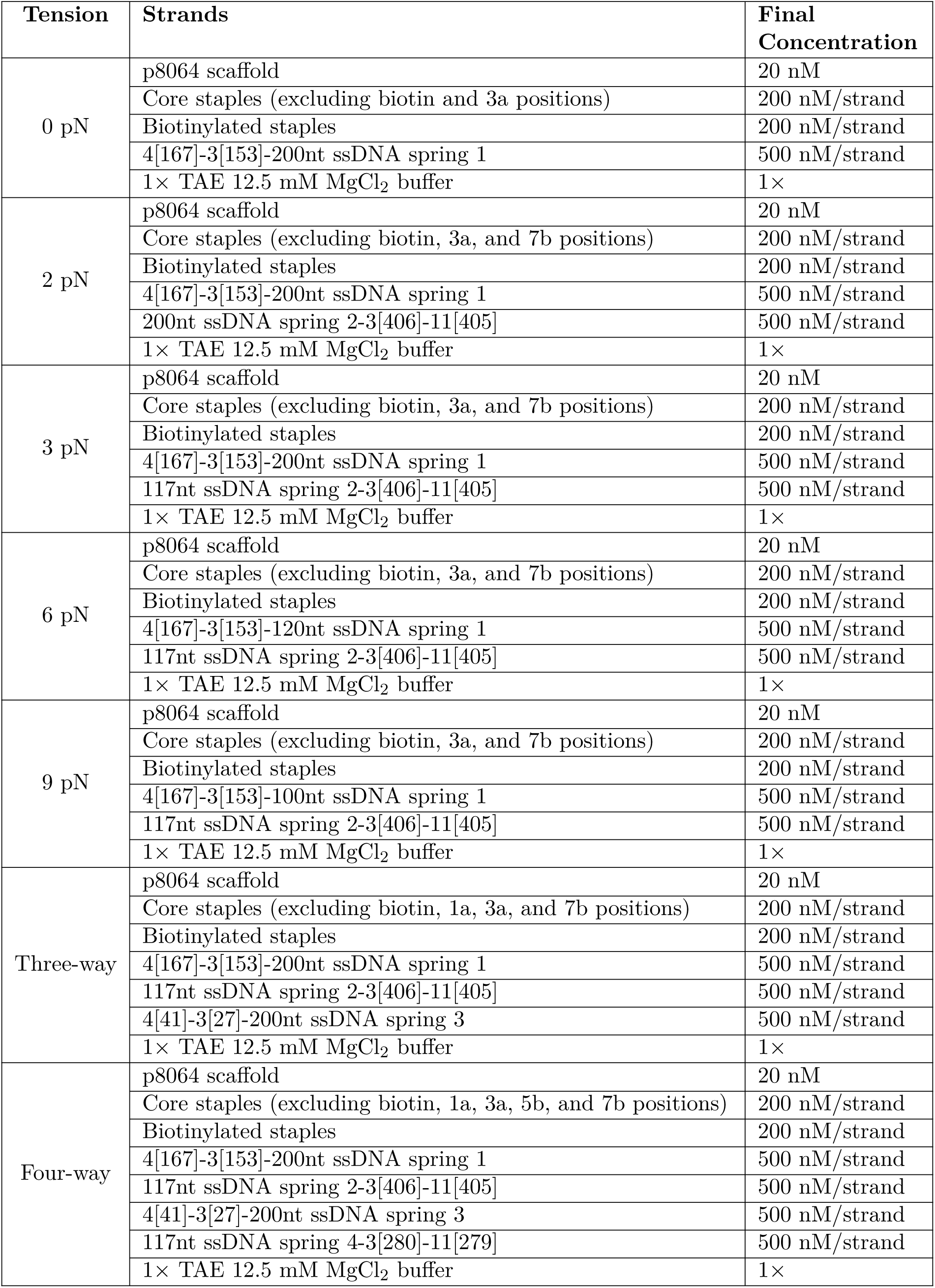
MAESTRO mixing concentrations across different tension conditions.

**Table S5.**
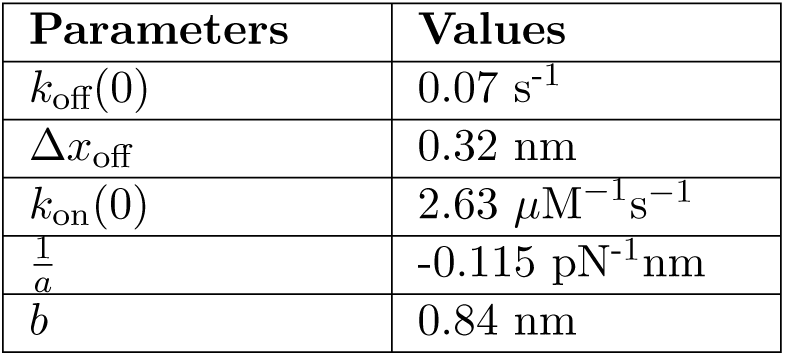
Fitted values of kinetics parameters for short oligos under tension.

**Fig. S1.**
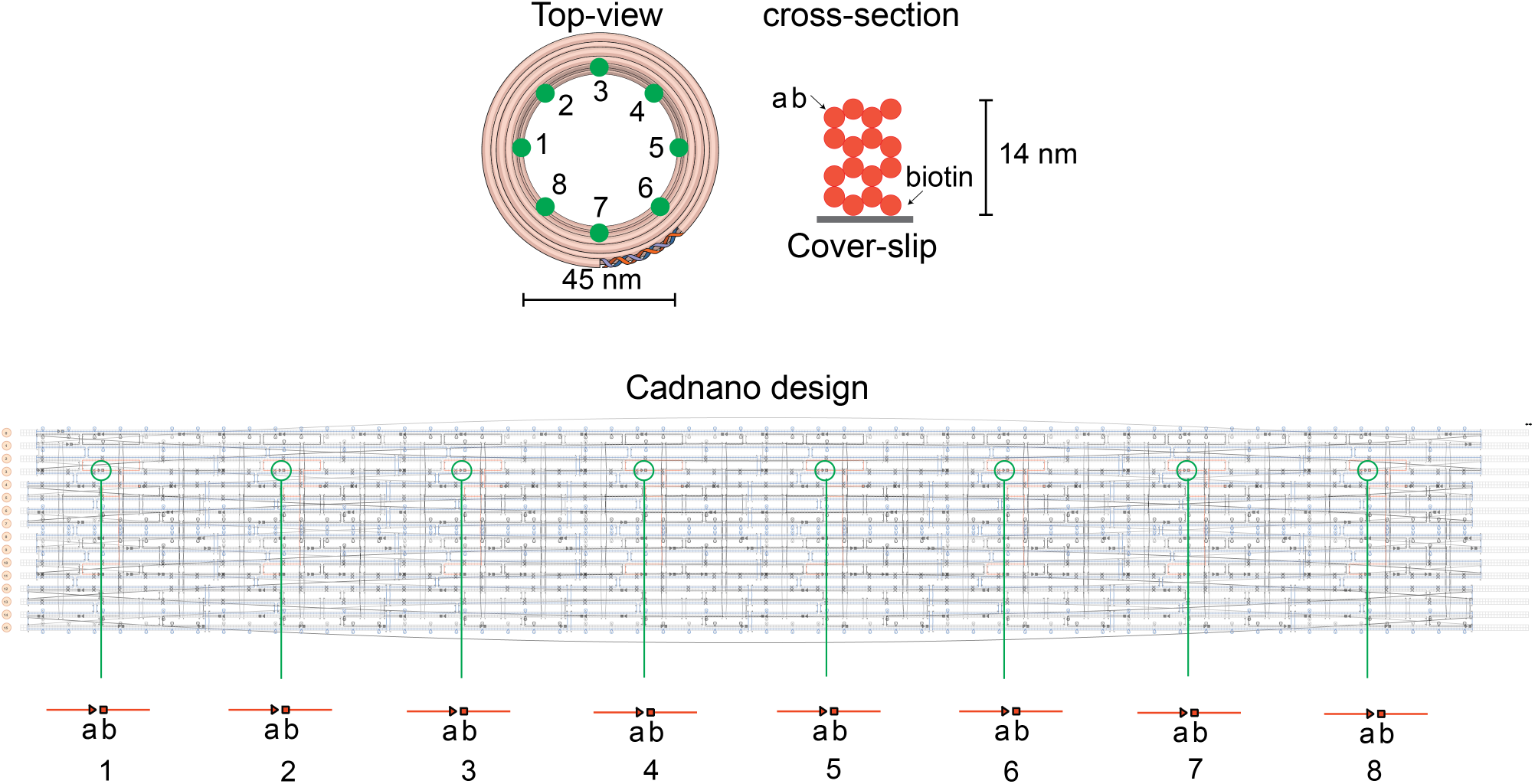
MAESTRO schematics (top) and cadnano design (bottom) showing the position of points 1a,b to 8a,b corresponding to some strands in Table S1.

**Fig. S2.**
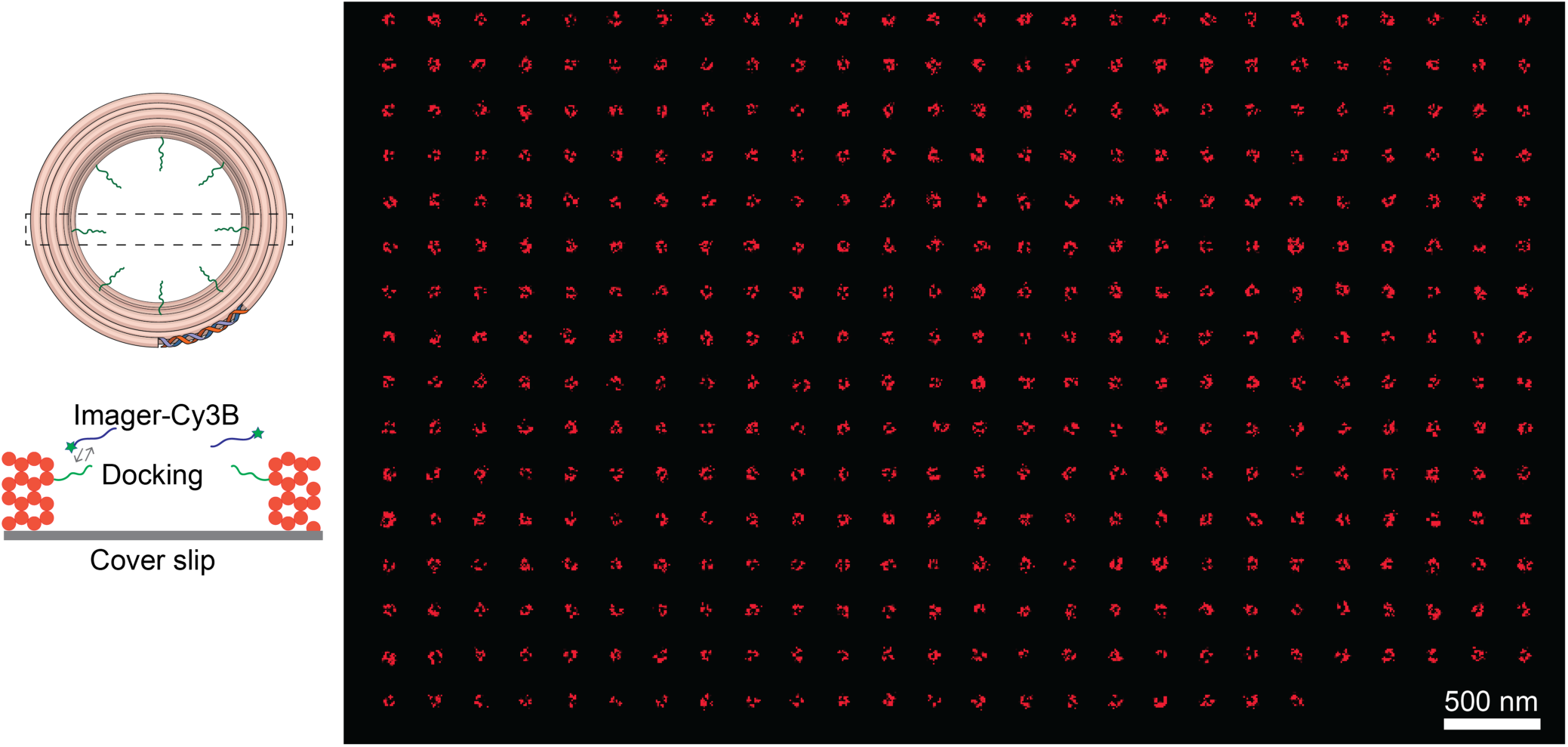
MAESTRO DNA-PAINT schematics illustrating the arrangement of docking strands and the organization of all selected molecules in an array.

**Fig. S3.**
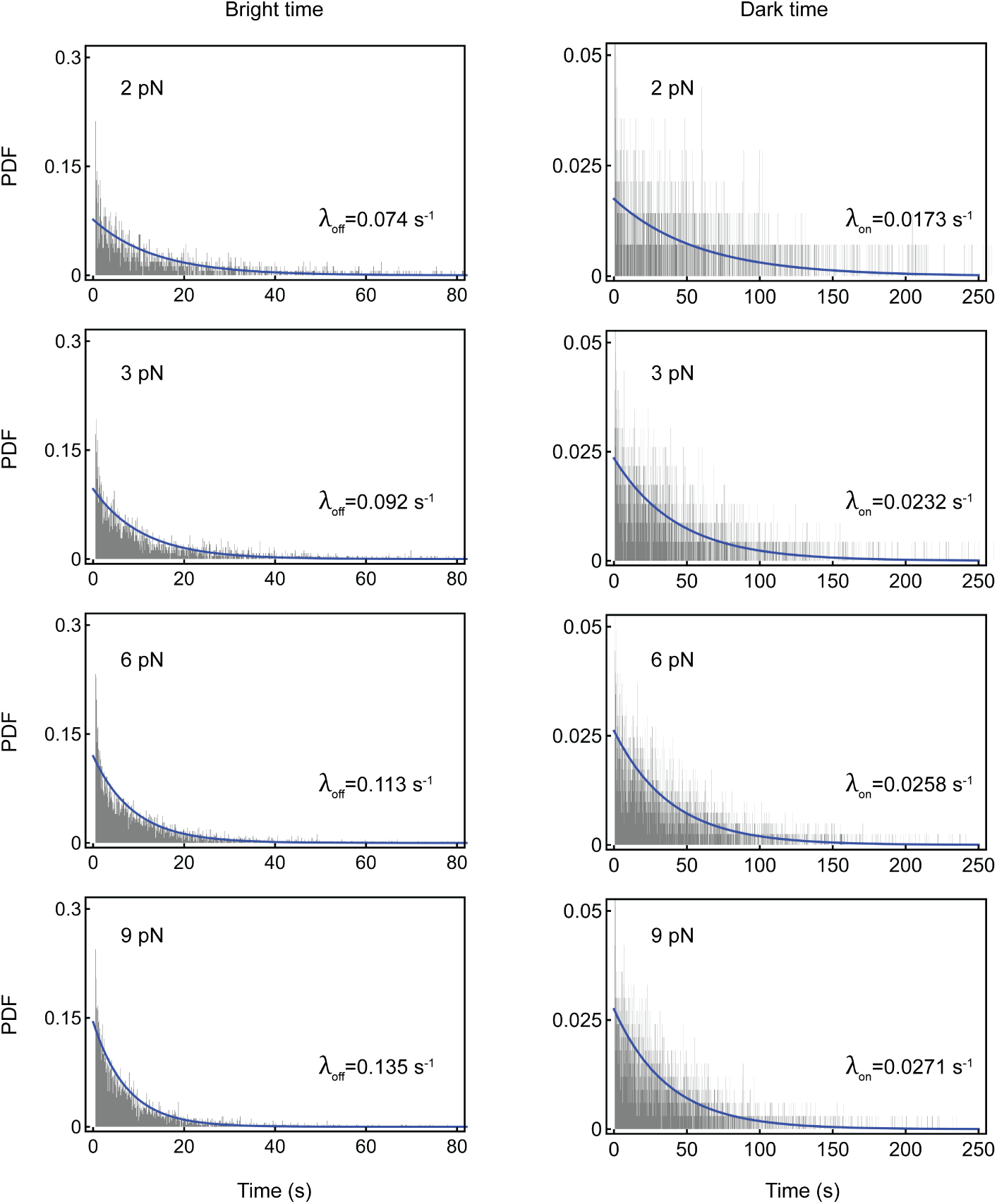
Histogram plots of bright (left) and dark times (right) of short oligos under various tension forces, using a bin width of 100 ms—equal to the camera exposure time—along with exponential fit curves (blue).

**Fig. S4.**
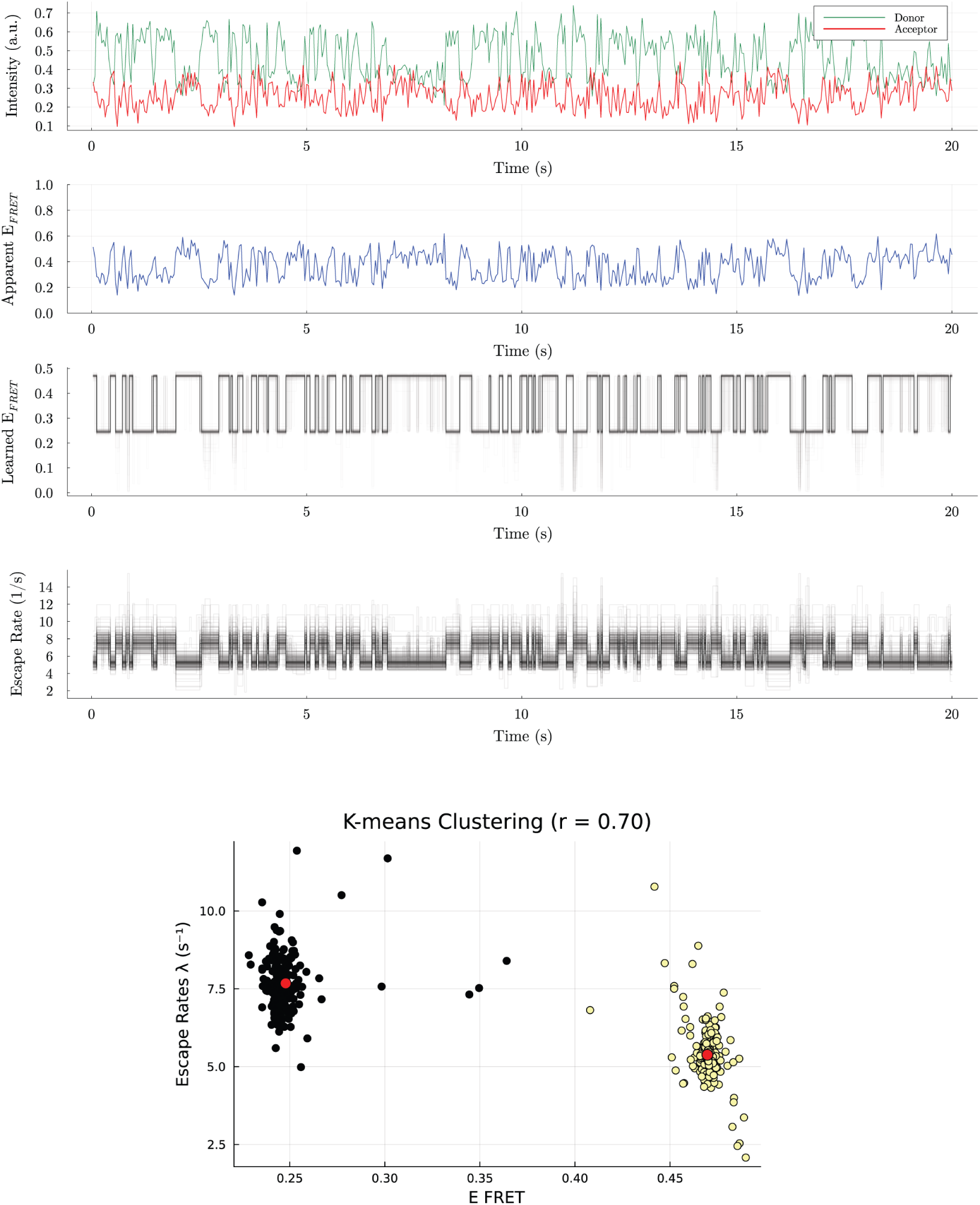
An example of a smFRET trajectory showing donor’s and acceptor’s emissions at 0 pN (1st row), the apparent FRET efficiency (2nd row), the learned FRET efficiency (3rd row), the learned escape rates (4th row), and the escape rates k-means clustering performed on MCMC samples generated by BNP-FRET to extract the mean escape rate λ values of Iso-I and Iso-II used to calculate the ratio r = λ_Iso-I_/λ_Iso-II_ where red dots show the mean values for each cluster colored in black and yellow (5th row)

**Fig. S5.**
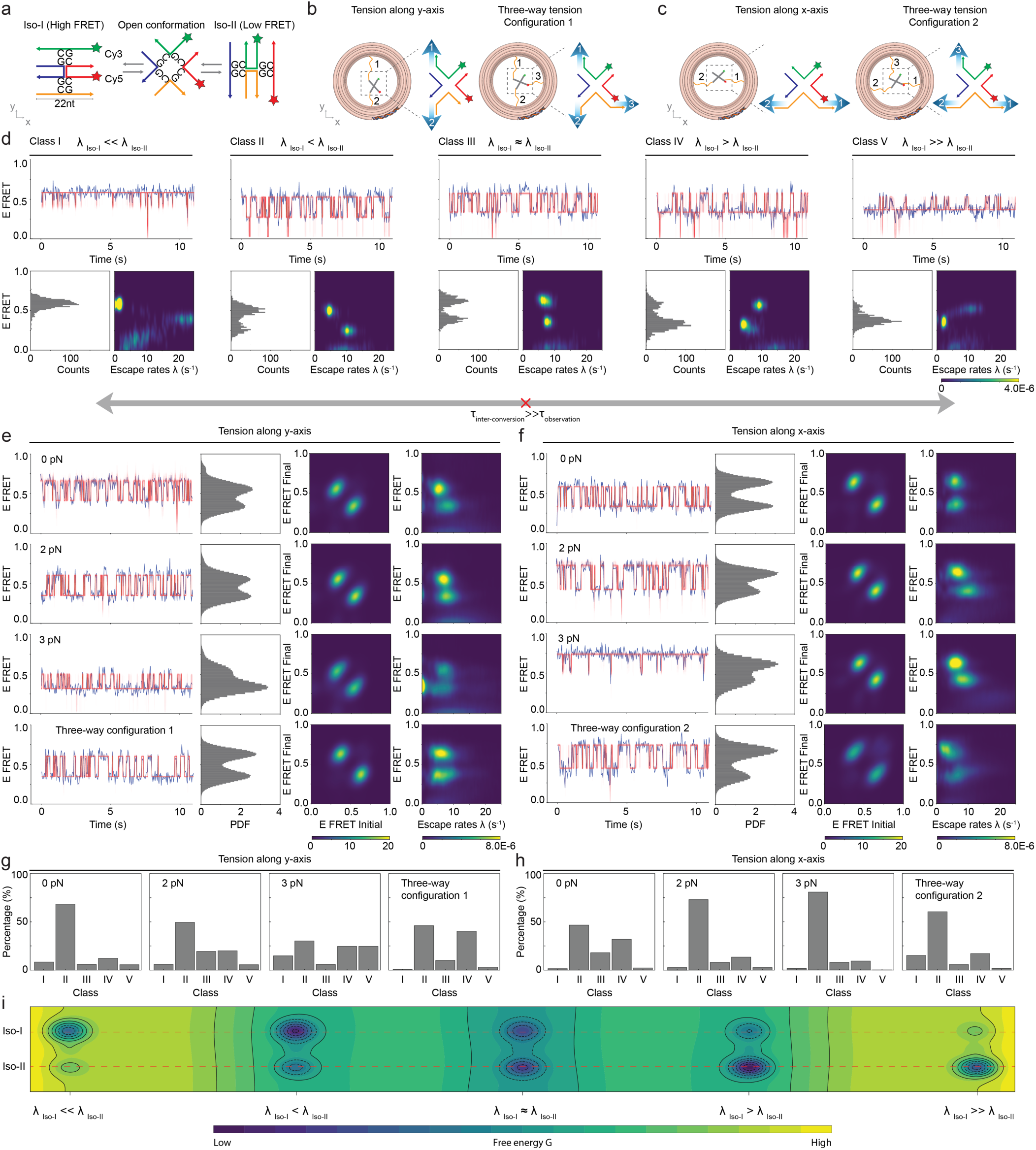
(Previous page) HJs under single-axial and 3-way tension forces configuration. <u>For comparison with main figures, panels d and portions of e,f repeat data from</u> Figs. 2 and 3. (**a**) Schematic of the J7 Holliday junction (HJ) show Iso-I, Iso-II, and open conformations, with a FRET pair (Cy3 and Cy5) attached to the ends of two arms. (**b, c**) Schematics illustrating the HJ connection to MAESTRO for different tension configurations: single-axial tension along the y and x-axes and two configurations of 3-way tension forces. Labels 1, 2, and 3 represent the distinct ssDNA entropic springs. (**d**) Five classes of smFRET trajectories grouped based on the relative escape rates of conformations. Top panels show representative smFRET (blue) and HMM (red) trajectories. Bottom panels show the corresponding smFRET histograms and escape rate heatmap. The absence of interconversion between classes suggests that τ_inter-conversion_ ≫ τ_observation_. (**e, f**) smFRET(blue) and BNP-FRET trajectories(red), ensemble smFRET histograms, transition density plots, and escape rate heatmaps compiled from n >150 molecules under different tension force configurations: (e) y-axis and 3-way tension forces configuration 1, (f) x-axis and three-tension force configuration 2. (**g, h**) Histograms showing the percentage of HJ molecules classified into each of the five dynamic classes under different tension conditions: (g) y-axis and 3-way tension forces configuration 1, (h) x-axis and 3-way tension forces configuration 2. (**i**) Qualitative visualization of the inferred energy landscapes corresponding to five dynamic classes.

Following the work by Hyeon et al. ^42^, where they are able to classify Holliday junction smFRET traces based on kinetics, here we use a similar empirical approach to categorize the kinetic behavior. We found that our data can be categorized into five distinct classes. Intuitively, these five classes can be distinguished based on the ratio of the mean escape rates, r = λ_Iso-I_/λ_Iso-II_. We identified the mean escape rate values using k-means clustering of MCMC samples for each smFRET trace(see Fig. S5). That is, we have extremal classes I (r < 0.2) and V (r > 5.0), where the Holliday junction gets stuck in either of the two conformations, Iso-I (high FRET) or Iso-II (low FRET); class III (0.9 < r < 1.1), where the Holliday junction switches between the two conformations at approximately the same rate; and classes II and IV, where the Holliday junction escape rates take intermediate values between the previously defined classes, as shown in Fig. S5(d) for a tension of 0 pN. These classes showed clear differences in their smFRET histograms and escape rate profiles (Fig. S5(d)). This variability likely arises from the rugged energy landscape of the junction, as previously proposed by Hyeon et al. ^42^, where complex interactions between Mg^2+^ ions and the junction structure create several kinetic traps. Consistent with their findings, our trajectories show no interconversion at 0 pN between classes within a 50 sec observation window.

Increasing the tension along the y-axis using springs 1 and 2 shifts the population toward the Iso-II state, thereby reducing the relative occupancy of Iso-I. This tension-induced population shift is clearly visible in the ensemble FRET histograms at 0, 2, and 3 pN (Fig. S5(e), column 2 panels). However, introducing spring 3 to apply 3-way tension forces balances the conformation populations, resulting in nearly equal occupancy of Iso-I and Iso-II (Fig. S5e, column 2, row 4).

Across all conditions, most molecules transitioned between two well-defined FRET states with efficiencies of E_FRET_ ∼ 0.3 (Iso-II) and E_FRET_ ∼ 0.6 (Iso-I), a hallmark of the canonical HJ behavior. ^46^ This dynamics is captured as two dense hotspots in the transition density heat maps (Fig. S5e, column 3). As the tension along the y axis increases, the escape rate distribution shows that λ_Iso-II_ changes to slower values, while λ_Iso-I_ changes to faster rates (Fig. S5(e)), column 4, compare row 1 to row 3). Under 3-way tension, however, the escape rates of both conformations become more balanced and span a wider range, in contrast to the asymmetric distribution observed at 0 pN (Fig. S5(e), column 4, compare rows 1 and 4).

We observe the opposite trend when tension was applied along the x-axis (Fig. S5(f)). That is, increasing the force from 0 to 3 pN favors the Iso-I state over Iso-II, as shown in the ensemble FRET histograms (Fig. S5(f), column 2, rows 1–3). Transitions predominantly occur between the two main conformations (Fig. S5f, column 3), but the escape rate behavior reverses: λ_Iso-I_ slows down while λ_Iso-II_ increases with tension (Fig. S5(f), column 4, compare row 1 with row 3).

We also note that in the case of 3pN tension along y-axis, the HJ gets stuck in Iso-II, as indicated by the bright spot near the 0 s*^−^*^1^ escape rate at low FRET efficiency (E_FRET_) in Fig. S5(e), row 3, column 4, respectively. We also observed this behavior when the tension was along x-axis, as observed in the trace shown in Fig. S5(f), row 3, column 1, albeit with far fewer molecules, as indicated by the lack of a similar bright spot at a high E_FRET_ in the corresponding ensemble escape rate heatmap.

**Fig. S6.**
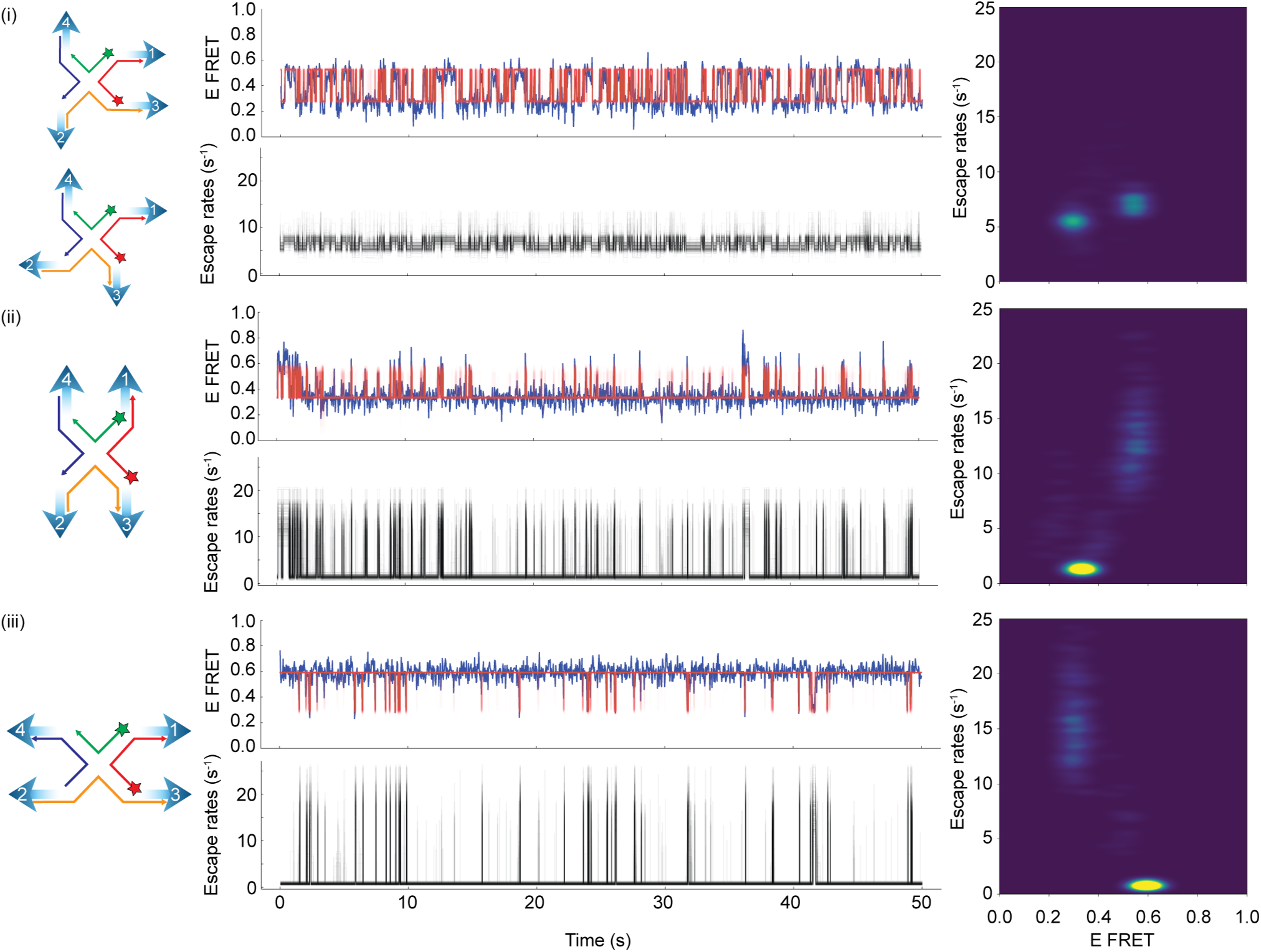
Additional examples of smFRET traces from the four-way tension force configuration, illustrating the proposed direction of tension forces on the molecules, along with their corresponding trace and escape rate heatmaps. These examples include traces that switch between Iso-I and Iso-II (i), remain stuck at Iso-II (ii), and remain stuck at Iso-I (iii).

**Fig. S7.**
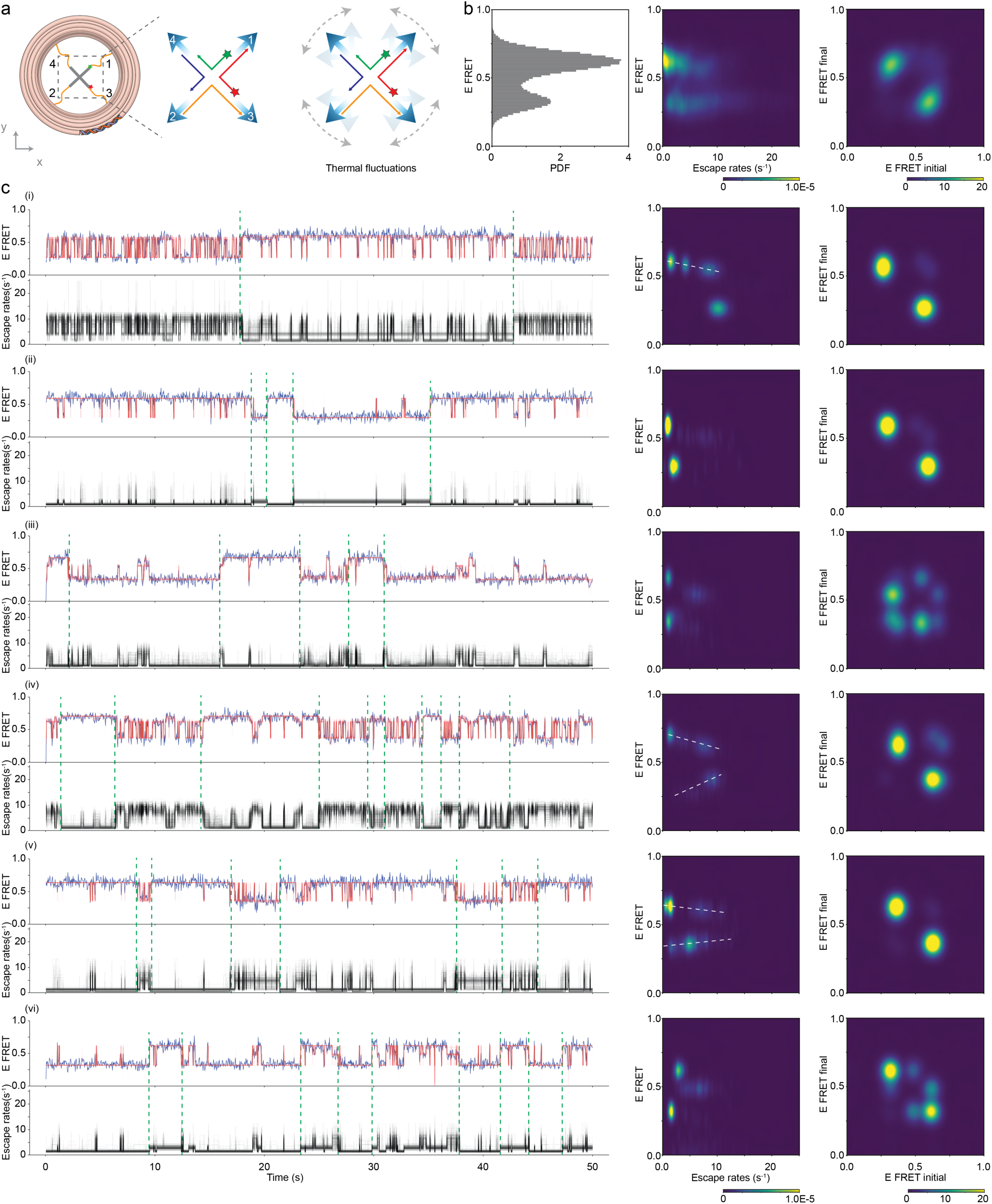
(Previous page) Interconversion between smFRET dynamic classes in J7 HJs under multi-axial tension from four-way tension forces. <u>For comparison with main figures, traces 1 and portions of 4, and 6 are repeat data from</u> Fig. 3. (**a**) Schematic of the four-way tension configuration on a Holliday junction (HJ), showing both the structural setup (left and middle panels) and the possible directions in which each entropic spring can stochastically pull the junction owing to thermal fluctuations (right panel). (**b**) Ensemble smFRET histogram, escape rate distributions, and transition density heatmaps (n=139 HJ). (**c**) Six representative smFRET trajectories (blue) overlaid with BNP-FRET trajectories (red) (top-left panels), accompanied by their corresponding escape rate trajectories (bottom-left panels), extracted escape rates (middle panels), and transition density heatmaps (right panels). Green dashed lines indicate visually identified interconversion points between smFRET dynamic classes based on changes in escape rate behavior. White dashed lines in some heatmaps highlight subtle shifts in E_FRET_ between distinct Iso-I and Iso-II conformations.

In the first example (Fig. S7(c)i), the molecule transitions between fast-switching states (class II, III, and IV) and class I, where the junction becomes mostly trapped in the Iso-I conformation. The corresponding escape rate trajectory and heatmap revealed multiple degenerate states with escape rates of ∼ 1 s*^−^*^1^, ∼ 4 s*^−^*^1^, and ∼ 9 s*^−^*^1^, all near the same FRET efficiency (E_FRET_∼0.6). However, states with faster escape rates tended to exhibit slightly lower E_FRET_. These Iso-I states transition with a single Iso-II state (E ∼ 0.25) characterized by a ∼ 10 s*^−^*^1^ escape rate. The transition density heatmap confirmed that major transitions occurred between these two main states.

The second trace (Fig. S7(c)ii) shows four interconversions between classes I and V within the 50 sec observation window. In the corresponding escape rate trajectory and heatmap, higher escape rates were observed during the transitions between Iso-I and Iso-II. The Iso-I state (E ∼ 0.6) exhibits a strong escape rate density at λ_Iso-I_ ∼ 0.5 s*^−^*^1^, whereas the Iso-II state (E ∼ 0.3) shows a prominent density at λ_Iso-II_ ∼ 2 s*^−^*^1^. The transition density heatmap confirmed that dominant transitions occurred between these two conformations.

In the third trace (Fig. S7(c)iii), five interconversions between class I (E_FRET_ ∼ 0.6, λ_Iso-I_ ∼ 0.5 s*^−^*^1^) and class V (E_FRET_ ∼ 0.3, λ_Iso-II_ ∼ 0.5 s*^−^*^1^) are clearly visible and separated by green dashed lines. The presence of intermediate transition states (0.3 < E_FRET_ < 0.6) is supported by multiple hotspots in the transition density heatmap, and these states exhibit faster escape rates in the range 5 s*^−^*^1^ < λ_transition_ < 10 s*^−^*^1^.

The fourth trace (Fig. S7(c)iv) shows ten interconversions between class I and faster-switching conformation classes (II and III). The Iso-I state in these fast-switching classes has a slightly lower FRET efficiency (E_FRET_∼0.6) than the class I Iso-I conformation (E_FRET_∼0.7), as observed in the escape rate heatmap. Transitions between the two Iso-I states are visible as faint hotspots in the transition density heatmap, along with the dominant Iso-I and Iso-II transitions.

A more complex dynamic behavior is seen in the fifth trace (Fig. S7(c)v), where the molecule interconverts between class I, class V, and class II. Multiple clusters in the escape rate plot for both Iso-I and Iso-II suggest dynamic heterogeneity under four-way tension forces. Additionally, Iso-II states with slower escape rates tend to exhibit lower FRET efficiencies, suggesting the presence of distinct energy levels. This observation is consistent with the behavior previously noted for Iso-I in the first and fourth trace.

In the sixth trace (Fig. S7(c)vi), a distinct transition state appears at E_FRET_ ∼ 0.5, visible in the smFRET trajectory, escape rate heatmap, and transition density heatmap. This state exists alongside clear interconversions between classes I and V.

Together, these examples demonstrate that four-way tension forces enable J7 HJs to explore a broader conformational energy landscape, facilitating multiple interconversions between dynamic classes within the 50 sec observation window. This is likely due to the simultaneous pulling of all four arms by entropic springs, allowing access to a wide range of force directions and conformational pathways. Such interconversions were not observed under tension-free and single-axial tension conditions.

**Fig. S8.**
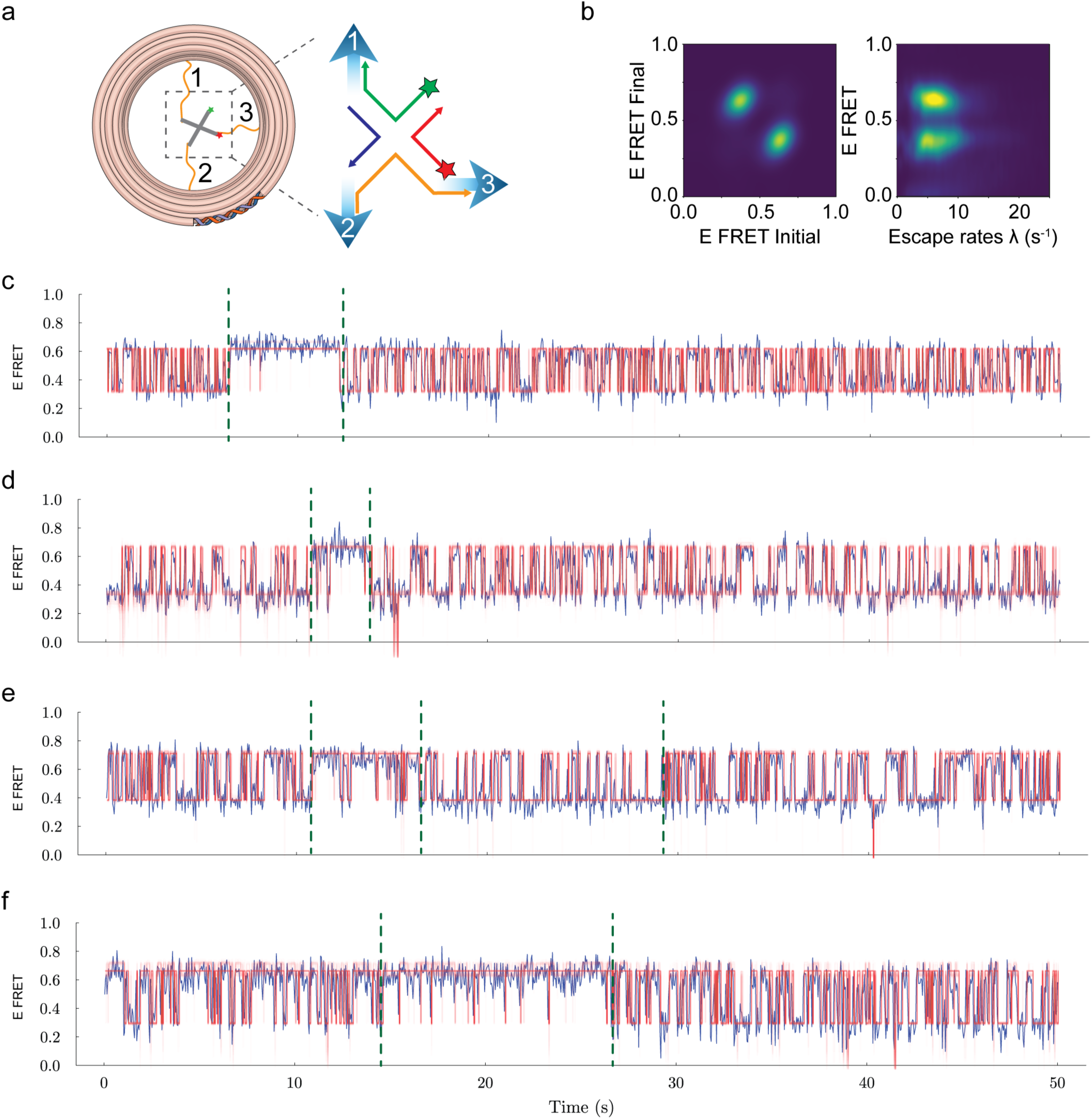
Kinetic class nterconversion in 3-way tension. (**a**) Schematic of 3-way tension on a Holliday junction. (**b**) Escape rates and transition density heatmaps. (**c-f**) Examples of FRET dynamics showing kinetic class interconversion. Green dashed lines indicate visually identified interconversion points.

**Fig. S9.**
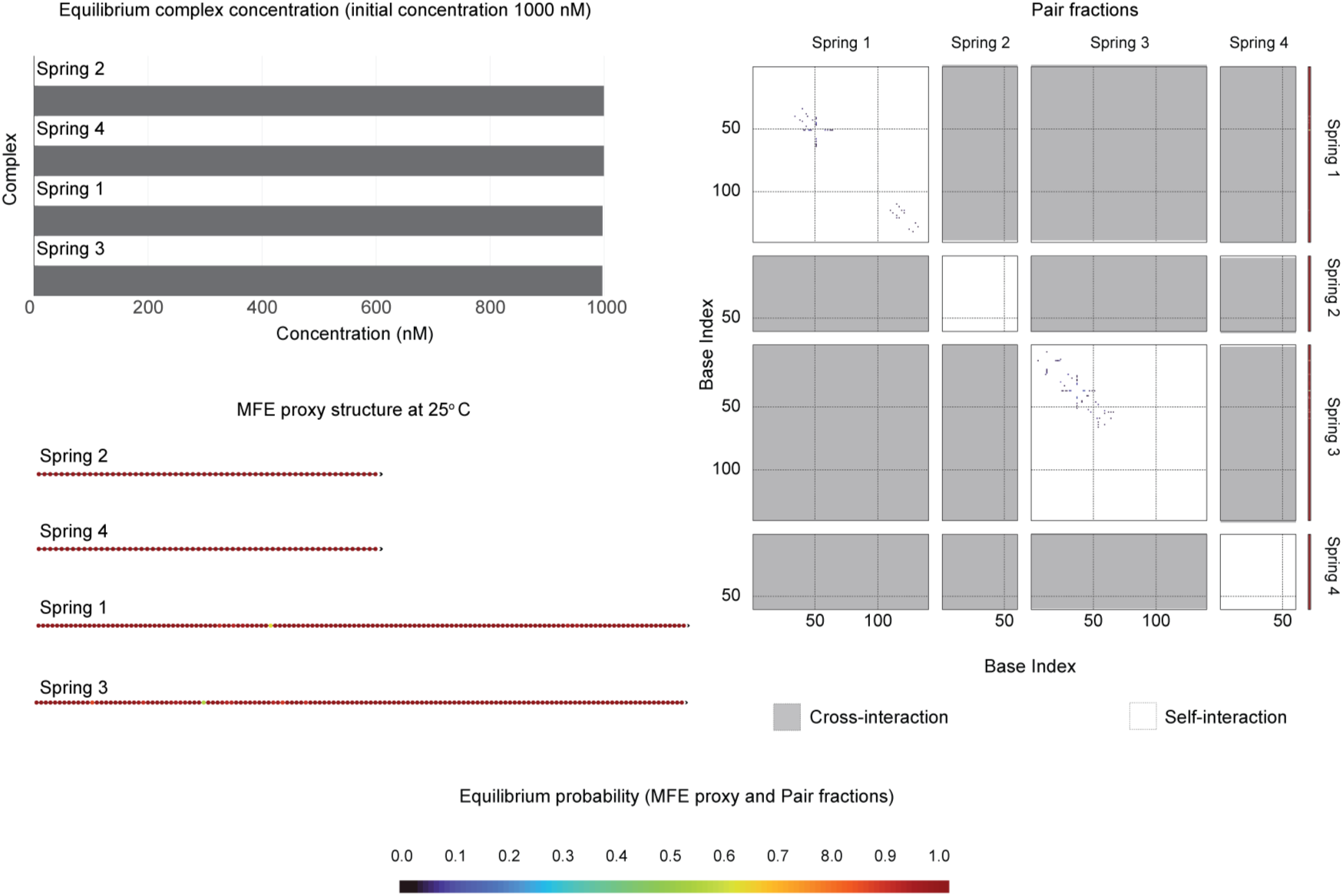
NUPACK analysis of the ssDNA springs at 25 *^◦^*C showing the equilibrium complex concentrations, pair fractions, and minimum free energy (MFE) proxy structures. The results indicate that all ssDNA springs exhibit negligible self- and cross-interactions, as evidenced by the MFE proxy structures and pair interaction profiles.

