## Supporting Information for "Multi-axial DNA origami force spectroscopy unlocks conformational dynamics hidden under single-axial tension"

| Name | Sequence | Note |
|---|---|---|
| 11[196]-4[210] | CAATAGATAAATTAATAGACAAACCCTCAATTGTAGCGA | Core staples |
| 6[447]-14[448] | AGAGGAAGCGCAGACGGTCACAAGAACTGACCTTACGAGAAA | Core staples |
| 14[20]-8[504] | CATCAACTTTTGTTCCATATTTTGTTAAAATTTTGCC | Core staples |
| 4[83]-3[69] | GAGCCGGACTTGTAGAACAGAAACAAAAAATCCCGTAT | Core staples |
| 8[216]-15[223] | AAAGAAGACAGGGAGAAACAATACAGATGATGAGTAACA | Core staples |
| 8[153]-15[160] | TATTACCGCGGTTGTAGCAATACGGTTGCTTAGTGTAGC | Core staples |
| 11[217]-9[226] | GTAAAAACCATAATTCATCAATATATGATTATACG | Core staples |
| 0[90]-8[91] | TGATATTCCCATCCTGTTTGATGGTGCCAGCTGCCTAATCCGGAAGC | Core staples |
| 3[7]-6[7] | TAGCATGTCACCAACGGTAATCGTAGCAAACAATGTATAAG | Core staples |
| 12[272]-11[258] | CACGCTCAACAGTAGGGTTCTGAATTAGATTTTCAGGT | Core staples |
| 14[195]-12[210] | TCCTTGCAGAGAAGTATTAGACTTTACAAACACATCAAAA | Core staples |
| 6[51]-11[48] | CCCGCGGCGAAATGTAAAACGACGGCCAATTAAGTTTCGCTAAGGCTG | Core staples |
| 3[343]-11[342] | AAAGCCCACAAAACCAGAACAGAATCAAGTGTAGCAC | 6b position |
| 0[279]-8[280] | TTAAGCAAAAACACCCTGAACAACTAACGAGTTTTGAAGTTTAGCGA | Core staples |
| 6[321]-14[322] | AACAAAGACTATCTTACCGATAAGACGGAATACCGAAACGCA | Core staples |
| 11[70]-4[84] | GCAGGCGCTTTTGCATCAGTGCTGGTCTGGTCAACCAGT | Core staples |
| 14[335]-8[315] | TATAAAACAAAAGAACAGGAAACGCAATAATAAGAAG | Core staples |
| 9[290]-4[294] | TACCGAGGCGTCCTTAAATCAAGATTGAATCTTATTTGCCTCCCAA | Core staples |
| 14[321]-12[336] | AAGACGAAAAGAAAATACATACATAAAGGTGGACAAATCA | Core staples |
| 15[252]-14[273] | AAAGAGGCATTTTCGAGCCAGTAATAAGAGACCAACAA | Core staples |
| 4[272]-3[258] | TTTCTTCTGACCTAAATTTAATTAAGAATTTATCAATA | Core staples |
| 11[133]-4[147] | CCTAAAGGGAAGAAGCACTCCAACGTCAAAGTCAATACG | Core staples |
| 6[177]-11[174] | GGATACCTACATCTATCGGCCTTGCTGGATTAGTAAGTCTGTTGAGAA | Core staples |
| 8[468]-15[475] | CAATACTGAGGACGACGATAAAACGTTGGGATTTAATTT | Core staples |
| 9[416]-4[420] | TAATTAGTTGCGATATATTCGGTCGCGATCGTCGGGTAGCACTTTT | Core staples |
| 11[227]-0[217] | AAATAACAGTAAATAACCGTACATACTATATGGACCCTGTAATACTT | Core staples |
| 11[154]-9[163] | TACGCCAAGGGAGAGCACGTATAACGTACTATTTC | Core staples |
| 4[356]-3[342] | TGATATTAGAATGGTTTGATGATACAGGAGTGTTCTCATT | 6a position |
| 11[490]-9[505] | AACGCCAGGTAGAACTAACGGAACATAAGAACGCAAAAGAA | Core staples |
| 11[91]-9[100] | GTCATAAATGCGTGCATCAGACGATCAGCGGTTCG | Core staples |
| 6[195]-14[196] | AATCAAAACACCTCAAATATTTAGAGCTTTAGGAGTATTAAA | Core staples |
| 14[132]-12[147] | GAAAGATGCCTTGACGGGGAAAGCCGGCGAACACTTTTAG | Core staples |
| 15[287]-11[289] | TACCGACAAAAGGTATGTTCAGGATAAGTAATTTACGAGCATTTC | Biotin position |
| 4[293]-3[279] | TCCAAATAATGAAAGACGGGAGAATTAACTGACTTTTAAC | 5a position |
| 7[21]-1[16] | CACGGGAACTAAACAGGAAGATGAG | Core staples |
| 4[398]-3[384] | AAAAACATGAAAGTATTGCAGGTCACTGAATTTACCCC | Core staples |
| 0[321]-6[304] | ATTAACAGAGAGATAGAACATAAAAACAGGGAAGTGAGCGCTTTTA | Core staples |
| 3[448]-6[448] | CCATGTTACTTTGAAATCCGCGACCTCGCCTGAACTTTGAA | Core staples |
| 15[315]-14[336] | ACACCACGGAATAAGTTTATTTTGTCACAATTGCAACA | Core staples |
| 3[469]-11[468] | GAGCTTTTCAAAGACTATCTCGTTTACCATACCACAT | 8b position |
| 6[258]-14[259] | ATAAGGATATTAATGGTTTGTCTTACTTTAGTATATGTAATT | Core staples |
| 11[164]-0[154] | ATCCCCATCACCCAGTAAATTGGCATAAAACATATTTTAAATGCAAT | Core staples |
| 6[384]-14[385] | TTTTGCAAGAAGAGGCTGAGATCACCATATAAGTCCACCACC | Core staples |
| 3[28]-11[27] | TGGGCGCTGCTCGTGGTGAGCTGGCGAAAATAAAGCG | 1b position |
| 12[461]-11[447] | AACCCAAATCAACGTAAAAATGAATCTAAAGGAATTGG | Core staples |
| 8[279]-15[286] | ACCTCCCGGTTCATCGTAGGAATAGAACGCGATATAAAG | Core staples |
| 14[272]-8[252] | CGCCAACCATATGCACATTACTAGAAAAAGCCCAAAA | Core staples |
| 11[7]-4[21] | AATAGGAACGGCCAGCTCAGAAAAGCCCCAACCATTTGC | Core staples |
| 12[146]-11[132] | ACGCCCCCGATTTAGAGCTGTGCACCGCGTGCCTGTAC | Core staples |
| 11[301]-9[316] | CAAGAACATCCTCCTGAACAAGAAACGACAATCAAATCAGA | Core staples |
| 9[506]-1[503] | GTTCGAACGAGGCATAGTAATTGCATAAACGAGGCTGAAT | Core staples |
| 9[65]-0[56] | TTCGCCAGCGATCGGTGCGAAAGAGATGTTTACCCGTTTTTGCCGGAG | Core staples |
| 0[216]-8[217] | TTGCGGGACTTTGGGTTATATAAAATCAATACATCAAGACCTGAGCA | Core staples |
| 3[133]-6[133] | TTTGGAACAAGCGTTGAGTGTTGTTAAGAATAGTCTATCAG | Core staples |
| 11[322]-4[336] | TATTACGCAGAGGCATGATAGCCCTTTTTAACCCAAATA | Core staples |
| 8[377]-7[398] | GCCCCCTTTCAGAACCGCCTCAGTACCAGGCGGATAACA | Core staples |
| 15[161]-11[163] | GGTCACGCTGCGCGTAGGGCGCGTGCTTTAGGAGGCCGATTAAGA | Biotin position |
| 14[6]-12[21] | TGAGCTTACCGCGTCTGGCCTTCCTGTAGCCAGGGAAACC | Core staples |
| 0[307]-4[305] | ATCATACAGGCAAGGCGGGTAATCGCATTAATAGCAGCCTTTTAT | Core staples |
| 8[62]-7[83] | CACGACGTCGTACAGCGCGGCAGCACCGTCGGTGGTGAA | Core staples |
| 11[49]-9[64] | CGCAACCTTTCCCAGGAAGATCGCATAGATGGGCCAGGGTT | Core staples |
| 11[28]-9[37] | CCATTCGTGCCGGACGACGACAGTAGCATCTGCAA | Core staples |
| 3[217]-11[216] | TAAGACAAAACATGAGTGCCTTTTACATCACTTGCAC | 4b position |
| 4[461]-3[447] | GTTAGCCGGAACGAGGCTTTGAGGACCAACCTAAAACT | Core staples |
| 12[20]-11[6] | AGCCATCAAAAATAATTATTATTACAAAAGGAATTACC | Core staples |
| 0[468]-8[469] | CTGGAAGTTGAGAGGTCATTTTTGGTCTTTAAAATATTCATAGCGTC | Core staples |
| 0[118]-4[116] | AAGGCCGGAGACAGTCCGAAATCCGCTGGTCTTCACCGCCTGGGG | Core staples |
| 4[178]-15[188] | CCTGACATTTAAAAGATAACATCGCCGCTACAACCACCACACCCGCC | Core staples |
| 11[238]-9[253] | ATTGCGAATGGATTGTTTGGATTATAGCGGAAAATACCAAG | Core staples |
| 14[209]-8[189] | ACAACTCGCACTAATGGAAGGTTATCTAAAATAGAAG | Core staples |

| | | |
|---|---|---|
| 0[27]-8[28] | CTATCAGGGATAAACGATGCTGATAGACTTTCGGATAACGCTTTCAG | Core staples |
| 3[259]-6[259] | GTTAATTTCATGTTTTCAAATATATCAAAGAACGGCGTTAA | Core staples |
| 9[353]-4[357] | GACTTTTTCATCGCCTCCCTCAGAGCCCCTCAGCCAGCATTGGCCT | Core staples |
| 1[504]-6[493] | ATAATGAAAACAGGTCAGGATTAGAGAGCTTAATTAATG | Core staples |
| 11[175]-9[190] | GTGTTTCTAAACCCTCGTTAGAATCTTAATGCACTTGCCTG | Core staples |
| 7[399]-0[385] | TCGCCCACAGAAGGATTAGGATGTTAATGCGCAAATGGTCAATAA | Core staples |
| 6[6]-14[7] | CAAATGACAGTTGATAATCATTTTTTCATTAAATATTAAATG | Core staples |
| 0[496]-4[494] | ATGTTTTAAATATGCAGCTTAGAGTACCTTAACCAGACCGGAAAG | Core staples |
| 6[69]-14[70] | CCAACCACGCGTCAGCGTGGCGGGGTTTACGGCTCCCTGCGG | Core staples |
| 14[69]-12[84] | CTGGTTGCTCGGGCGCGGTTGCGGTATGAGCCCCGCGGGC | Core staples |
| 15[504]-14[21] | ATGAGTAACAACCCGTCGGATTCTCCGTGGGAGGCTTT | Core staples |
| 0[132]-6[115] | TTCAAAAAAATCAACCGTTGCAGCAAGCGGTCCAGGCAAAATCTTT | Core staples |
| 1[17]-0[497] | AATCTCATTGCCTGAGAGTCTGGCTGTAGCTCAAC | Core staples |
| 6[132]-14[133] | GGCGACATTGAACGTGGACTAAATCGTTGGGGTCGAAGGGAA | Core staples |
| 11[259]-4[273] | ATAAAGCCAAATTACAAATAAATACCGACCGTCCGATTT | Core staples |
| 0[69]-6[52] | AAATTAATTCGTCTTCAAAAAAAGCCGCACAGGCAACGCGGTCAGT | Core staples |
| 0[55]-4[53] | AGGGTAGCTATTTTTGTCCGGCAGGCCTTTACATCGACATAAGCG | Core staples |
| 4[482]-3[468] | GAAAGACCAAAGCGTAATTGCTCCTTTTGATATCTAATTC | 8a position |
| 3[280]-11[279] | GTCAAAAAGAAATTTCCACGTTTTTATTTTAAATCGG | 5b position |
| 9[227]-4[231] | GATTTCAATTAAAACAAAATTAATTAGGAAACATTGCTTCCTTAGA | Core staples |
| 9[380]-0[371] | CGGTGGTAACCATCGATAGGAGGTTGCCCTCAGAGTAACAAGCTATAT | Core staples |
| 0[433]-4[431] | TGCGAACGAGTAGATTGAAACAAACACTAAACTACGAAGGCACTA | Core staples |
| 11[343]-9[352] | CATTACCCAGCAGGAAATTATTCATAGGGAAGTCA | Core staples |
| 4[367]-15[377] | GATTGACAGCAGCACGTTTTCATTCAACCGACCAGCGCCAAAGACAA | Core staples |
| 4[20]-3[6] | CGATCATATGTACCCCGAAAAGATTAGCAAACTCCAAC | Core staples |
| 15[441]-14[462] | ACGAACGAGTAGTAAATTGGGCTTGAGATGGAGAGGCT | Core staples |
| 7[273]-0[259] | GCGGGAGGCGTGTGATAAATAAGCGAGAATAAAGCCTCAGAGCAT | Core staples |
| 0[342]-8[343] | ATCAATTCAGTACTGGTAATAAGACCAGAACACCGGAACAATCAAAA | Core staples |
| 11[416]-0[406] | GAATAAATCTCGGAACGAACCCTCACAAGCGCTAGTTTGACCATTAG | Core staples |
| 8[503]-7[20] | AGAGGGGGAAACAGTTCAATTTAAATTGTAAACGTTAGC | Core staples |
| 15[350]-11[352] | AATTCATATGGTTTATTGAGGGTAAAGGTTTGGGAATTAGAGCAT | Biotin position |
| 8[188]-7[209] | AACTCAAATTTGACGCTCACAGTTGAAAGGAATTGAGAT | Core staples |
| 0[447]-6[430] | GTTGATTTGTATCATGCGAAAGAGGCAAAAGAATAGTACAACTTTT | Core staples |
| 11[353]-0[343] | TAGCAGTAGCGGCCGCCGAGCCGCCTTTTAACAGCTGAAAAGGTGGC | Core staples |
| 12[335]-11[321] | CCTATGTTAGCAAACGTTAATATCCCGGGTATTAAACT | Core staples |
| 9[128]-0[119] | GAATTGATCTTCGCGTCCGGAGGCGGCTGCCCGCCCTTATGGGTGAGA | Core staples |
| 15[35]-11[37] | GCGGATTGACCGTAATAACCGTTCGGCCTGGCACCGCTTCTGGCC | Biotin position |
| 9[101]-4[105] | TAATATACGAGGAGTGAGCTAACTCATCGGGAACAACGCGCAGGGT | Core staples |
| 9[164]-4[168] | TTTGTAATATCAACGCTCATGGAAATTATTTACTAAAAGGTCTGAC | Core staples |
| 0[370]-4[368] | TTTCATTTGGGGCGCGGGGGTCACATGGCTAAAGCGCAGTCTGAC | Core staples |
| 11[427]-9[442] | GAACAATTTCTGTTTCCAGACGTTACGCCTGTGCTTGCTTT | Core staples |
| 15[476]-11[478] | CAACTTTAATCATTGTATACCATAAAACGAAAGATTCATCAGAAT | Biotin position |
| 4[493]-15[503] | AGGAAGCGGAGAGCAAGGCTTTTTGGCTCATTGAATTACCTTATGCG | Core staples |
| 15[378]-14[399] | AATTTCAGGGATAGCAAGCCCAATAGGAACCGTCCACC | Core staples |
| 4[419]-3[405] | TCATGAGTAATGCCAACACTCATCTTTGACCCCCGGGTAA | 7a position |
| 0[244]-4[242] | GTACCAAAAACATTATTAAATGCCTACCTTAGAGTCAATAGTTTT | Core staples |
| 4[209]-3[195] | TAGCATCACCTTGCTGAAGAGATAGTTAATGCGCGAGA | Core staples |
| 6[492]-11[489] | ACCAAATGCTTTTAATAGTAAAATGTTTTAGCGAGACACTATATACAT | Core staples |
| 9[443]-0[434] | CGAGGCCGCGAATAATAATTACAGAGAGGCCGCGGAGATTCCCAATTC | Core staples |
| 8[405]-15[412] | TGACAACAGGCTCCAAAAGGAGCTAGTTAGCCATGTACC | Core staples |
| 15[189]-14[210] | GCTGCCCGAACGTTATTAATTTTAAAAGTTTGGATTCG | Core staples |
| 4[104]-3[90] | GGTTTTTGATTGCCTTGCCCCAGCAGGCGAAAGGCGGGCA | 2a position |
| 3[385]-6[385] | TATTATTCTGACCTGCCTATTTCGGTATAAACATAGCGGGG | Core staples |
| 4[304]-15[314] | TTAAGTTACCTCATCATAGCAAGAAACAACAAAGTAATTCTGTCCAG | Core staples |
| 11[101]-0[91] | GGGGAGTTGAGAATCGGCACCTGTCGTGGTTCAAATCACCATCAATA | Core staples |
| 14[146]-8[126] | GAGAAAGGAGGTGCCCTCACCCAAATCAAGTTATTGT | Core staples |
| 3[70]-6[70] | AACGGAACGTGACCAGAGCACATCCCGTCGCTGCAAGAATG | Core staples |
| 4[41]-3[27] | TAAATTTGTTGTGTAGTGATGAAGGGTAAAGTTGAGCAGT | 1a position |
| 6[366]-11[363] | ACCACGCCACCCATTAGCGTTTGCCATCGTAGCGCCGTAATCAAGGCC | Core staples |
| 4[115]-15[125] | CGCCGGGGATGAGCCATAGCTGTCGGGTTACCAGCAAATCGTTAACG | Core staples |
| 6[429]-11[426] | GCGGTGAGGCTTAACAGCTTGATACCGATGTATCGACGTTGAAGAAAG | Core staples |
| 0[258]-6[241] | AAAGCTACGCAAGATTAATCATAGGTCTGAGAGATGATGCAATTTT | Core staples |
| 7[84]-0[70] | GCCTGGGGTGCAGCAGCAACCGGCAGCCTAACCGTTCTAGCTGAT | Core staples |
| 15[224]-11[226] | TTATCATTTTGCGGAATATTCCATCCTGAAGGGTTAGAACCTCAG | Biotin position |
| 3[406]-11[405] | AATACGGAAGTTAGCATCCAAAAAAAAGGAAGTTTCA | 7b position |
| 4[335]-3[321] | AATGAAATAGCAATAGCAGCCATATACAGAGAGAATAG | Core staples |
| 11[112]-9[127] | CCAGCATCTGTGAGTGTCACTGCGCCAGATGCTTCCTGTGT | Core staples |
| 3[91]-11[90] | ACAGCTCTTTTCTTAATGGATCCCCGGGTCATTCACG | 2b position |
| 4[52]-15[62] | GATACGGAAGGCCTCTGGGTAACGCGCATCGTGGGATAGGTCACGTT | Core staples |
| 7[147]-0[133] | CCATTGCAAGGGCGAAAAACCGCCCGAGAAATGTGTAGGTAAAGA | Core staples |
| 4[146]-3[132] | TGGAGTCCACTATTAAATGCGTATTGCCCTGAGAGAAG | Core staples |

| | | |
|---|---|---|
| 7[210]-0[196] | GAAACAAATACAATATCTGGTCTGCCACGGAAGCCTTTATTTCAA | Core staples |
| 9[254]-0[245] | TTATGCAGTTTAACGTCAGTCGTCGCATTACCTATCCAATAATCGGTT | Core staples |
| 9[317]-0[308] | TATACTCCCAAGTACCGCAAAAATAACTACAATAATATCATCCAATAA | Core staples |
| 3[154]-11[153] | AATATTCGTAAGACACGAGCAAATTAACCTAGAACGG | 3b position |
| 12[83]-11[69] | CGTCGCACTCAATCCGCCCAGCCAGTGTTGGGAAGGTT | Core staples |
| 4[230]-3[216] | ATCCTTGGCTGAGATTTAACCTCCGGCTTAGGGATTAGAT | 4a position |
| 14[83]-8[63] | ACTGTTGGGAGGTGTACCATCCCACGCAACCACCAGT | Core staples |
| 8[90]-15[97] | ATAAAGTGTCACCGAGCTCGAATGCCGGTGCAACATCCC | Core staples |
| 11[280]-9[289] | CTGTCTGTAGAATTTATCAACAATACTAATGCCAT | Core staples |
| 12[209]-11[195] | TTTACATTTGAGGATTTAGCGGGAGTTATAATCAGTGT | Core staples |
| 11[448]-4[462] | ATATTCATTATAATCTTGAATCATAAGGGAACTTATCGC | Core staples |
| 11[38]-0[28] | ATTCTTACGCCAAGGGATTGAGAGATTGCCGTAGAGATCTACAAAGG | Core staples |
| 4[241]-15[251] | TCCTGTAAAATGAATTTGCTTTGTTATCATCACAAAGAAACCACCAG | Core staples |
| 9[38]-4[42] | GGCGGTGCCAACTCACCGGAAACAATGAATTTGAGCTCTCCAAACT | Core staples |
| 11[290]-0[280] | CTTAAGCAAGCGAGCCTAACCAACGAGTCAGAAAAGAATTAGCAAAA | Core staples |
| 11[479]-0[469] | GCAGCATAACCTATAGTCAAAATCAGCGGATGACTAAAGTACGGTGT | Core staples |
| 8[125]-7[146] | TATCCGCTGCGTTGCGCTTGGCCCACTACGTGAACCAAG | Core staples |
| 11[469]-9[478] | TCAACTTTGAGAAAATCTACGTTAAGTCAGGAACC | Core staples |
| 14[258]-12[273] | TAGGCGGACCTTAATTGAGAATCGCCATATTTTGACCAAT | Core staples |
| 3[196]-6[196] | AAAATCTAAAGCGAGCCAGCAGCAATGCAACAGAGTTGGCA | Core staples |
| 11[364]-9[379] | GGAAACAGCCATGAATTATCACCGTGCGACATCGGCATTTT | Core staples |
| 6[114]-11[111] | CCAGCATTAATTCACAATTCCACACAACCATGGTCTCCTCACTTTCTG | Core staples |
| 0[195]-6[178] | CGCAAGGCACCGCCATACTGATAGCCCTAAAACAGGCGGTCAAAAT | Core staples |
| 7[336]-0[322] | GAGCCACCCAGAAAAGTAAGCAAGAATTGTACTAATAGTAGTAGC | Core staples |
| 14[384]-12[399] | CTCATGGCACAGAACCGCCACCCTCAGAACCGAAACTTTC | Core staples |
| 14[447]-12[462] | CACCAAAGTCAAAGCTGCTCATTCAGTGAATAAATTTAGG | Core staples |
| 0[153]-8[154] | GCCTGAGTTACACCAGCAGAAGAGATTCACCACAGGAAACAGAACAA | Core staples |
| 6[240]-11[237] | TAATCATTTAACAGGCGAATTATTCATTCGCCTGAATACAGTAAAGAA | Core staples |
| 8[440]-7[461] | ATTTCTTAGCAGGGAGTTACAGATGAACGGTGTACAGGA | Core staples |
| 6[303]-11[300] | TCCTAGTTGCTAGGTATTCTAAGAACGCGCGCCCAGAGAACATCATTC | Core staples |
| 15[63]-14[84] | GGAATGGGTAAAGGTTTCTTTGCTCGTCATACCGGGTC | Core staples |
| 8[314]-7[335] | GCTTATCCTTTTGCACCCAGTTACCAGAAGGAAACCGCA | Core staples |
| 9[479]-4[483] | AAAAAGACTGGATTGAATCCCCCTCATAAATCAAGAAGCAAAGCCC | Core staples |
| 0[384]-6[367] | CCTGTTTGTGCCCGAAGTTCCAGTAAGCGTCATAGTGCCTTGAGCC | Core staples |
| 8[342]-15[349] | TCACCGGAACTTTGCCTTTAGCGGTAAATATCAATAGAA | Core staples |
| 15[413]-11[415] | GTAACACTGAGTTTCACAGACATGTCGTCTATGGGATTTTGCTGA | Biotin position |
| 4[430]-15[440] | AAGAACGGCTTTTTCGTTTATCAAGCATTCCGTCACCAGTACAAACT | Core staples |
| 7[462]-0[448] | ATCGTCATCCCCGAACTGACCATAAATTGTTCATTCCATATAACA | Core staples |
| 15[126]-14[147] | GCCGAAAGGAGCGGGCGCTAGGGCGCTGGCATGGTGGC | Core staples |
| 8[27]-15[34] | AGGTGGAGAAGGGGGATGTGCTGCCAGTTTGAACAAACG | Core staples |
| 11[385]-4[399] | TCAGGAGGTTCATAGGTGTACTCCTCAAGAGACTCCATT | Core staples |
| 4[167]-3[153] | CTGAAAGTTTGAATTAAAAATACCGAACGAACGGACAGAC | 3a position |
| 8[251]-7[272] | TCGCGCAGAATTTCATTTAATAAACACCGGAATCATATT | Core staples |
| 14[398]-8[378] | CTCAGAGATAGCCCACGTGCCGTCGAGAGGGTTCATA | Core staples |
| 14[461]-8[441] | TGCCCTGCATCAAGCGACCAGGCGCATAGGCTGGTGA | Core staples |
| 0[181]-4[179] | TTTTAGAACCCTCATAGAGGTGATCGCCATGGCTATTAGTCTAAC | Core staples |
| 11[406]-9[415] | GCGGAGTAAACACGATCTAAAGTTTGCCCTCACTT | Core staples |
| 15[98]-11[100] | TTACACTGGTGTGTTCTGCAGCCCAGCGCGTGCTGCGGCCAGCCG | Biotin position |
| 9[191]-0[182] | AGTATCCGAGGCCACCGAGCTGGCCATCGTCTGGTATTAAATAAAAAT | Core staples |
| 0[405]-8[406] | ATACATTTCCCCAGCGATTATACGCAGCGAAGCATAACCGCCGACAA | Core staples |
| 3[322]-6[322] | CAAGAAACAATCAAGCCCAATAATAAACCCACAGATAGCCG | Core staples |
| 12[398]-11[384] | AATAGTACCGCCACCCTCCGACTTGGTCACCAATGAAC | Core staples |
| 1a-4[41]-3[27] | TAAATTTGTTGTGTAGTGATGAAGGGTAAAGTTGAGCAGTAAGATCTACATA | DNA-PAINT docking at 1a position |
| 2a-4[104]-3[90] | GGTTTTTGATTGCCTTGCCCCAGCAGGCGAAAGGCGGGCAAAGATCTACATA | DNA-PAINT docking at 2a position |
| 3b-3[154]-11[153] | GATCTACATAAAAATATTCGTAAGACACGAGCAAATTAACCTAGAACGG | DNA-PAINT docking at 3b position |
| 4a-4[230]-3[216] | ATCCTTGGCTGAGATTTAACCTCCGGCTTAGGGATTAGATAAGATCTACATA | DNA-PAINT docking at 4a position |
| 5a-4[293]-3[279] | TCCAAATAATGAAAGACGGGAGAATTAACTGACTTTTAACAAGATCTACATA | DNA-PAINT docking at 5a position |
| 6a-4[356]-3[342] | TGATATTAGAATGGTTTGATGATACAGGAGTGTTCTCATTAAGATCTACATA | DNA-PAINT docking at 6a position |
| 7a-4[419]-3[405] | TCATGAGTAATGCCAACACTCATCTTTGACCCCCGGGTAAAAGATCTACATA | DNA-PAINT docking at 7a position |
| 8a-4[482]-3[468] | GAAAGACCAAAGCGTAATTGCTCCTTTTGATATCTAATTCAAGATCTACATA | DNA-PAINT docking at 8a position |
| Docking-mid-10nt-3Aspacer | GGATAAGGAAGGTGGGCAGGAAAGATCTACATAAAACGTTTGTTAGGGATTGGGTG | DNA-PAINT docking for tension study |
| biotin-4T-15[35]-11[37] | /5Biosg/TTTTGCGGATTGACCGTAATAACCGTTCGGCCTGGCACCGCTTCTGGCC | Biotinylated strands |
| biotin-4T-15[98]-11[100] | /5Biosg/TTTTTTACACTGGTGTGTTCTGCAGCCCAGCGCGTGCTGCGGCCAGCCG | Biotinylated strands |
| biotin-4T-15[161]-11[163] | /5Biosg/TTTTGGTCACGCTGCGCGTAGGGCGCGTGCTTTAGGAGGCCGATTAAGA | Biotinylated strands |
| biotin-4T-15[224]-11[226] | /5Biosg/TTTTTTATCATTTTGCGGAATATTCCATCCTGAAGGGTTAGAACCTCAG | Biotinylated strands |
| biotin-4T-15[287]-11[289] | /5Biosg/TTTTTACCGACAAAAGGTATGTTCAGGATAAGTAATTTACGAGCATTTC | Biotinylated strands |
| biotin-4T-15[350]-11[352] | /5Biosg/TTTTAATTCATATGGTTTATTGAGGGTAAAGGTTTGGGAATTAGAGCAT | Biotinylated strands |
| biotin-4T-15[413]-11[415] | /5Biosg/TTTTGTAACACTGAGTTTCACAGACATGTCGTCTATGGGATTTTGCTGA | Biotinylated strands |
| biotin-4T-15[476]-11[478] | /5Biosg/TTTTCAACTTTAATCATTGTATACCATAAAACGAAAGATTCATCAGAAT | Biotinylated strands |

| | | |
|---|---|---|
| 200nt ssDNA spring 2-3[406]-11[405] | CCTGCCCACCTTCCTTATCCTTTTTTTTTTTTTTTTTTTTTTTTTTTTTT TTTTTTTTTTTTTTTTTTTTTTTTTTTTTTTTTTTTTTTTTTTTTTTTTT TTTTTTTTTTTTTTTTTTTTTTTTTTTTTTTTTTTTTTTTTTTTTTTTTT TTTTTTTTTTTTTAATACGGAAGTTAGCATCCAAAAAAAAGGAAGTTTCA | 200nt ssDNA spring b-staple at 7b position |
| 117nt ssDNA spring 2-3[406]-11[405] | CCTGCCCACCTTCCTTATCCTTTTTTTTTTTTTTTTTTTTTTTTTTTTTT TTTTTTTTTTTTTTTTTTTTTTTTTTTTTTAATACGGAAGTTAGCATCCA AAAAAAAGGAAGTTTCA | 117nt ssDNA spring b-staple at 7b position |
| 4[167]-3[153]-200nt ssDNA spring 1 | CTGAAAGTTTGAATTAAAAATACCGAACGAACGGACAGACTCTCTCCTCC TTCTCCTCTCTCTCCTTCCTTCCTCCCTTACCATCCCTCCGCTCTTTCTC TCCCTTTCCCTCCTCCTCTCTTTTTTTTTTTTTTTTTTCCCTCCTTTCCT CTTCACACTCTCCCTTCACACTCCCACTCCCACCCAATCCCTAACAAACG | Staple at 3a position-200nt ssDNA spring a |
| 4[167]-3[153]-120nt ssDNA spring 1 | CTGAAAGTTTGAATTAAAAATACCGAACGAACGGACAGACTTTTTTTTTT TTTTTTTTCCCTCCTTTCCTCTTCACACTCTCCCTTCACACTCCCACTCC CACCCAATCCCTAACAAACG | Staple at 3a position-120nt ssDNA spring a |
| 4[167]-3[153]-100nt ssDNA spring 1 | CTGAAAGTTTGAATTAAAAATACCGAACGAACGGACAGACCTCCTTTCCT CTTCACACTCTCCCTTCACACTCCCACTCCCACCCAATCCCTAACAAACG | Staple at 3a position-100nt ssDNA spring a |
| 4[41]-3[27]-200 nt ssDNA spring 3 | TAAATTTGTTGTGTAGTGATGAAGGGTAAAGTTGAGCAGTTTCCTTCCCT TTATCTCCCTTTTACCTCCTCCACCCGCCCACCTTTCTCCCATATTCCAC CTCTCTCCCTTCCTCTCCTTTTTTTTTTTTTTTTTTTCTCCTCTTCCTCT TTCCTCTTCCCTCTTTCTCTCCTCCTCTTCCCTTTTCTTTCTCACATTAT | Staple at 1a position-200 nt ssDNA spring c |
| 117 nt ssDNA spring 4-3[280]-11[279] | TTCCTTTCATCCTTACTCTTTTTTTTTTTTTTTTTTTTTTTTTTTTTTTT TTTTTTTTTTTTTTTTTTTTTTTTTTTTTTGTCAAAAAGAAATTTCCACG TTTTTATTTTAAATCGG | 117 nt ssDNA spring d-Staple at 5b position |

**Table S1** **MAESTRO structure, DNA-PAINT, and ssDNA entropic spring strands.** The scaffold is p8064. Core staples are all the strands that form the structure. Biotinylated staples were modified with biotin to immobilize MAESTRO on coverslip surfaces through a BSA-biotin-streptavidin-biotin-DNA origami arrangement. DNA-PAINT docking strands were used to acquire super-resolution DNA-PAINT images of MAESTRO (Fig. 1e, right panel). ssDNA entropic spring strands are selected to form MAESTRO with specific tension force configurations. Positions 1 (a,b) to 8 (a,b) correspond to specific sites on the inner sidewall of MAESTRO based on (Fig. S1).

| Name | Sequence | Tension Configuration |
|---|---|---|
| x-extension to spring 2<br>r<br>Cy5-b<br>Cy3-h-extension to spring 1 | GGATAAGGAAGGTGGGCAGGTTTCCCAGTTGAGCGCTTGCTAGGG<br>CCCACCGCTCGGCTCAACTGGG<br>/Cy5/CCCTAGCAAGCCGCTGCTACGG<br>/Cy3/CCGTAGCAGCGCGAGCGGTGGGTTTCGTTTGTTAGGGATTGGGTG | Single-axial y-axis |
| Cy3-h<br>Cy5-b<br>r<br>x-extension to spring 2 and spring 1 | /Cy3/CCGTAGCAGCGCGAGCGGTGGG<br>/Cy5/CCCTAGCAAGCCGCTGCTACGG<br>CCCACCGCTCGGCTCAACTGGG<br>GGATAAGGAAGGTGGGCAGGTTTCCCAGTTGAGCGCTTGCTAGGG<br>TTTCGTTTGTTAGGGATTGGGTG | Single-axial x-axis |
| x-extension to spring 2 and spring 3<br>r<br>Cy5-b<br>Cy3-h-extension to spring 1 | GGATAAGGAAGGTGGGCAGGTTTCCCAGTTGAGCGCTTGCTAGGGT<br>TTATAATGTGAGAAAGAAAAGG<br>CCCACCGCTCGGCTCAACTGGG<br>/Cy5/CCCTAGCAAGCCGCTGCTACGG<br>/Cy3/CCGTAGCAGCGCGAGCGGTGGGTTTCGTTTGTTAGGGATTGGGTG | Three-way tension 1 |
| Cy3-h-extension to spring 3<br>Cy5-b<br>r<br>x-extension to spring 2 and spring 1 | /Cy3/CCGTAGCAGCGCGAGCGGTGGGTTTATAATGTGAGAAAGAAAAGG<br>/Cy5/CCCTAGCAAGCCGCTGCTACGG<br>CCCACCGCTCGGCTCAACTGGG<br>GGATAAGGAAGGTGGGCAGGTTTCCCAGTTGAGCGCTTGCTAGGGTT<br>TCGTTTGTTAGGGATTGGGTG | Three-way tension 2 |
| Cy3-h<br>r-extension to spring 4<br>x-extension to spring 2 and spring 3<br>Cy5-b-extension to spring 1 | /Cy3/CCGTAGCAGCGCGAGCGGTGGG<br>AAGAGTAAGGATGAAAGGAATTTCCCACCGCTCGGCTCAACTGGG<br>GGATAAGGAAGGTGGGCAGGTTTCCCAGTTGAGCGCTTGCTA<br>GGGTTTATAATGTGAGAAAGAAAAGG<br>/Cy5/CCCTAGCAAGCCGCTGCTACGGTTTCGTTTGTTAGGGATTGGGTG | Four-way tension |

**Table S2** **HJ strands for various tension forces configuration.** Set of DNA strands to form HJ for specific tension force configuration. Each set consists of 4 strands in equimolar final concentration of 200 nM in a final 1× TAE 12.5 mM $Mg^{2+}$.

| **Strands** | **Final Concentration** |
| --- | --- |
| p8064 scaffold | 20 nM |
| Core staples (staples at biotin, 1a, 2a, 3b, 4a, 5a, 6a, 7a, and 8a positions should be excluded) | 200 nM/strand |
| Biotinylated staples | 200 nM/strand |
| DNA-PAINT dockings | 1000 nM/strand |
| 1× TAE 12.5 mM $MgCl_2$ buffer | 1× |

**Table S3 MAESTRO mixing concentrations for DNA-PAINT assay.**

| Tension | Strands | Final Concentration |
|---|---|---|
| 0 pN | p8064 scaffold | 20 nM |
| | Core staples (excluding biotin and 3a positions) | 200 nM/strand |
| | Biotinylated staples | 200 nM/strand |
| | 4[167]-3[153]-200nt ssDNA spring 1 | 500 nM/strand |
| | 1× TAE 12.5 mM $MgCl_2$ buffer | 1× |
| 2 pN | p8064 scaffold | 20 nM |
| | Core staples (excluding biotin, 3a, and 7b positions) | 200 nM/strand |
| | Biotinylated staples | 200 nM/strand |
| | 4[167]-3[153]-200nt ssDNA spring 1 | 500 nM/strand |
| | 200nt ssDNA spring 2-3[406]-11[405] | 500 nM/strand |
| | 1× TAE 12.5 mM $MgCl_2$ buffer | 1× |
| 3 pN | p8064 scaffold | 20 nM |
| | Core staples (excluding biotin, 3a, and 7b positions) | 200 nM/strand |
| | Biotinylated staples | 200 nM/strand |
| | 4[167]-3[153]-200nt ssDNA spring 1 | 500 nM/strand |
| | 117nt ssDNA spring 2-3[406]-11[405] | 500 nM/strand |
| | 1× TAE 12.5 mM $MgCl_2$ buffer | 1× |
| 6 pN | p8064 scaffold | 20 nM |
| | Core staples (excluding biotin, 3a, and 7b positions) | 200 nM/strand |
| | Biotinylated staples | 200 nM/strand |
| | 4[167]-3[153]-120nt ssDNA spring 1 | 500 nM/strand |
| | 117nt ssDNA spring 2-3[406]-11[405] | 500 nM/strand |
| | 1× TAE 12.5 mM $MgCl_2$ buffer | 1× |
| 9 pN | p8064 scaffold | 20 nM |
| | Core staples (excluding biotin, 3a, and 7b positions) | 200 nM/strand |
| | Biotinylated staples | 200 nM/strand |
| | 4[167]-3[153]-100nt ssDNA spring 1 | 500 nM/strand |
| | 117nt ssDNA spring 2-3[406]-11[405] | 500 nM/strand |
| | 1× TAE 12.5 mM $MgCl_2$ buffer | 1× |
| Three-way | p8064 scaffold | 20 nM |
| | Core staples (excluding biotin, 1a, 3a, and 7b positions) | 200 nM/strand |
| | Biotinylated staples | 200 nM/strand |
| | 4[167]-3[153]-200nt ssDNA spring 1 | 500 nM/strand |
| | 117nt ssDNA spring 2-3[406]-11[405] | 500 nM/strand |
| | 4[41]-3[27]-200nt ssDNA spring 3 | 500 nM/strand |
| | 1× TAE 12.5 mM $MgCl_2$ buffer | 1× |
| Four-way | p8064 scaffold | 20 nM |
| | Core staples (excluding biotin, 1a, 3a, 5b, and 7b positions) | 200 nM/strand |
| | Biotinylated staples | 200 nM/strand |
| | 4[167]-3[153]-200nt ssDNA spring 1 | 500 nM/strand |
| | 117nt ssDNA spring 2-3[406]-11[405] | 500 nM/strand |
| | 4[41]-3[27]-200nt ssDNA spring 3 | 500 nM/strand |
| | 117nt ssDNA spring 4-3[280]-11[279] | 500 nM/strand |
| | 1× TAE 12.5 mM $MgCl_2$ buffer | 1× |

**Table S4 MAESTRO mixing concentrations across different tension conditions.**

| **Parameters** | **Values** |
|---|---|
| $k_{\mathrm{off}}(0)$ | 0.07 $s^{-1}$ |
| $\Delta x_{\mathrm{off}}$ | 0.32 nm |
| $k_{\mathrm{on}}(0)$ | 2.63 $\mu M^{-1} s^{-1}$ |
| $\frac{1}{a}$ | -0.115 $pN^{-1}nm$ |
| $b$ | 0.84 nm |

**Table S5  Fitted values of kinetics parameters for short oligos under tension.**

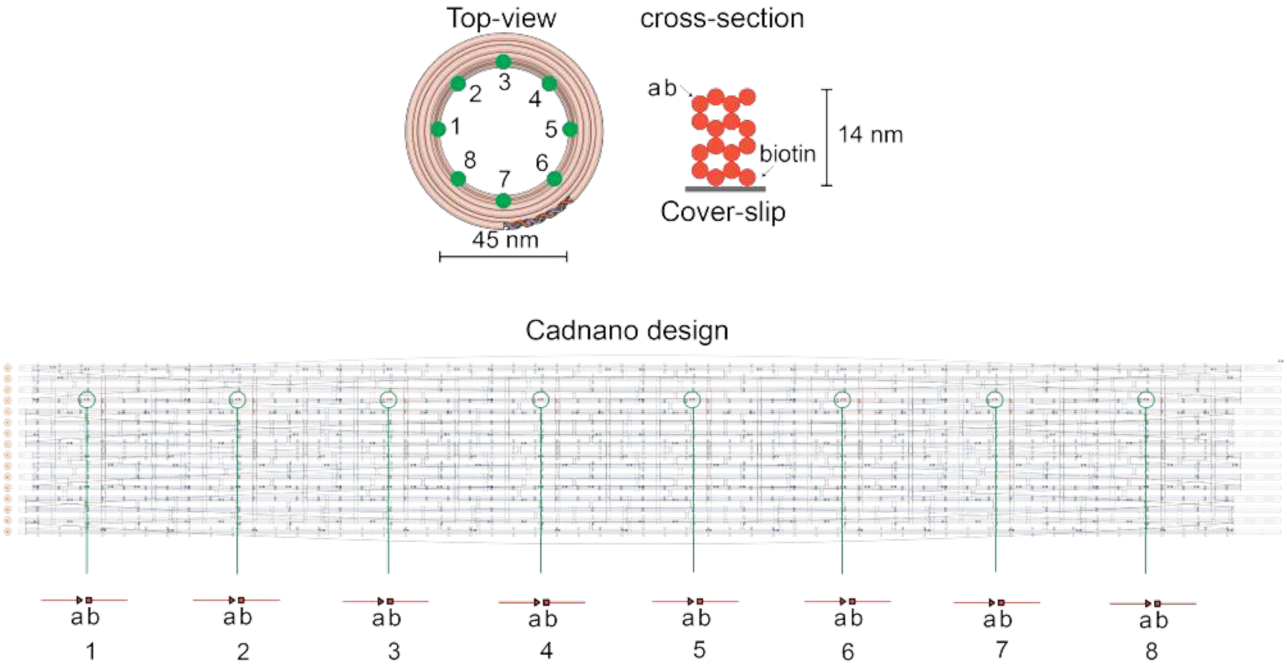

**Fig. S1 MAESTRO schematics (top) and cadnano design (bottom) showing the position of points 1a,b to 8a,b corresponding to some strands in Table S1**

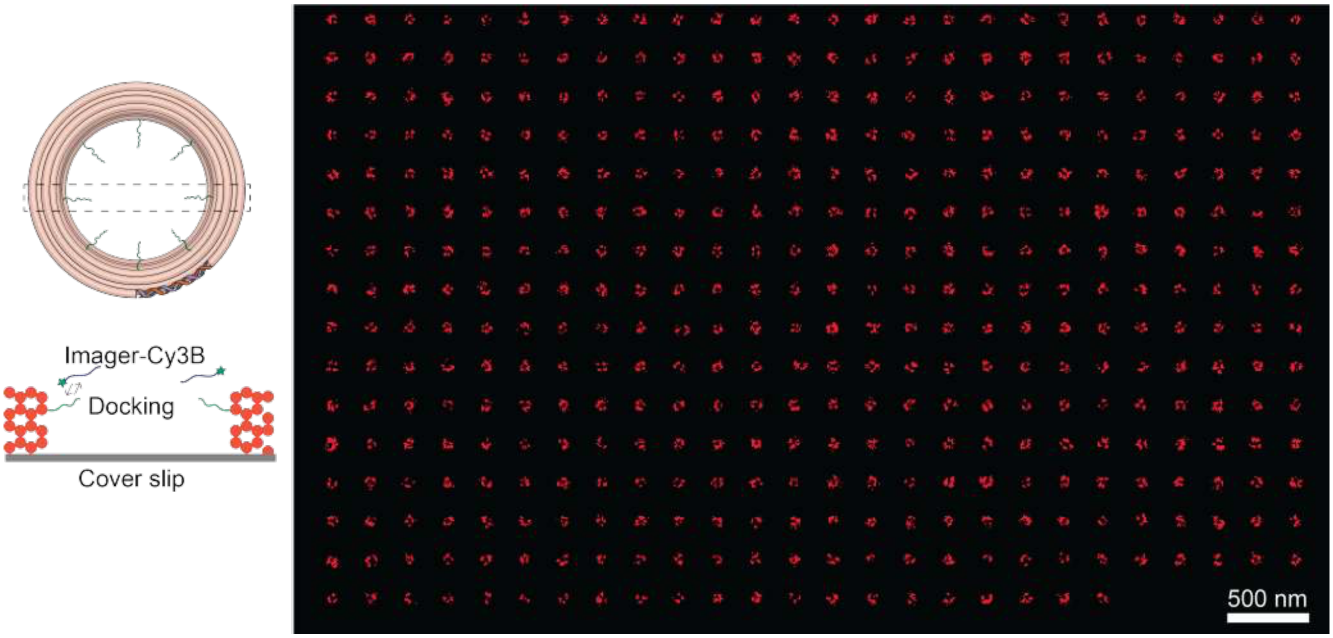

**Fig. S2** **MAESTRO DNA-PAINT schematics illustrating the arrangement of docking strands and the organization of all selected molecules in an array.**

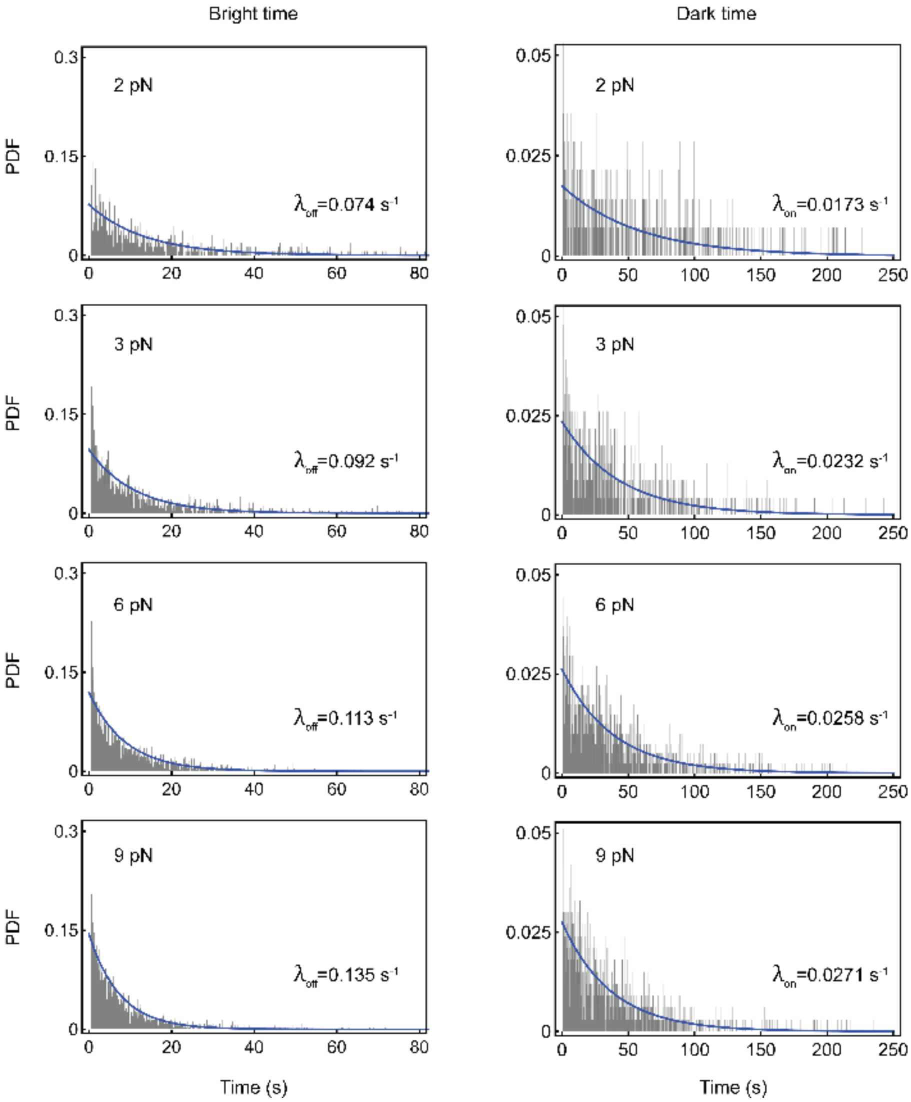

**Fig. S3 Histogram plots of bright (left) and dark times (right) of short oligos under various tension forces, using a bin width of 100 ms—equal to the camera exposure time—along with exponential fit curves (blue).**

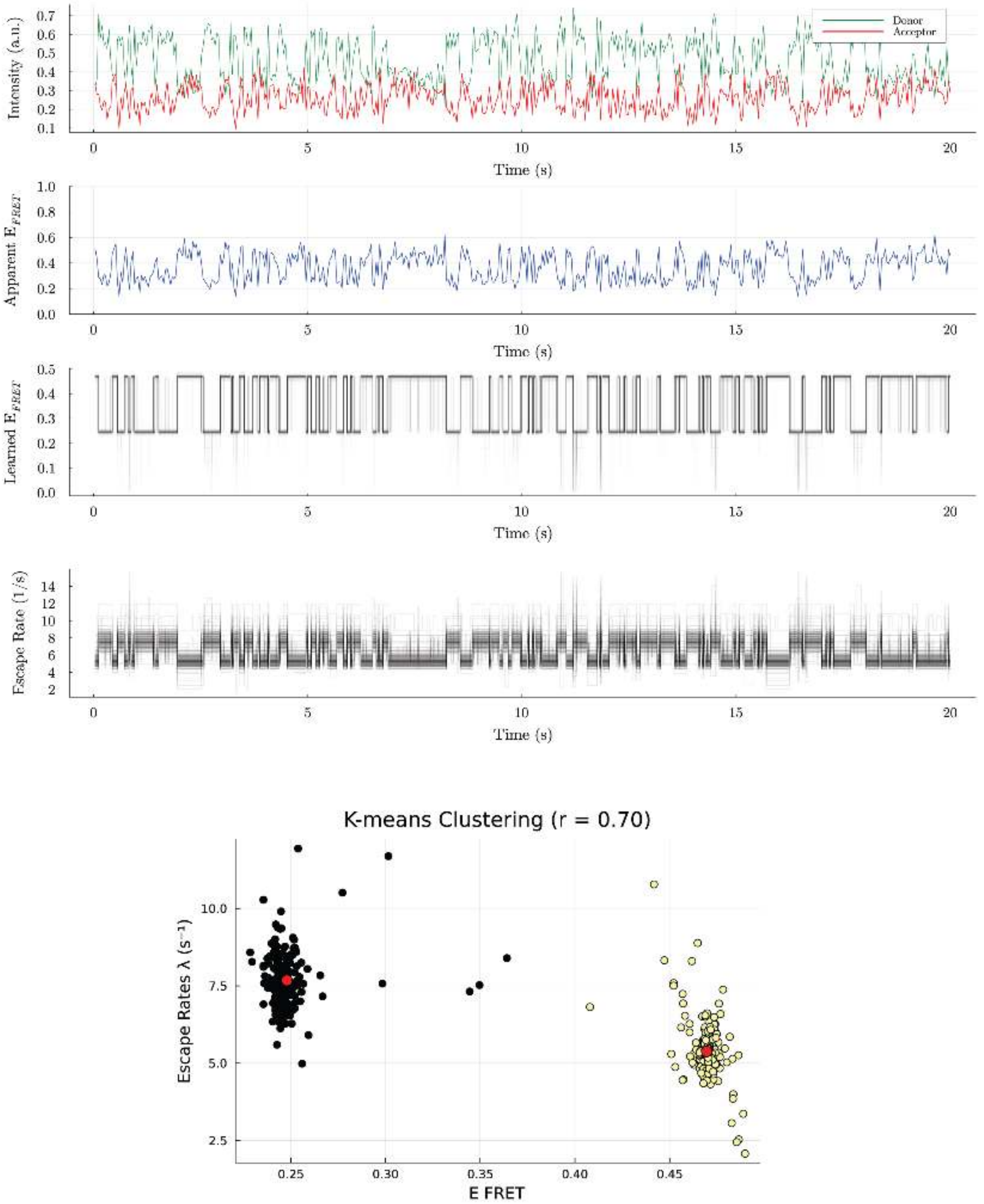

**Fig. S4** **An example of a smFRET trajectory showing donor's and acceptor's emissions at 0 pN (1st row), the apparent FRET efficiency (2nd row), the learned FRET efficiency (3rd row), the learned escape rates (4th row), and the escape rates k-means clustering performed on MCMC samples generated by BNP-FRET to extract the mean escape rate $\lambda$ values of Iso-I and Iso-II used to calculate the ratio $r = \lambda_{\text{Iso-I}}/\lambda_{\text{Iso-II}}$ where red dots show the mean values for each cluster colored in black and yellow (5th row)**

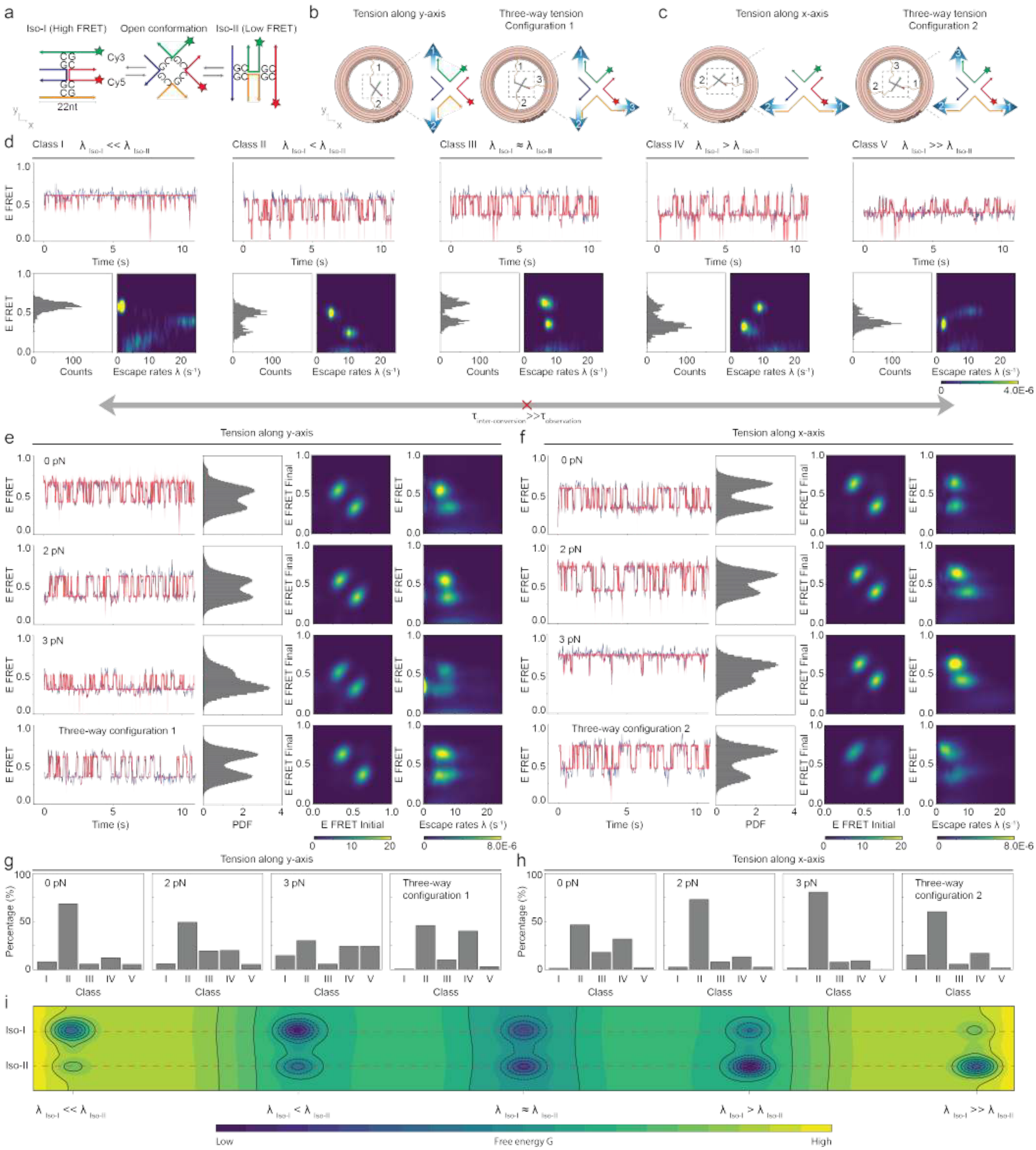
a
Iso-I (High FRET)
Open conformation
Iso-II (Low FRET)
Cy3
Cy5
22nt
b
Tension along y-axis
Three-way tension Configuration 1
c
Tension along x-axis
Three-way tension Configuration 2
d
Class I λ Iso-I << λ Iso-II
Class II λ Iso-I < λ Iso-II
Class III λ Iso-I ≈ λ Iso-II
Class IV λ Iso-I > λ Iso-II
Class V λ Iso-I >> λ Iso-II
E FRET
Time (s)
Counts
Escape rates λ (s⁻¹)
4.0E-6
τ inter-conversion >> τ observation
e
Tension along y-axis
0 pN
2 pN
3 pN
Three-way configuration 1
PDF
E FRET Initial
E FRET Final
8.0E-6
f
Tension along x-axis
Three-way configuration 2
g
Tension along y-axis
Percentage (%)
Class
h
Tension along x-axis
i
Iso-I
Iso-II
λ Iso-I << λ Iso-II
λ Iso-I < λ Iso-II
λ Iso-I ≈ λ Iso-II
λ Iso-I > λ Iso-II
λ Iso-I >> λ Iso-II
Low
Free energy G
High

**Fig. S5** **(Previous page) HJs under single-axial and 3-way tension forces configuration.** For comparison with main figures, panels d and portions of e,f repeat data from Figs. 2 and 3. (**a**) Schematic of the J7 Holliday junction (HJ) show Iso-I, Iso-II, and open conformations, with a FRET pair (Cy3 and Cy5) attached to the ends of two arms. (**b, c**) Schematics illustrating the HJ connection to MAESTRO for different tension configurations: single-axial tension along the $y$ and $x$-axes and two configurations of 3-way tension forces. Labels 1, 2, and 3 represent the distinct ssDNA entropic springs. (**d**) Five classes of smFRET trajectories grouped based on the relative escape rates of conformations. Top panels show representative smFRET (blue) and HMM (red) trajectories. Bottom panels show the corresponding smFRET histograms and escape rate heatmap. The absence of interconversion between classes suggests that $\tau_{\text{inter-conversion}} \gg \tau_{\text{observation}}$. (**e, f**) smFRET(blue) and BNP-FRET trajectories(red), ensemble smFRET histograms, transition density plots, and escape rate heatmaps compiled from $n$ >150 molecules under different tension force configurations: (e) $y$-axis and 3-way tension forces configuration 1, (f) $x$-axis and three-tension force configuration 2. (**g, h**) Histograms showing the percentage of HJ molecules classified into each of the five dynamic classes under different tension conditions: (g) $y$-axis and 3-way tension forces configuration 1, (h) $x$-axis and 3-way tension forces configuration 2. (**i**) Qualitative visualization of the inferred energy landscapes corresponding to five dynamic classes.

Following the work by Hyeon et al.[42], where they are able to classify Holliday junction smFRET traces based on kinetics, here we use a similar empirical approach to categorize the kinetic behavior. We found that our data can be categorized into five distinct classes. Intuitively, these five classes can be distinguished based on the ratio of the mean escape rates, $r = \lambda_{\text{Iso-I}}/\lambda_{\text{Iso-II}}$. We identified the mean escape rate values using k-means clustering of MCMC samples for each smFRET trace(see Fig. S5). That is, we have extremal classes I ($r < 0.2$) and V ($r > 5.0$), where the Holliday junction gets stuck in either of the two conformations, Iso-I (high FRET) or Iso-II (low FRET); class III ($0.9 < r < 1.1$), where the Holliday junction switches between the two conformations at approximately the same rate; and classes II and IV, where the Holliday junction escape rates take intermediate values between the previously defined classes, as shown in Fig. S5(d) for a tension of 0 pN. These classes showed clear differences in their smFRET histograms and escape rate profiles (Fig. S5(d)). This variability likely arises from the rugged energy landscape of the junction, as previously proposed by Hyeon et al.[42], where complex interactions between $Mg^{2+}$ ions and the junction structure create several kinetic traps. Consistent with their findings, our trajectories show no interconversion at 0 pN between classes within a 50 sec observation window.

Increasing the tension along the $y$-axis using springs 1 and 2 shifts the population toward the Iso-II state, thereby reducing the relative occupancy of Iso-I. This tension-induced population shift is clearly visible in the ensemble FRET histograms at 0, 2, and 3 pN (Fig. S5(e), column 2 panels). However, introducing spring 3 to apply 3-way tension forces balances the conformation populations, resulting in nearly equal occupancy of Iso-I and Iso-II (Fig. S5e, column 2, row 4).

Across all conditions, most molecules transitioned between two well-defined FRET states with efficiencies of $E_{\text{FRET}} \sim 0.3$ (Iso-II) and $E_{\text{FRET}} \sim 0.6$ (Iso-I), a hallmark of the canonical HJ behavior.[46] This dynamics is captured as two dense hotspots in the transition density heat maps (Fig. S5e, column 3). As the tension along the $y$ axis increases, the escape rate distribution shows that $\lambda_{\text{Iso-II}}$ changes to slower values, while $\lambda_{\text{Iso-I}}$ changes to faster rates (Fig. S5(e)), column 4, compare row 1 to row 3). Under 3-way tension, however, the escape rates of both conformations become more balanced and span a wider range, in contrast to the asymmetric distribution observed at 0 pN (Fig. S5(e), column 4, compare rows 1 and 4).

We observe the opposite trend when tension was applied along the $x$-axis (Fig. S5(f)). That is, increasing the force from 0 to 3 pN favors the Iso-I state over Iso-II, as shown in the ensemble FRET histograms (Fig. S5(f), column 2, rows 1–3). Transitions predominantly occur between the two main conformations (Fig. S5f, column 3), but the escape rate behavior reverses: $\lambda_{\text{Iso-I}}$ slows down while $\lambda_{\text{Iso-II}}$ increases with tension (Fig. S5(f), column 4, compare row 1 with row 3).

We also note that in the case of 3pN tension along $y$-axis, the HJ gets stuck in Iso-II, as indicated by the bright spot near the 0 $s^{-1}$ escape rate at low FRET efficiency ($E_{\text{FRET}}$) in Fig. S5(e), row 3, column 4, respectively. We also observed this behavior when the tension was along $x$-axis, as observed in the trace shown in Fig. S5(f), row 3, column 1, albeit with far fewer molecules, as indicated by the lack of a similar bright spot at a high $E_{\mathrm{FRET}}$ in the corresponding ensemble escape rate heatmap.

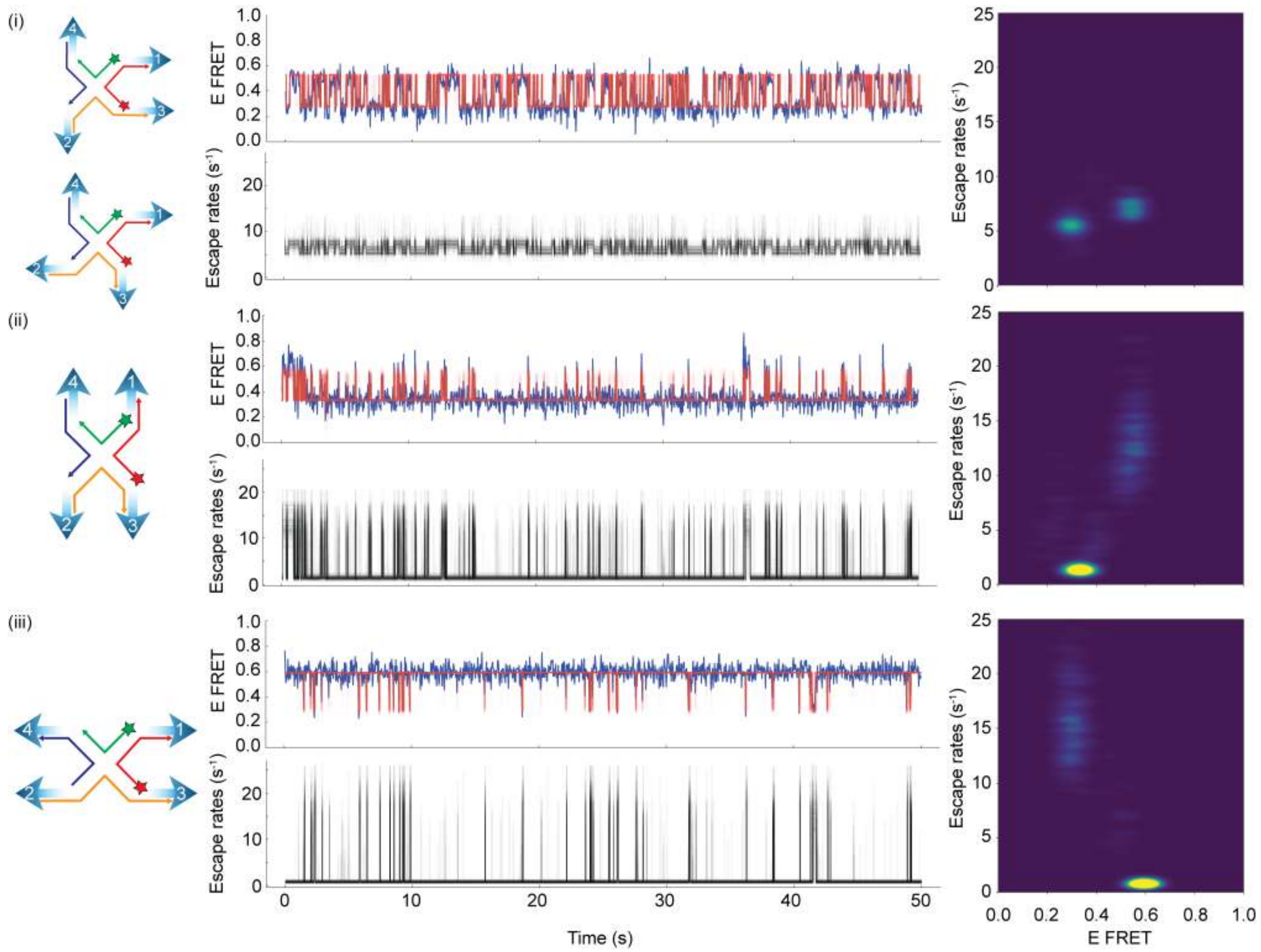

**Fig. S6 Additional examples of smFRET traces from the four-way tension force configuration, illustrating the proposed direction of tension forces on the molecules, along with their corresponding trace and escape rate heatmaps.** These examples include traces that switch between Iso-I and Iso-II ($i$), remain stuck at Iso-II ($ii$), and remain stuck at Iso-I ($iii$).

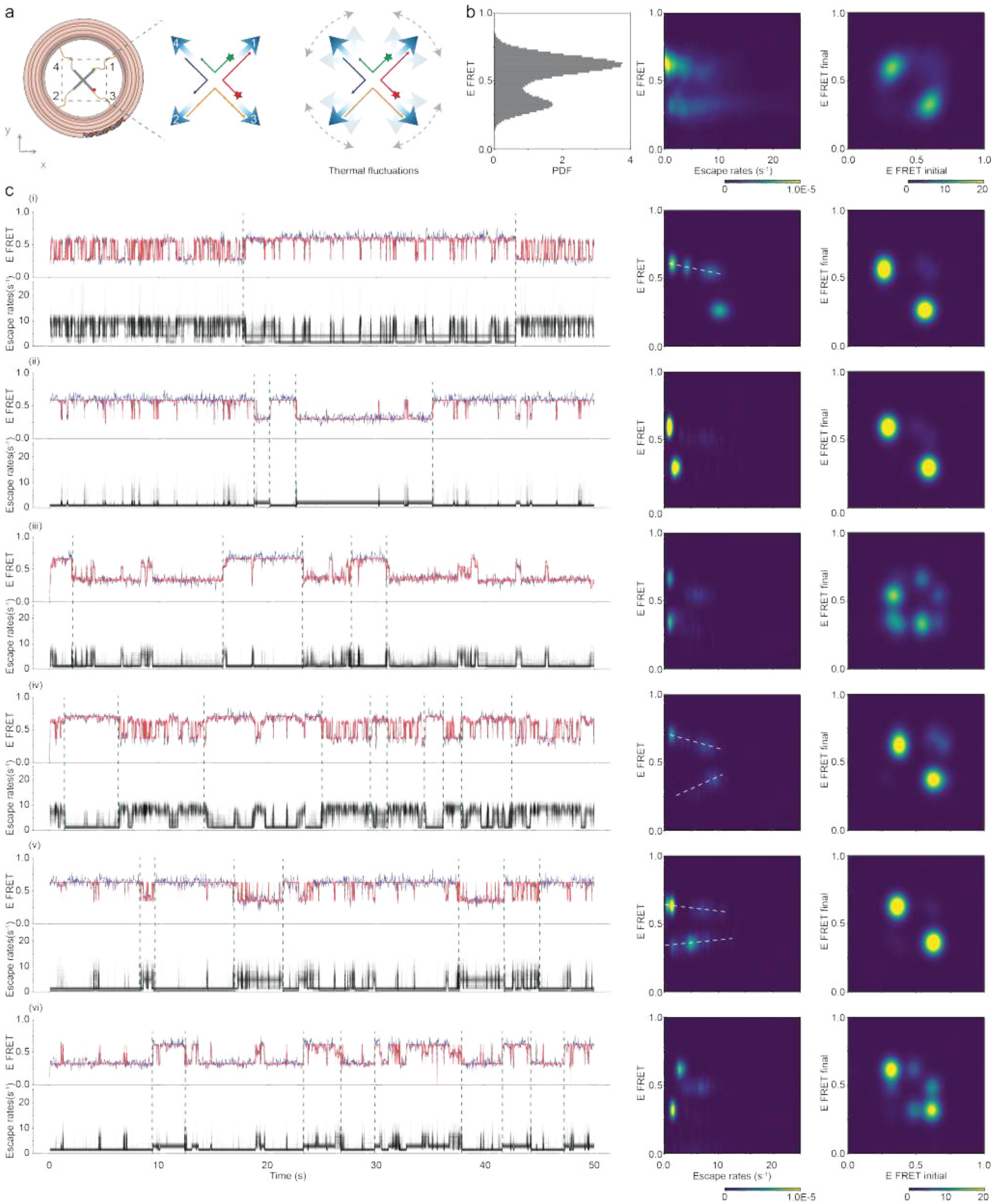
a
b
c
Thermal fluctuations
E FRET
PDF
Escape rates (s⁻¹)
E FRET final
E FRET initial
1.0E-5
(i)
(ii)
(iii)
(iv)
(v)
(vi)
Time (s)

**Fig. S7** **(Previous page) Interconversion between smFRET dynamic classes in J7 HJs under multi-axial tension from four-way tension forces.** For comparison with main figures, traces 1 and portions of 4, and 6 are repeat data from Fig. 3. (**a**) Schematic of the four-way tension configuration on a Holliday junction (HJ), showing both the structural setup (left and middle panels) and the possible directions in which each entropic spring can stochastically pull the junction owing to thermal fluctuations (right panel). (**b**) Ensemble smFRET histogram, escape rate distributions, and transition density heatmaps ($n$=139 HJ). (**c**) Six representative smFRET trajectories (blue) overlaid with BNP-FRET trajectories (red) (top-left panels), accompanied by their corresponding escape rate trajectories (bottom-left panels), extracted escape rates (middle panels), and transition density heatmaps (right panels). Green dashed lines indicate visually identified interconversion points between smFRET dynamic classes based on changes in escape rate behavior. White dashed lines in some heatmaps highlight subtle shifts in $E_{FRET}$ between distinct Iso-I and Iso-II conformations.

In the first example (Fig. S7(c)i), the molecule transitions between fast-switching states (class II, III, and IV) and class I, where the junction becomes mostly trapped in the Iso-I conformation. The corresponding escape rate trajectory and heatmap revealed multiple degenerate states with escape rates of $\sim 1$ s$^{-1}$, $\sim 4$ s$^{-1}$, and $\sim 9$ s$^{-1}$, all near the same FRET efficiency ($E_{FRET}\sim0.6$). However, states with faster escape rates tended to exhibit slightly lower $E_{FRET}$. These Iso-I states transition with a single Iso-II state ($E \sim 0.25$) characterized by a $\sim 10$ s$^{-1}$ escape rate. The transition density heatmap confirmed that major transitions occurred between these two main states.

The second trace (Fig. S7(c)ii) shows four interconversions between classes I and V within the 50 sec observation window. In the corresponding escape rate trajectory and heatmap, higher escape rates were observed during the transitions between Iso-I and Iso-II. The Iso-I state ($E \sim 0.6$) exhibits a strong escape rate density at $\lambda_{\text{Iso-I}} \sim 0.5$ s$^{-1}$, whereas the Iso-II state ($E \sim 0.3$) shows a prominent density at $\lambda_{\text{Iso-II}} \sim 2$ s$^{-1}$. The transition density heatmap confirmed that dominant transitions occurred between these two conformations.

In the third trace (Fig. S7(c)iii), five interconversions between class I ($E_{\text{FRET}} \sim 0.6$, $\lambda_{\text{Iso-I}} \sim 0.5$ s$^{-1}$) and class V ($E_{\text{FRET}} \sim 0.3$, $\lambda_{\text{Iso-II}} \sim 0.5$ s$^{-1}$) are clearly visible and separated by green dashed lines. The presence of intermediate transition states ($0.3 < E_{\text{FRET}} < 0.6$) is supported by multiple hotspots in the transition density heatmap, and these states exhibit faster escape rates in the range $5$ s$^{-1} < \lambda_{\text{transition}} < 10$ s$^{-1}$.

The fourth trace (Fig. S7(c)iv) shows ten interconversions between class I and faster-switching conformation classes (II and III). The Iso-I state in these fast-switching classes has a slightly lower FRET efficiency ($E_{FRET}\sim0.6$) than the class I Iso-I conformation ($E_{FRET}\sim0.7$), as observed in the escape rate heatmap. Transitions between the two Iso-I states are visible as faint hotspots in the transition density heatmap, along with the dominant Iso-I and Iso-II transitions.

A more complex dynamic behavior is seen in the fifth trace (Fig. S7(c)v), where the molecule interconverts between class I, class V, and class II. Multiple clusters in the escape rate plot for both Iso-I and Iso-II suggest dynamic heterogeneity under four-way tension forces. Additionally, Iso-II states with slower escape rates tend to exhibit lower FRET efficiencies, suggesting the presence of distinct energy levels. This observation is consistent with the behavior previously noted for Iso-I in the first and fourth trace.

In the sixth trace (Fig. S7(c)vi), a distinct transition state appears at $E_{\text{FRET}} \sim 0.5$, visible in the smFRET trajectory, escape rate heatmap, and transition density heatmap. This state exists alongside clear interconversions between classes I and V.

Together, these examples demonstrate that four-way tension forces enable J7 HJs to explore a broader conformational energy landscape, facilitating multiple interconversions between dynamic classes within the 50 sec observation window. This is likely due to the simultaneous pulling of all four arms by entropic springs, allowing access to a wide range of force directions and conformational pathways. Such interconversions were not observed under tension-free and single-axial tension conditions.

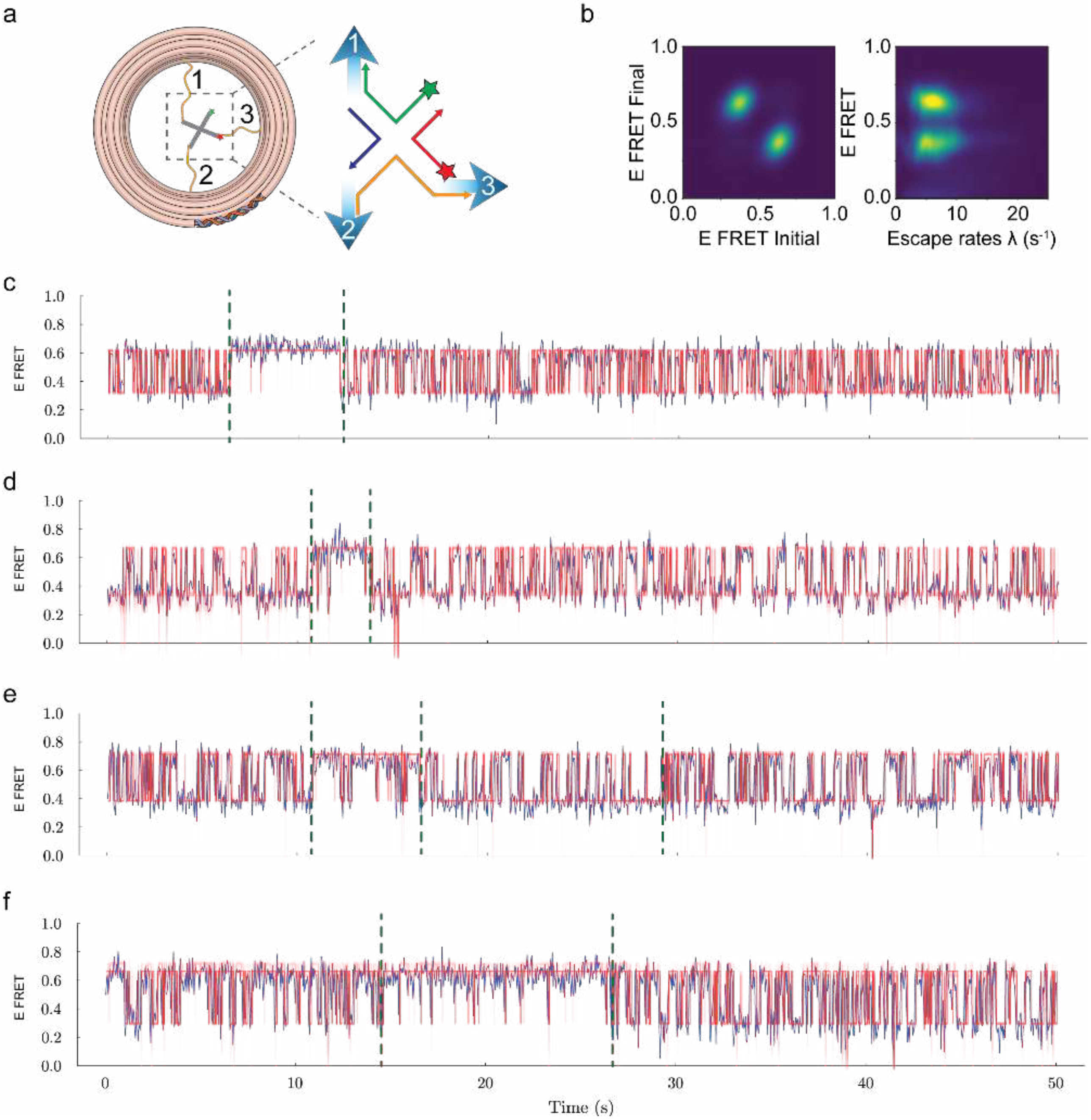

**Fig. S8 Kinetic class interconversion under 3-way tension.** (**a**) Schematic of 3-way tension on a Holliday junction. (**b**) Escape rates and transition density heatmaps. (**c-f**) Examples of FRET dynamics showing kinetic class interconversion. Green dashed lines indicate visually identified interconversion points.

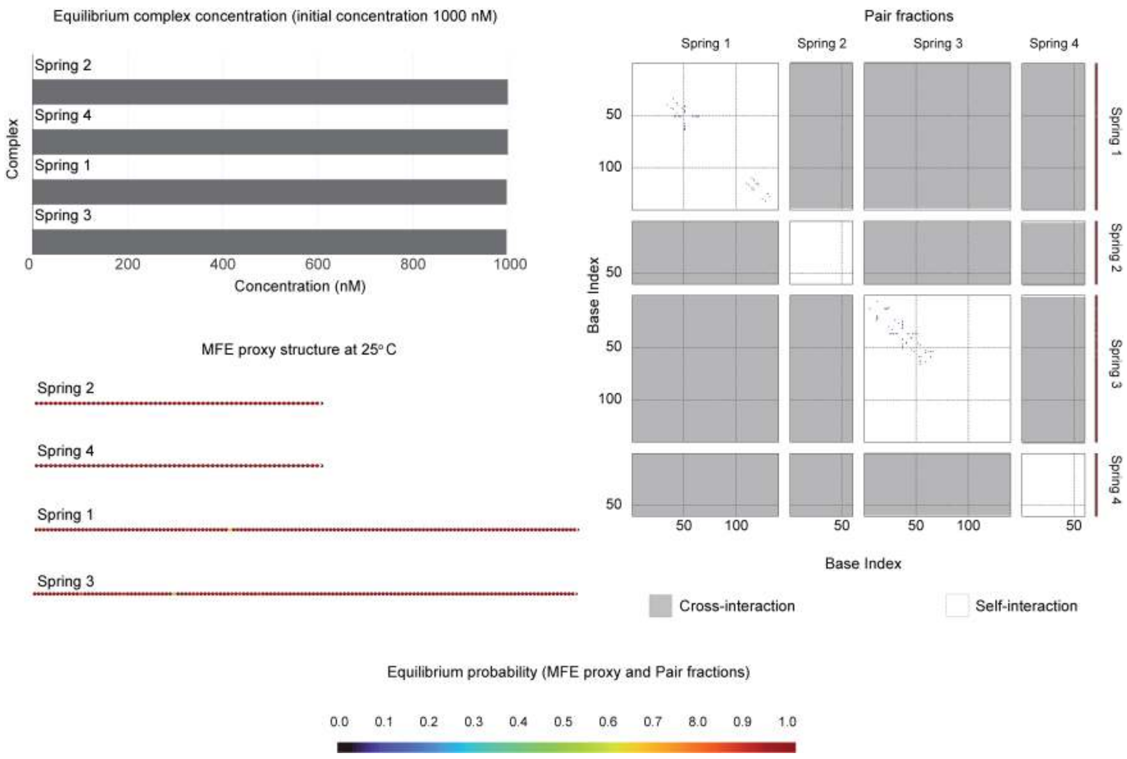

**Fig. S9 NUPACK analysis of the ssDNA springs at 25 °C showing the equilibrium complex concentrations, pair fractions, and minimum free energy (MFE) proxy structures.** The results indicate that all ssDNA springs exhibit negligible self- and cross-interactions, as evidenced by the MFE proxy structures and pair interaction profiles.
